# Comprehensive analysis of computational approaches in plant transcription factors binding regions discovery

**DOI:** 10.1101/2023.11.07.566153

**Authors:** Jyoti, Ritu, Sagar Gupta, Ravi Shankar

**Affiliations:** Studio of Computational Biology & Bioinformatics, The Himalayan Centre for High-throughput Computational Biology, (HiCHiCoB, A BIC supported by DBT, India), Biotechnology Division CSIR-Institute of Himalayan Bioresource Technology (CSIR-IHBT), Palampur (HP), 176061, India; Academy of Scientific and Innovative Research (AcSIR), Ghaziabad, Uttar Pradesh- 201002

**Keywords:** Transcription factors, Transcription factor binding regions, Machine learning, Deep learning, TF-DNA binding specificity

## Abstract

Transcription factors (TFs) are regulatory proteins which bind to a specific DNA region known as the transcription factor binding regions (TFBRs) to regulate the rate of transcription process. The identification of TFBRs has been made possible by a number of experimental and computational techniques established during the past few years. The process of TFBR identification involves peak identification in the binding data, followed by the identification of motif characteristics. Using the same binding data attempts have been made to raise computational models to identify such binding regions which could save time and resources spent for binding experiments. These computational approaches depend a lot on what way they learn and how. These existing computational approaches are skewed heavily around human TFBRs discovery, while plants have drastically different genomic setup for regulation which these approaches have grossly ignored. Here, we provide a comprehensive study of the current state of the matters in plant specific TF discovery algorithms. While doing so, we encountered several software tools’ issues rendering the tools not usable to researches. We fixed them and have also provided the corrected scripts for such tools. We expect this study to serve as a guide for better understanding of software tools’ approaches for plant specific TFBRs discovery and the care to be taken while applying them, especially during cross-species applications. The corrected scripts of these software tools are made available at https://github.com/SCBB-LAB/Comparative-analysis-of-plant-TFBS-software.

## Introduction

Transcription factor (TF) proteins play a crucial role in gene regulation, binding to specific DNA regions known as transcription factor binding regions (TFBRs). Additionally, the presence of co-factors, cooperative TF-DNA binding **[1]**, chromatin accessibility **[2]**, nucleosome occupancy **[3]**, indirect co-operativity via competition with nucleosomes **[4]**, pioneer TFs **[5]**, flanking region environment **[6,7]**, and DNA methylation **[8]** have been reported recently to affect the TF-DNA binding specificity. TFs may also detect the shape features of their binding regions, commonly referred to as “shape readout” **[9,10]**.

TFs bind to their binding sites through their DNA-binding domain (DBD) **[11]** typically containing an oligomerization site, a transcription regulatory domain, and a nuclear localization signal **[12]**. However, the DBDs of TFs are not sufficient or necessary for their localization to most of their target promoters. Instead, binding to the majority of target promoters relies additively on numerous others factors spread across their entire intrinsically disordered regions (IDRs), which are a long, flexible polymeric tail comprised of hundreds of amino acids **[13,14]**. Information from sequence similarity and their tertiary structure with a DBD is frequently used to categorize the TFs into different families **[15–18]**. With the increasing number of three dimensional DBD structures being solved or predicted, a hierarchical classification for plant TFs is now possible **[15,19–21]**. Hence, among 56 recognized plant TF types, 50 were classified into one of the nine superclasses: Plant-TFClass Helix–turn–helix (HTH) domains, Beta-hairpin exposed by an alpha/beta-scaffold, all-alpha-helical DNA-binding structures, Beta-sheet binding to DNA, Zinc-coordinating DNA-binding domains, Beta-barrel DNA-binding domains, Basic domains, Alpha-helices exposed by beta-structures, Immunoglobulin-fold, and the remaining six were added to the ‘Yet undefined DBD’ **[15]**. Since then, comparative genomic studies have identified additional TFs and putative TF families in plant genomes **[22]**. Although some of these classifications acknowledge homology between different families, none of them propose a hierarchical, higher-rank organization. Model plant species like Ar*abidopsis thaliana,* have approximately 2,300 TFs, constituting 8.3% of its total genes, with 45% not found in mammals **[18]**. Crop species like wheat (Tr*iticum aestivum),* rice (Or*yza sativa),* and canola (Br*assica napus)* **[23–25] a**lso have a similar percentage of 5.7%, 6.5%, and 6.1%, TFs in their genomes, respectively. Unlike animals, plants have undergone drastic horizontal expansion which is the case of multi-copied genes in plants where copies also vary among themselves. The same has reflected to all elements of plant genomes including the TFs which too display high variability **[11,26–31]**. This is why plants have several members in any given TF family, and at the same time their binding preferences also vary a lot. The abundance, variety, and diversity of DNA-binding specificity in plant TFs suggest a potentially highly layered role for TFs and their transcriptional regulations in plants **[11,26–31]**. H**e**nce, genomes of plants contain a wide variety of TF-encoding genes which include a group of plant-specific TFs **[11,26–31] t**hat evolved over time.

The progress in TF-DNA interaction studies has been substantial for both experimental and computational approaches as illustrated in **Figure 1** and **Supplementary Material S1 Table 1**. *In vitro* methods like SELEX **[32]**, and protein binding microarrays (PBM) **[33]**, facilitate the characterization of low-affinity TFBRs. *In vivo* methods such as ChIP-Chip **[34]** and ChIP-seq **[35]** highlight specific protein-DNA interactions in natural contexts. Advanced variants of ChIP-seq like ChIP followed by exonuclease digestion and sequencing (ChIP-exo) **[36]** and ChIP-nexus **[37]** enhance resolution and accuracy. However, in an effort to reduce some of the inherent biases and limitations of ChIP-seq, an *in situ* chromatin profiling technique named Cleavage under targets and release using nuclease (CUT&RUN) was developed as an alternate but comparable technique **[38,39]**. In addition to these, DAP-seq **[40]** provides only static images of TF binding regions, as it constructs cistrome and epicistrome maps for various species by combining *in vitro* production of affinity-purified TFs with next-generation sequencing of a genomic DNA library. These high-throughput sequencing experiments generated >10,000 individual datasets, which are available in the NCBI SRA, covering more than 40 plant species. Among which approximately 790 plant ChIP and DAP-seq datasets are publicly available at Plant Cistrome Database **[41]**, *Arabidopsis* RegNet **[42]**, PlantPAN 4.0 **[43]** PCBase 2.0 **[43]**, and ChIP-Hub **[44]** databases for *Zea mays*, *A. thalaiana*, *Oryza sativa*, *Glycine max*, *Solanum lycopersicum*, and *A. lyrata*. The ChIP-seq and DAP-seq dataset retrieved from these resources are useful for building computational model for TFBRs and associated binding motifs across the TFBRs.

**Figure 1:**
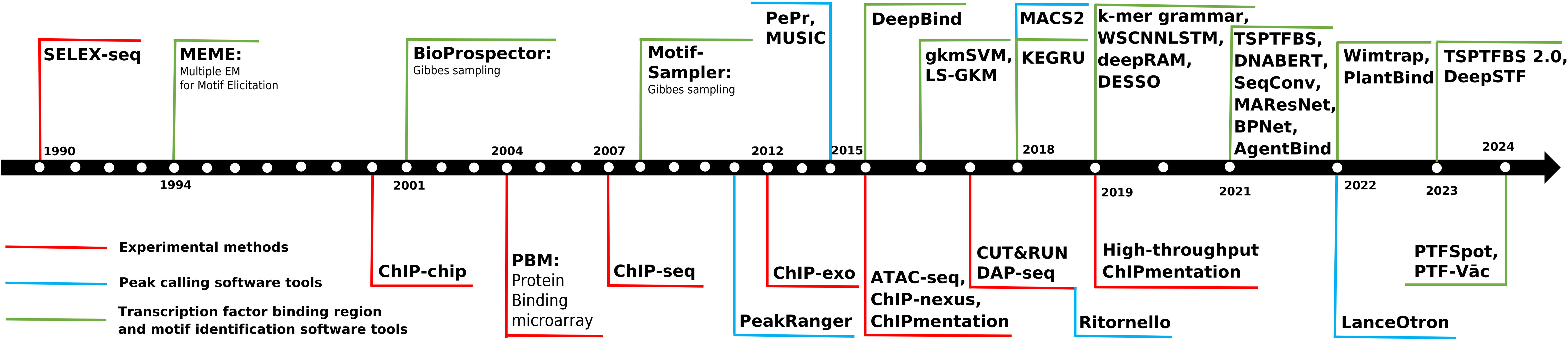
Timeline of the experimental and computation approaches. A timeline of the major milestones in the area of transcription factors and their binding regions research. As can be seen here, PWM based approaches to detect TFBRs have remained prime. High-throughput techniques at the later stage revolutionized the way and volume of data generation for downstream analysis of TFBRs identification.

**Table 1:** List of existing software tools for transcription factor binding regions identification used for comparative benchamrking.

| S.No. | Software Tools | Algorithm | Year | Standalone/<br>Webserver<br>(S/W) | Species | Datasource |
| --- | --- | --- | --- | --- | --- | --- |
| 1 | DeepBind [60] | CNN | 2015 | S | Human | ENCODE<br>project [96] |
| 2 | gkmSVM [58] | SVM | 2016 | S |  |  |
| 3 | LS-GKM [59] | SVM | 2016 | S |  |  |
| 4 | KEGRU [97] | RNN | 2018 | S |  |  |
| 5 | *k*-mer grammar [77] | Bag-of-*k*-mers | 2019 | S | Plant | Two MNase HS TFs, two ChIP-seq, core promoter [103-108] |
| 6 | WSCNNLSTM [99] | CNN+RNN | 2019 | S | Human | ENCODE project |
| 7 | deepRAM [61] | CNN/RNN | 2019 | S |  |  |
| 8 | DESSO [62] | CNN | 2019 | S |  |  |
| 9 | TSPTFBS [63] | CNN | 2021 | S | Plant | Plant Cistrome Database [41] |
| 10 | DNABERT [47] | Transformer | 2021 | S | Human | ENCODE project |
| 11 | SeqConv [79] | CNN | 2021 | S | Plant | GSE137972 [86] |
| 12 | MAResNet [98] | CNN+RNN | 2021 | S | Human | ENCODE project |
| 13 | BPNet [101] | CNN | 2021 | S | Mouse | Embryonic stem cell (ChIP-nexus and ChIP-seq) |
| 14 | AgentBind [64] | CNN and BiLSTM | 2021 | S | Human | ENCODE project |
| 15 | Wimtrap [78] | XGBoost | 2022 | S | Plant | *Arabidopsis* RegNet [42] |
| 16 | PlantBind [80] | Attention-based | 2022 | S | Plant | Plant Cistrome Database |
| 17 | TSPTFBS 2.0 [65] | DenseNet | 2023 | S/W | Plant | Plant Cistrome Database, GSE137972 and ChIP-Hub [44] |
| 18 | DeepSTF [100] | CNN, Transformer encoder, and Bi-LSTM | 2023 | S | Human | ENCODE project |
| 19 | PTFSpot [66] | DenseNet + Transformer | 2023 | S/W | Plant | ChIP-seq and DAP-seq |

As depicted in **Figure 2**, with the increase in high-throughput experimental techniques and subsequently volume of experimentally validated data has presented numerous computational challenges. These challenges can be broadly classified into three major stages in TF-DNA interaction studies: 1) Correctly identifying the TF binding peaks from the binding experimental datasets, 2) Modeling the TF-DNA binding regions from the identified peak datasets, and 3) Capturing the TF-DNA binding motifs or biologically relevant motifs from these binding regions **[45]**. First two stages are highly critical and most important ones. Though, the second and third stages of computational challenges are usually dealt together, both the problems are very different from each other in actual. They need to be seen separately. The first challenge involves obtaining optimal peaks from experimental binding data collected from techniques listed in the **Supplementary Material S1 Table 1**, as the data may be susceptible to bias and noise **[46]**. A crucial step is to reduce such bias in high-throughput datasets, which can significantly impact subsequent downstream computational and biological analyses **[46]**. Peak calling involves multiple separate analyses like read shifting, background estimation, enrichment peak identification, significance analysis, and artifact removal. Peak calling approaches for identifying binding regions in ChIP-seq datasets have been developed, as listed in **Supplementary Material S1 Table 2**. To assess the effectiveness of ChIP-seq peak callers, the present study focused upon some of contemporary tools for peak finding as listed in **Supplementary Material S1 Table 2**. These tools have been studied for their merits and demerits while evaluated for various parameters. Based on that a final recommendation for their most reliable use has been given.

**Figure 2:**
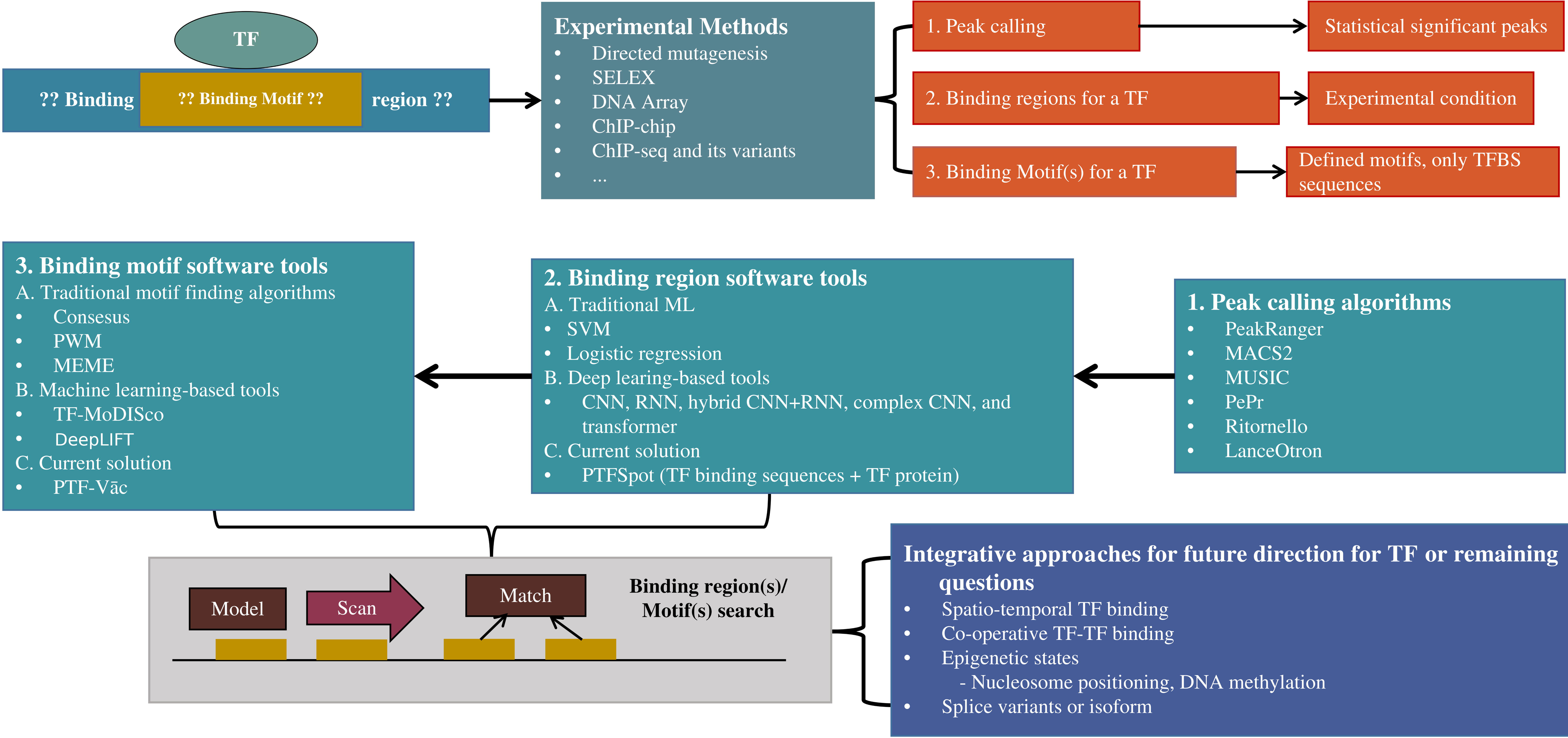
Outline of the study. The primary aim of this study is to comprehensively identify transcription factor (TF) binding regions and motifs across the entire genome. Initially, a limited set of experimentally determined regions serves as the foundation. Subsequently, (1) peak calling techniques are applied to detect regions of heightened TF activity, (2) followed by the construction of models for binding regions and (3) motifs using a combination of traditional methods and advanced deep learning software tools. These models are then employed in a genome-wide scan to uncover additional instances of TF binding sites and motifs. Besides enhanced binding sites and motif models, additional, genomic TF binding in response to spatio-temporal, co-operative TF-TF binding epigenomic, and differential binding by splice variants or the isoform of TFs data can be used in an integrative fashion to improve the accuracy of binding region or motif search.

**Table 2:** List of major errors encountered in benchmarking tools during program execution.

| S.No. | Type of Errors | Benchmarking tools | Major Errors |
| --- | --- | --- | --- |
| 1 | ModuleNotFoundError | KEGRU | cannot import name 'pad_sequences' from 'keras.preprocessing.sequence' (/home/csir/.local/lib/python3.8/site-packages/keras/preprocessing/ |
|  |  |  | sequence.py)<br>No module named 'keras_preprocessing'<br>No module named 'keras.layers.embeddings' |
|  |  | *k-mer* grammar | cannot import name 'joblib' from 'sklearn.externals' (/home/csir/miniconda3/lib/python3.10/site-packages/sklearn/externals/__init__.py)<br>No module named 'sqlalchemy' |
|  |  | WSCNNLSTM | No module named 'ANDNoisy'<br>No module named 'keras.engine.topology' |
|  |  | SeqConv | cannot import name 'batch normalization' from 'keras.layers.normalization' (/home/scbb/.local/lib/python3.8/site-packages/keras/layers/normalization/__init__.py)<br>No module named 'keras.layers.advanced_activations' |
|  |  | DESSO | No module named 'constants' |
|  |  | PlantBind | No module named 'google.protobuf' |
| 2 | TypeError | KEGRU | Word2Vec.__init__() got an unexpected keyword argument 'size',<br>int() argument must be a string, a bytes-like object or a real number, not 'NoneType'<br>Model.fit() got an unexpected keyword argument 'nb_epoch'<br>can only concatenate str (not "NoneType") to str |
|  |  | SeqConv | 'float' object cannot be interpreted as an integer<br>'dict_keys' object does not support indexing |
|  |  | PlantBind | '>=' not supported between instances of 'str' and 'float' |
| 3 | KeyError | WSCNNLSTM | 'acc', 'val_acc' |
| 4 | logicError | KEGRU | *k-mer* in lower case |
| 5 | MemoryError | WSCNNLSTM | tensorflow.python.framework.errors_impl.InternalError: Failed copying input tensor from /job:localhost/replica:0/task:0/device:CPU:0 to /job:localhost/replica:0/task:0/device:GPU:0 in order to run _EagerConst: Dst tensor is not initialized. |
| 6 | FileNotFoundError | KEGRU | No such file or directory: '/home/szhen/szhen/DBLSTM/test2/init_pos_5gram_2stride' |
|  |  | SeqConv | can't open file 'prepare.py': [Errno 2] No such file or directory<br>No such file or directory: 'fa/zm.fa'<br>No such file or directory: 'train/zm_tf_train.txt'<br>can't open file 'generator.py': [Errno 2] No such file or directory<br>No such file or directory: 'maize4.chrsize'<br>can't open file 'SeqConv.py': [Errno 2] No such file or directory |
|  |  | TSPTFBS | No such file or directory: 'Example/ABF2.fasta'<br>No such file or directory: 'Example/neg.fasta' |
|  |  | PlantBind | can't open file 'PlantBind/main.py': [Errno 2] No such file or directory |
|  |  | TSPTFBS 2.0 | No such file or directory: '/home/hlcheng/corn/sample/neg_sample.npy' |
| 7 | RuntimeError | DNABERT | CUDA error: no kernel image is available for execution on the device |
|  |  | PlantBind | Default process group has not been initialized, please make sure to call init_process_group. |
| 8 | OSError | DNABERT | 'PATH_TO_DNABERT_REPO/src/transformers/dnabert-config/bert-config-6/config.json' was not found in model name list. |
| 9 | Error in install.packages | gkmSVM | Error in install.packages("ROCR", "kernlab", "seqinr") |
| 10 | AttributeError | *k*-mer_grammar | 'TfidfVectorizer' object has no attribute 'get_feature_names'. Did you mean: 'get_feature_names_out'? |
|  |  | WSCNNLSTM | del_num = range(len(history_all)): 'range' object has no attribute 'pop' |
| 11 | Packages versions errors | BPNet | Required specific version of python3.6 and required packages in specific versions |

The outcome from the first stage of the Peak-calling problem is the direct input to the next stage of the problem solving: Development of computational models for TFBRs from the qualified binding data. Such models thereafter allow us to computationally detect the TF binding regions across the genome(s) instead of relying upon time and money consuming costly experiments. To address the second problem for TFBRs identification, as highlighted in **Figure 2**, a range of of published software tools to identify TFBRs, have been studied to determine their usability, issues, strengths, and performance, and recommendations for their uses. Most of these tools are predominantly based on traditional machine learning (ML) and Deep-Learning (DL) algorithms (**Supplementary Material S1 Table 3**). DL and ML approaches, both supervised and unsupervised, can be used to find TFBRs. In supervised learning, models that identify binding regions based on sequence features are trained using labeled data. Without the need for labeled data, unsupervised techniques like principal component analysis (PCA), autoencoders, and various clustering algorithms are used to find hidden patterns However, the application of purely unsupervised DL methods for TFBR discovery presents significant challenges, and currently, there are no tools based solely on unsupervised learning for this task **[47–49]**. Also, a recent study found that the unsupervised approaches are not suitable to detect TF-DNA interactions **[50]**.

**Table 3:** Comparative list of TFBS structural variance and structural and docking analysis for *Homo sapiens* TF protein and binding site and *Homo sapiens* TF protein and *Mus musculus* binding motif.

| Animal |  |  |  |  |
| --- | --- | --- | --- | --- |
| S.No. | Name of TF | Docking score |  | TOMTOM (p-value) |
|  |  | *Homo sapiens* TF protein and binding site | *Homo sapiens* TF protein and *Mus musculus* binding motif |  |
| 1 | CREB1 | -87.663 | -83.25 | 4.61132E-05 |
| 2 | FOXA2 | -103.36 | -101.972 | 1.27124E-06 |
| 3 | IRF1 | -130.24 | -128.384 | 1.05945E-06 |
| 4 | PAX5 | -114.7 | -112.45 | 8.82535E-10 |
| 5 | TBXT | -97.57 | -96.41 | 6.50144E-05 |

Initially the algorithm development mainly involved on traditional techniques for motif findings employing various statistical approaches like Gibb’s sampling and Expectation maximization were used **[51–55]** as very limited experimental binding data would be generated. This also became one of the founding reasons why finding TFBR and TFBS were seen as same problem despite being drastically different as emerges now. The identified regions by these approaches were used to develop position weight matrices which would scan the sequences to identify TFBR/TFBS for any given transcription factors. At first such approach look supervised clustering approaches, but in actual they need at least and essentially the specific information for the binding TF besides several random tweaking of parameters by the user itself. Therefore, it is really difficult to claim that tools attained purely unsupervised learning on data to identify TFBS/TFBR. The area further matured and involved ML approaches **[47]**. For instance, Wong *et al*. proposed a method, *k*mer HMM, employing belief propagation based on Hidden Markov Models (HMMs) **[56]**, another unsupervised approach. Ghandi *et al.* introduced gkmSVM, a gapped-*k*mer support vector machine (SVM) classifier, to train on protein and DNA sequences **[57–59]**. The groundbreaking work of DeepBind in 2015 introduced DL approaches to identify protein-DNA binding regions **[60]**. DL models like Convolutional Neural Networks (CNNs) can automatically learn crucial features directly from the input data, unlike traditional ML techniques that often require manual feature engineering. Subsequent software tools developed for TFBRs identification followed the basic principles of the DeepBind algorithm to construct their model architecture **[61–64]**. Most recent and advanced software tools, such as TSPTFBS 2.0 **[65]**, DNABERT **[47]**, and PTFSpot **[66]**, have utilized some much evolved CNN architectures and transformers.

Time to time grasping on latest biological findings have improved these algorithms. The area which had started with merely reporting the motifs and calling it TFBS, started to acknowledge the role of multiple motifs in DNA binding and thus *k*-mers based information infusion happened **[56,67,68]**. Some studies revealed that the flanking regions surrounding potential interaction sites in nucleic acids effectively capture the contributory information from the local environment **[6,7,69–73]**. These regions are crucial for determining the positioning of TFs, as they facilitate a localized search for appropriate binding regions **[72]**. The contextual information may include co-occurring motifs, sequence and position-specific details, as well as structural or shape information. One such acknowledgment was done recently while using DNA shape information in contextual manner involving flanking regions around the TFBS to better identify the TF binding regions while employing a ML approach of Random Forests **[73]**. Therefore, contextual factors emerge as very strong discriminators between negative and positive instances of transcription factor binding. Progress is this direction appear very crucial for any successful algorithm development to detect TF-DNA interactions.

Now coming to one of the most crucial issues of this area and development is the enormous skew and lack of universal approach in finding TF-DNA interactions. As evident from **Supplementary Material S1 Table 3**, out of 38 software tools, 30 were developed using common human experimental ChIP-seq, ChIP-exonuclease, ChIP-nexus, ATAC-seq **[74]**, DNase-seq **[75]**, and PBM binding data. Only eight tools have been developed for plant-specific TFBRs identification. Three of them were traditional ML based **[76–78]**, while five were DL based **[63,65,66,79,80]** software tools developed to discover plant TFBRs. Thus, as reflected by the gap in the volume of experimental binding data between vertebrates and plants, similar huge lag in terms of plant specific tools was also observed. And as will transpire in the results and discussion below, hardly any of these plant specific tools used plant specific biology to improve their algorithms. They have just carried on the same philosophy and algorithms which were mainly developed at animal datasets specifically. The only plant related thing which appears mainly incorporated is the incorporation of plant specific binding datasets for training and learning without any major addressing of biological properties which could make them really relevant to plants or the issues faced in detection of plant specific TF-DNA interactions. This is why, as will reflect in the study done in this work, when animal based tools were trained in the same plant based data we found them performing at par of those which claimed themselves as plant specific tools.

Unlike animal, where most of focus has been on human, the plants pose entirely different set of challenges, which largely unaddressed **[11,26–31]**. As will transpire in large in the discussion section, this is important to note that there is a significant difference between animals and plants genomes in terms of variability. Plants display much higher variations for most of the genomic elements **[11,26–31]**. Compared to animals, plants have wide range of complex repeats which also display enormous variability in their coverage and forms **[81,82]**. Thus, it is very likely that a TF specific model built for one species may not work in other. Ironically, like with vertebrates where human has been the center of all major attention, in plants most work has been centered around the model species of *Arabidopsis thaliana*, a small genome plant with equally small nondescript genomic complexity. Most of the plant based models have been developed mainly for this species, which are thereafter used across other species to detect TFBS/TFBR, which has been an immensely wrong practice done for years, as reported recently **[66]**. The present work has also looked into this aspect which has actually compounded the issues with plant TFBR/TFBS discovery immensely and guides towards the recent solution to all this.

The final matter dealt in this study is the 3^rd^ stage of computational challenges, motif discovery from the TFBRs (**Figure 2)** which has been intertwined with the problem of TFBR discovery for a long time, despite being a different category of problem. Interestingly, as also discussed above, TF-DNA interactions discovery started with this same problem of motif finding and had been largely limited for a very long time to this one only, grossly neglecting the fact that TF-DNA interaction is more about context and surroundings than just locating some motifs. With shifts in our understanding towards TF-DNA interactions, we are now at a state where TFBRs reported from the previous computational steps or binding data are used to report any such motifs. Most of the existing approaches primarily focus on identifying common over-represented patterns or motifs for TF-DNA binding, for which several computational resources and algorithms have been developed over the years. For a long time it has been believed that motif found in the bound DNA data were the binding factors to TFs. And thus for most of the TF binding region discovery, finding such motifs and calling them TFBS became a norm **[60–65,77–80]**. However, that could be a wrong practice done for a long time as several other factors besides enriched motifs have been found responsible **[66,76]**. The reported TFBS motifs occur substantially even in non-binding data also. And, as discussed above, in plants such TFBSs and TFs have been found displaying extensive divergence, resulting into highly variable binding preferences and motifs **[11,26–31]**.

The current motif discovery approach are more like a statistical practice, which has been made default to TFBR discovery **[61,62,83,84]**. Motifs need be dealt just like a potential marker. However, motif finding reports need to be decoupled from TFBR discovery and should be done once a TFBR is notified. Off late such practice has got attention with DL methods for TFBR detection, where motifs are identified after TFBRs confirmation. Yet, all of them primarily take statistical enrichment route in their core, and are highly sensitive towards the volume of data provided. The last part of the present work highlights some of the approaches made in the area and finally discusses a generative deep learning solution to motif finding with much higher credibility and non-sensitivity towards data volume as well as noise in such data. **Figure 2** gives an abstract snapshot of the overall study undertaken here.

## Material and methods

### Dataset for peak calling

The first stage was to analyze peak calling tools, which examine the distribution of TF-DNA binding across the entire genome using ChIP-seq and DAP-seq dataset. The peak calling process consists of three crucial steps: pre-processing, mapping, and peak finding **[85]**. During pre-processing, low-quality and incorrect reads are eliminated. In the mapping stage, reads are aligned to the reference genome, potentially resulting in duplicated reads mapped to different genomic sites. The mapping strategy can prioritize either unique reads for specificity or multiple read alignments for sensitivity. We downloaded FASTQ formatted raw sequencing ChIP-seq data for five TFs in *Zea mays*, namely MYB62, bZIP9, WRKY94, HB33, and EREB147, with control datasets from a specific study **[86]**, which were obtained from public repositories or experimental sources using enaBrowserTools **[87]**. Subsequently, adapter sequences were removed, and low-quality reads were filtered using Trimmomatic to ensure data integrity and quality for downstream analysis. Following pre-processing, the alignment tool Bowtie2 was employed to map the reads to the *Zea mays* reference genome. SAMtools was then used to convert the reads mapping file to sorted BAM format. Regions of TF enrichment were identified using different peak calling tools. A total of six tools were considered (PeakRanger, PePr, MUSIC, Ritornello, MACS2, and LanceOtron) **[88–93]**. These tools employ different statistical approaches for the peak calling to calculate read count densities and locate binding locations. The output of peak calling tools includes a list of peak sequences in various formats and sizes. Additionally, the noise inherent in peak-calling from TF ChIP-seq experiments requires the use of the Irreproducible Discovery Rate (IDR) framework **[94,95]**. This method adaptively thresholds and retains peaks that are reproducible and rank-concordant across multiple replicates. Subsequently, significant peaks were obtained from the tools, for which they used different statistical tests to finalize the peak. These peak sequences often contain short motifs serving as binding sites for particular TFs.

### Dataset for transcription factor binding regions

In the field of ML research, constructing a high-quality dataset is an essential and initial requirement. This dataset guides not only the quality of the raised model, but also their validity and generalization capabilities. Selection of suitable instances as positive and negative components, data cleansing, labeling, and resolving class imbalances are major parts of this crucial step. The dataset is often separated into training, validation, and test subsets to ensure reliable model building and evaluation. In this direction, the primary step of this study was to develop benchmarking datasets.

These datasets primarily comprise experimentally validated genomic regions where TFs bind, often identified through techniques like ChIP-seq and DAP-seq. Project like ENCODE (Encyclopedia of DNA Elements) has compiled extensive information on functional DNA sequences from diverse tissues and cell types in the human genome **[96]**. Additional TFs’ data information is accessible through various databases, including those mentioned in **Supplementary Material S1 Table 4**. Notably, repositories such as the Plant Cistrome Database, *Arabidopsis* RegNet, ChIP-Hub PlantPAN 4.0, and PCBase 2.0, offer comprehensive collections of such TF binding region datasets for multiple plant species. For our study, we gathered a comprehensive set of datasets for plant and animal TFs from some of these resources. This include a substantial volume of ChIP and DAP-seq data for *A. thalian*a and ChIP-seq data for *Zea mays* and *Oryza sativa* to conduct a fair evaluation of different software tools for TFBRs identification in plants and across other plant species. This dataset, previously utilized by tools like TSPTFBS, PlantBind, and TSPTFBS 2.0, encompasses peak data for 35 plant TF families (**Figure 3**).

**Figure 3:**
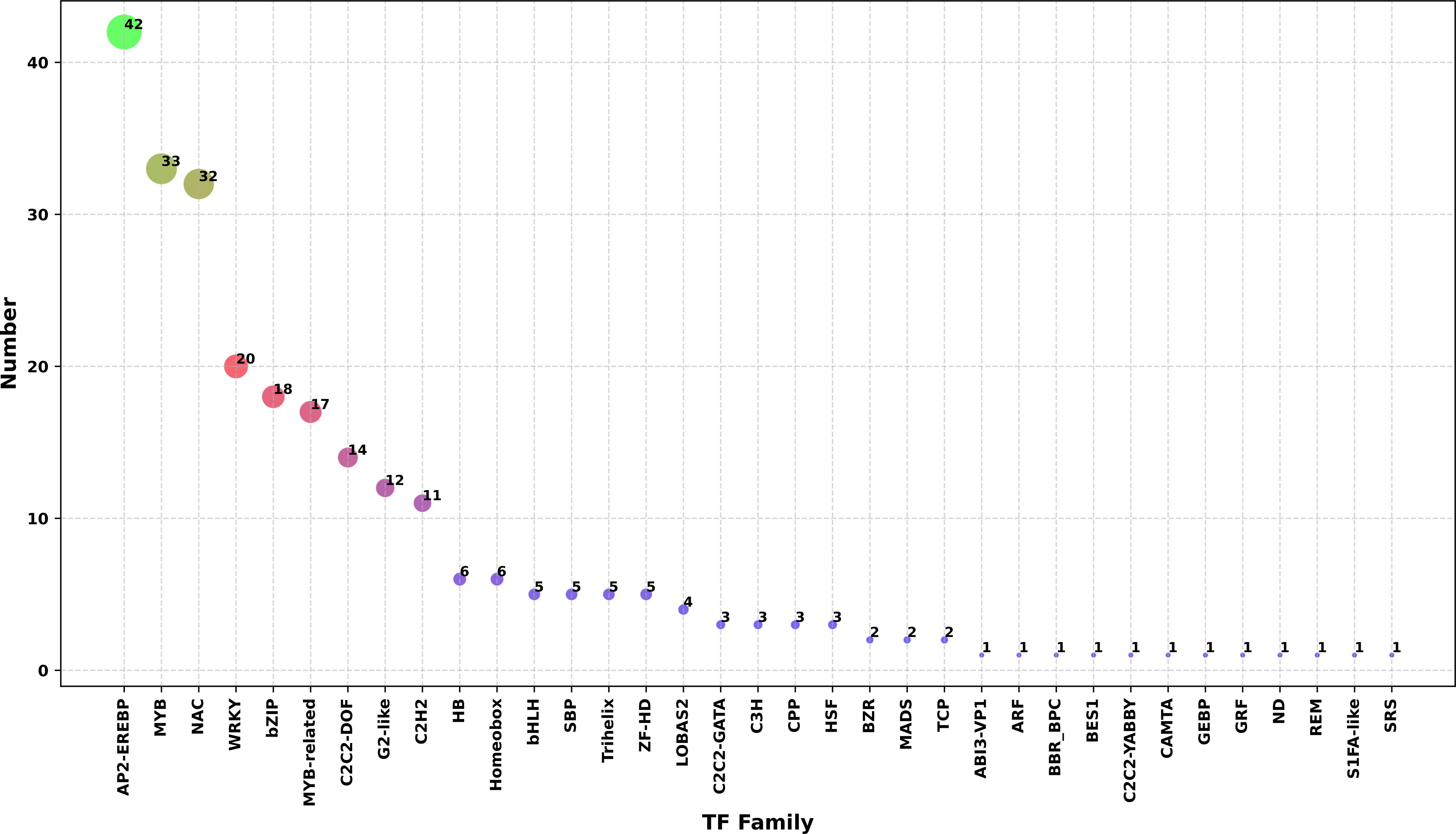
The family wise distribution of 265 transcription factors data for the Dataset “B”. Some of the transcription factors exhibit clear bias. Such bias is also due to more experiments taken for a particular family, and does not reflect the actual abundance in cell system.

**Table 4:** Comparative list of TFBS structural variance and structural and docking analysis for *Arabidopsis thaliana* TF protein and binding site and *Arabidopsis thaliana* TF protein and *Cannabis sativa* binding motif.

| Plant |  |  |  |  |
| --- | --- | --- | --- | --- |
| S.No. | Name of TF | Docking score |  | TOMTOM (p-value) |
|  |  | *Arabidopsis thaliana* TF protein and binding site | *Arabidopsis thaliana* TF protein and *Cannabis sativa* binding motif |  |
| 1 | bZIP18 | -94.542 | -85.949 | 0.0992451 |
| 2 | AT3G57600 | -93.568 | -84.661 | 0.0970465 |
| 3 | WRKY3 | -102.81 | -73.612 | 0.0919656 |
| 4 | ERF48 | -105.059 | -79.665 | 0.0808847 |
| 5 | CRF10 | -59.994 | -48.362 | 0.422613 |

To ensure the successful completion of our study, we systematically evaluated the performance of 19 existing software tools across three different types of datasets. Firstly, Dataset “A” was constructed which contains entries for which the compared software tools were originally developed. This initial phase aimed primarily to validate the performance claims made by these tools. For human TFBRs identification, the majority of software tools had used ChIP-seq datasets from the ENCODE project. Existing tools designed specifically for plant TFBRs identification utilized a variety of datasets including ChIP-seq **[77–79,86]**, micrococcal nuclease hypersensitivity (MNase HS) **[77]**, core promoter **[77]**, and DAP-seq data from sources such as the Plant Cistrome Database and *Arabidopsis* RegNet.

In the second phase of the study, we focused on model development and conducted objective assessments of all the 19 tools, designed for both animal and plant-based, using a plant-specific DAP-seq dataset. This part aimed to establish robust models and provide unbiased evaluations of the tools’ performance. Finally, in the third phase of the comparative analysis, we investigated the performance of pre-trained models on *A. thaliana* in a cross-species manner. This allowed us to assess their applicability for cross-species applications and further understand their versatility across different species.

### Positive dataset creation

Various tools have been utilized for identifying different types and numbers of TFBRs (**Table 1**). The focus of the initial part of this study was to verify the respective claimed performance of 19 different software tools. Most of these tools were primarily designed for human TFBRs identification (**Supplementary Table S1 Sheet 1**), including DeepBind, gkmSVM, KEGRU **[97]**, MAResNet **[98]**, WSCNNLSTM **[99]**, DeepSTF **[100]**, DNABERT **[47]**, BPNet **[101]**, and AgentBind **[64]**. The ChIP-seq experimental data for human TFBRs, which was obtained from the ENCODE project and processed following the documented methodologies of each software tool (**Figure 4A**). The software tools such as DESSO, MAResNet, and DNABERT utilized 690 experimental ChIP-seq datasets, representing 137 unique TFs. Other tools such as DeepBind, gkmSVM, LS-GKM, KEGRU, and DeepSTF used the data for 506, 467, 322, 125, and 165 ChIP-seq experiments, respectively. BPNet had used the pluripotency TFs Oct4, Sox2, Nanog, and Klf4 using the well-established mouse embryonic stem cell (ESC) model, for ChIP-nexus **[102]** and ChIP–seq data. AgentBind had employed ChIP-seq datasets from ENCODE project, and concentrated on 38 TFs that were active in GM12878 cell line. DESSO, deepRAM, and DNABERT employed a methodology similar to DeepBind for constructing positive datasets.

**Figure 4:**
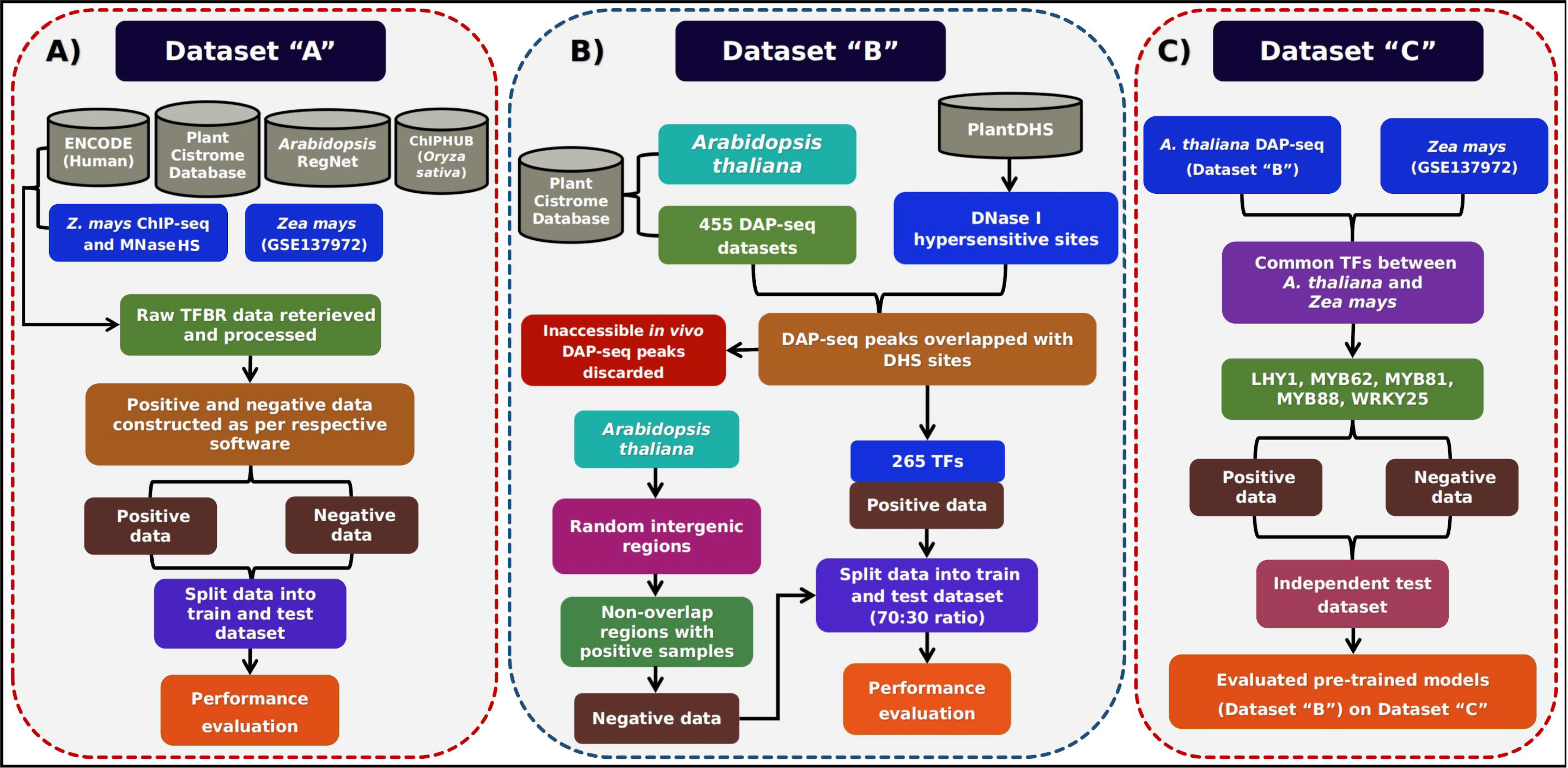
Flowchart representation of dataset formation. **(A)** The protocol followed for Dataset “A” creation, (**B)** Dataset “B” creation, and (**C)** Dataset “C” creation. The Dataset “A” was created for human and plant transcription factors. This dataset includes sequences from *Homo sapiens*, *Arabidopsis thaliana*, and *Zea mays*. The positive and negative instances within this dataset were generated following the same procedures as outlined in the respective implementing tools. The Dataset “B” was specifically designed for 265 *A. thaliana* TFs. In this case, random intergenic sequences were employed to produce negative datasets for each TF while ensuring the removal of any overlapping regions. As for the third dataset, Dataset “C”, it contained only five common TFs, named LHY1, MYB62, MYB81, MYB88, and WRKY25. This dataset serves as an independent test dataset to evaluate the software tools’ performance cross-species.

For the plant based TFBRs identification tools, peak files were retrieved from the Plant Cistrome Database, as well as from plant ChIP-seq and MNase HS experiments (**Supplementary Table S1 Sheet 1**). Additionally, specific strategies were adopted by some tools, for example, *k*-mer grammar focused on core promoter regions, binding loci from ChIP-seq peaks for two TFs (i.e., Homeobox KNOTTED1-KN1 and bZIP FASCIATED EAR4-FEA4), and open chromatin regions using MNase HS data from *Zea mays* **[103–108]**. SeqConv **[79]** used 104 maize TFBR datasets produced by ChIP-seq **[86]**, while TSPTFBS and PlantBind utilized large-scale DAP-seq datasets for TFs in *A. thaliana*. PlantBind aggregated all peaks obtained from 315 DAP-seq datasets to construct their positive dataset. The updated version of TSPTFBS 2.0 expanded its datasets which along with DAP-seq include ChIP-seq dataset for *Oryza sativa*, *Zea mays*, and *A. thaliana*, covering a total of 389 plant TFs. Here, PTFSpot utilized ChIP/DAP-seq peak data for 436 TFs, which were retrieved from PlantPAN 4.0 (for 54 TFs) and Plant Cistrome Database (for 387 TFs). PTFSpot constructed positive dataset, where sequences from the peak data were converted into instances after prime motifs were identified for each TF in the provided peaks data. Instances were generated by considering ±75 bases in both directions from the boundaries of the prime motif regions in both forward and reverse direction. In overall, this Dataset “A” encompassed sequences from human, mouse, *A. thaliana*, *Zea mays* and *Oryza sativa*.

In the second part of the this study, centered on Dataset “B”, we aimed to assess 19 existing software tools using a plant-specific TFBR dataset, enabling an objective evaluation of their performance and suitability for plant species. To accomplish this, we obtained the dataset utilized by Liu *et al.* **[63]**, which was downloaded and refined for model development across all software tools, incorporating DAP-seq data for 455 plant TFs from the Plant Cistrome Database for *A. thaliana*. To isolate high-confidence positive peaks, the DAP-seq peaks were overlapped with the DNase I hypersensitive sites (DHSs) to explore the relationship between chromatin accessibility **[75]**. DHSs represent regions of open chromatin, which are more favorable to TF binding. Consequently, TFBR datasets containing only unique peaks were retained and peaks inaccessible *in vivo* regions were discarded. After this procedure, DAP-seq data for 265 TFs were identified as the components for the positive dataset, which was utilized for training and evaluation purposes. As depicted in **Figure 4B**, we further refined the positive dataset for all 265 TFs by identifying non-overlapping regions with the negative instances for each TF. The positive instances ranged from 1,024 to 11,990 peaks, covering a total of 9,11,342 binding regions (**Supplementary Table S1 Sheet 2**).

### Negative dataset creation

The preparation of negative instances is a crucial step, arguably more so than that of the positive dataset, as it significantly impacts the model’s robustness, generalization, and real-world applicability. The quality of the negative dataset significantly influences the performance and reliability of the model **[6,66,109]**. Neglecting to construct a proper negative dataset can introduce biases, noises and lead to a misleading model. We noticed that the methodology for constructing the negative data for Dataset “A” varied across different tools.

Some tools, including DeepBind, deepRAM, MAResNet, and DeepSTF, generated negative instances by randomizing the positive sequences while preserving similar dinucleotide composition. In contrast, others tools such as gkmSVM, LS-GKM, KEGRU, WSCNNLSTM, DESSO, DNABERT, AgentBind, and *k*-mer grammar utilized random genomic sequences with matching repeat and GC content to the positive dataset. Notably, all these tools except *k*-mer grammar employed the “genNullSeqs” function of gkmSVM. This function identifies the nucleotide composition of a positive set of sequences and constructs a collection of genomic regions with comparable nucleotide composition and sequence length. Negative dataset regions for BPNet were retrieved from protein-attached chromatin capture (PATCH-CAP) for ChIP-nexus data. Specifically, for ChIP-seq negative dataset which consisting of 1 kb regions centered on the peak summits. TSPTFBS, TSPTFBS 2.0, and PlantBind opted to construct their negative datasets by randomly extracting intergenic regions of the *Arabidopsis* genome (TAIR10). Additionally, tools like Wimtrap **[78]** and SeqConv **[79]** selected random non-overlapping genomic sequences distinct from the positive instances. PTFSpot constructed the negative datasets by using the significant motifs found in ChIP/DAP-seq data (positive datasets) to scan them in the unbound regions in forward and reverse complementary motifs similar to the positive instances, with 75 bases flanking regions included for context **[7]**. Positive and negative instances were combined in a 1:1 ratio, while ensuring no overlap.

To ensure the robustness of Dataset “B”, we incorporated a significant pool of 1,89,799 genomic background sequences sourced from Liu *et al*. (2021), representing random intergenic regions of the *A. thaliana* genome (TAIR10). To refine this negative dataset further, we removed all overlapping regions between the negative and positive instances for each TF. These non-overlapping regions were then employed as the negative instances for each TF. It is noteworthy that many existing published software tools grossly neglected this refinement step, resulting in datasets where positive and negative instances overlap. This oversight compromises the reliability of the model and the reported performance **[110]**.

Another crucial factor influencing the performance of the model is dataset imbalance, where one class outweighs the other, leading to bias and class imbalance issues. To investigate the impact of class imbalance on model performance, we generated both balanced and imbalanced datasets for training. This approach allowed us to evaluate how the model performs under different data conditions and assess the effectiveness under class imbalance. We assessed the impact of using class-imbalance-aware loss functions in such scenario. The imbalanced datasets were generated with a 1:4 ratio of positive to negative samples for 265 TFs. We also generated balanced datasets and maintained consistent sequence lengths and sample numbers for both positive and negative instances across all 265 TFs. Throughout this study, this balanced dataset has been referred to as Dataset “B” (**Supplementary Table S1 Sheet 2**). Additionally, we ensured that there was no overlap between instances in the training and testing datasets to prevent memory and bias. However, to evaluate the performance of each software tool on this dataset accurately, we allocated 70% of the dataset for training purposes, while reserving the remaining 30% as a completely unseen test set. This standard approach ensures unbiased performance testing and eliminates the possibility of data memorization influencing the results.

In addition to Datasets “A” and “B”, we constructed an another independent test dataset named Dataset “C” to assess the generalizability of pre-trained models for cross-species TFBRs identification. The purpose of this cross-species study was to provide an understanding of relevant publications, applications falling under transfer learning, and associated challenges and solutions to the researchers interested in transfer learning. Due to data limitations for plant species, we found only five common TFs: LHY1 (MYB-related), MYB62 (MYB), MYB81 (MYB), MYB88 (MYB), and WRKY25 (WRKY), shared between *A. thaliana* and *Zea mays* (detailed in **Supplementary Table S1 Sheet 3**). For evaluating pre-trained models for these five TFs in *A. thaliana*, sourced from Dataset “B”, we tested them across the same TFs retrieved from Dataset “A” utilized by SeqConv. In the construction of Dataset “C” (depicted in **Figure 4C**), the ChIP-seq dataset for these common TFs in *Zea mays* was considered as the positive dataset. The negative dataset was randomly constructed from intergenic regions of *Zea mays*, ensuring no overlap with the positive instances. The test sequences from both positive and negative instances served as sources for objective comparative evaluation.

### Survey and categorization of the software tools to identify transcription factor binding regions

As already mentioned above, initially the TFBR discovery was limited to identifying motifs, which were detected using various unsupervised probabilistic approaches to raise profiles and scan sequences against them. Such approaches lacked the understanding of transcriptional regulation and context and flanking region’s while missing several biologically important features. However, recently developed computational approaches and algorithms to identify TFBRs are now capable to consider more relevant features, and can be classified into two major categories: Traditional ML and DL supervised and unsupervised approaches. The traditional ML algorithms learn from manually extracted the features through structured inputs. Traditional ML approaches can be classified into the most common types being SVMs, decision trees, and ensemble algorithms.

On the other hand, DL approaches are the most recent ones. They are founded on neural networks and differ from each other by means of their architecture and how they approach the input. Also, they automatically extract features and detect long range relationships and hidden features that are otherwise difficult to detect by manual and traditional feature extraction methods, as needed in non-DL methods. The DL methods can be categorized into six major categories: (i) Plain feed-forward networks, (ii) CNNs, (iii) Recurrent neural networks (RNNs), (iv) Complex CNNs, (v) Transformers (supervised and unsupervised), and (vi) Hybrid architecture which involves various combinations of above mentioned major architectures (illustrated in **Figure 5**). Fully connected feed-forward networks (FFNs) are integral parts of all DL architectures. For this study, we undertook 19 traditional ML and DL architectures listed in **Table 1**. These considered tools were categorized and are briefly explained below according to their algorithms.

**Figure 5:**
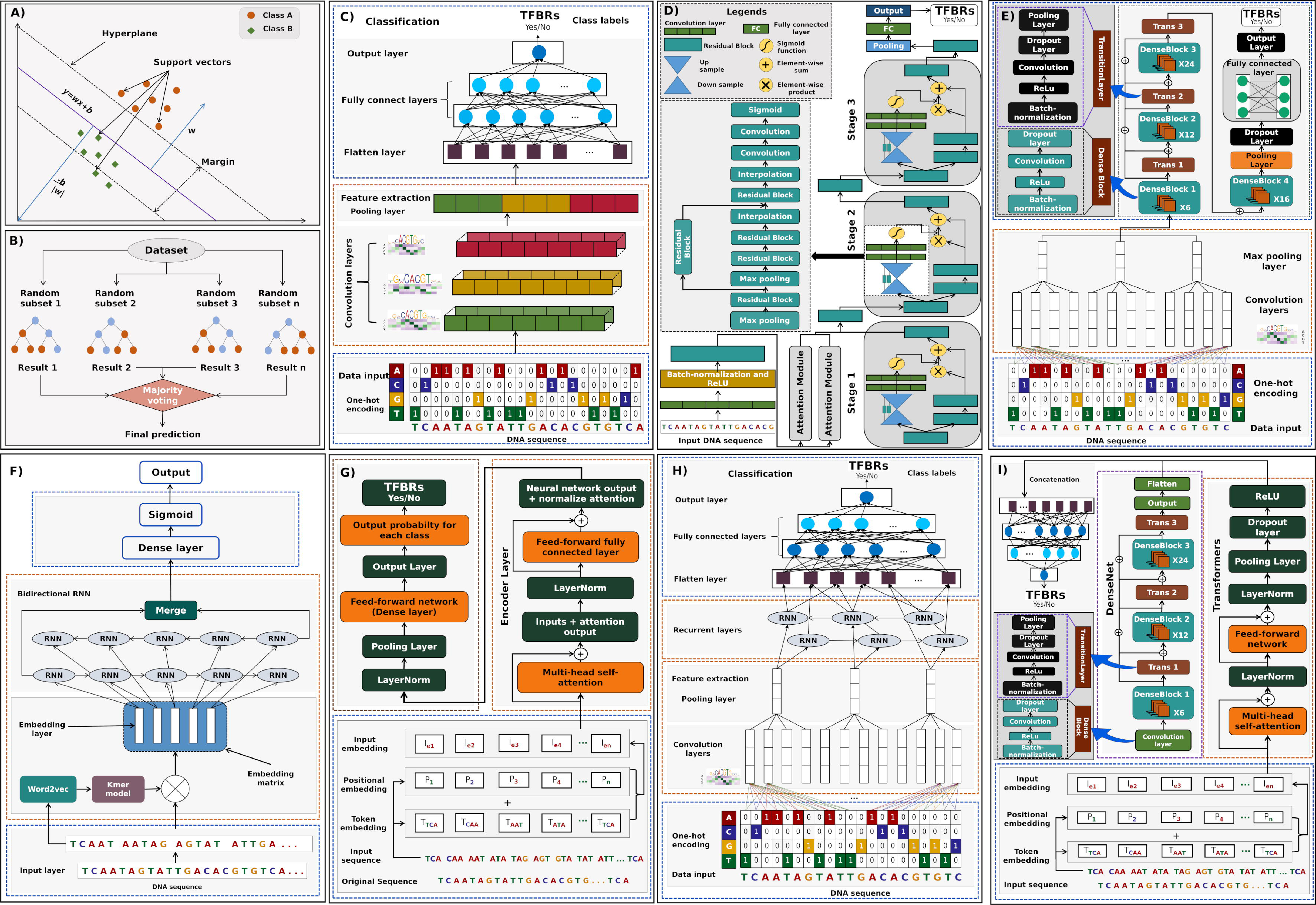
Overview of the machine and deep learning. A brief overview of the machine and deep learning architectures that have been used by 19 different software tools: **(A)** Support vector machine (SVM), **(B)** XGBoost, **(C)** CNN algorithm, **(D&E)** Complex CNN algorithms (ResNet and DenseNet), **(F)** RNN algorithm, **(G)** Transformer-based models, (**H)** hybrid CNN-RNN models, and **(I)** hybrid CNN and Transformer-based models. The input sequence is encoded as one-hot encoding (CNN and CNN-RNN hybrid), *k*-mer spectrum (SVM, RNN), and as token embedding (transformer based software tool).

### Traditional ML-based TFBRs identification software tools

The term “traditional machine learning”, a fundamental subfield of artificial intelligence, refers to a collection of statistical learning approaches for identifying and categorizing data based on well-defined features and established algorithms. Contrary to the recent DL techniques, traditional ML involves careful manual features’ extraction based on domain expertise and a variety of algorithms (**Figure 5 A&B**). Based on the type of learning algorithms applied, the TFBRs finding tools are discussed separately below.

**i) gapped-*k*mer-SVM (gkmSVM):** To account for variability in sequences and the length of TFBRs, gapped k-mers were introduced to represent longer motifs as *k*-mers with gaps at non-informative or degenerate TFBR positions (**Figure 5A**). By utilizing these gapped *k*-mers, gkmSVM enhances traditional *k*-mer-based SVM approaches, allowing it to capture more complex patterns that may include indels or mismatches within sequences. Implemented in C++ and R, gkmSVM functions as a binary classifier that learns to separate the input space using a hyperplane boundary. This tool incorporates three key steps: (i) Constructing the kernel matrix by calculating pairwise similarity scores using the “gkmsvm_kernel” function for all the sequences in the positive and negative sets, (ii) Training the SVM model with the “gkmsvm_train” function, and (iii) Classifying sequences using the “gkmsvm_classify” function. Additionally, gkmSVM includes the “genNullSeq” function, which generates a GC content, length, and repeat fraction-matched negative sequence set in the absence of a negative dataset. The “gkmsvm_delta” function is to evaluate the impact of sequence variants by calculating their scores.

**ii) LS-GKM:** LS-GKM is an updated version of gkmSVM, a SVM classifier designed to differentiate between regulatory and non-regulatory genomic sequences using *k*-mer or gapped *k*-mer frequency feature vectors (**Figure 5A**). In traditional SVM training, a kernel matrix, or Gram matrix, is calculated, containing all potential inner products between a collection of vectors for “*n*” training samples. However, as sample numbers increase, calculating this matrix directly with gapped k-mers becomes computationally infeasible. LS-GKM addresses this by implementing an efficient method to compute the complete kernel matrix with a run-time that grows linearly with “*n*” instead of the typical “*n²*”, enabling scalable SVM training on large sequence datasets. LS-GKM further optimizes performance by integrating gapped *k*-mer kernel functions within the LIBSVM framework **[111]** and introducing an approximation method for constructing the kernel matrix, significantly reducing computational complexity and memory requirements. Additionally, LS-GKM offers expanded kernels, including the radial basis function (RBF) kernel, referred to as gkmrbf-kernel and the center-weighted gkm-kernel (wgkm-kernel), which weights gapped *k*-mers based on their distance from the peak’s center. The combination of these kernels forms the “wgkmrbf-kernel”, and multi-threading functionality is added for further speedup. As a result, LS-GKM can efficiently handle much larger datasets than gkmSVM, with reduced time and memory usage.

**iii) *k*-mer grammar:** Mejía-Guerra and Buckler (2019) developed a computational framework which integrates potential TFBRs within genomic regions, utilizing k-mer sequence features. This tool integrates two natural language processing (NLP) approaches: (1) the “bag-of-words” model, to differentially weight keywords, and (2) a vector-space model using word2vec “vector-*k*-mers”, which captures semantic and linguistic relationships between words. The “bag-of-*k*-mers” model extracts information from individual *k*-mer frequencies and fits a logistic regression model using a matrix transformed by term frequency-inverse document frequency (TF*IDF) **[77]**. In contrast, the “vector-*k*-mers” model gathers data from *k-*mer co-occurrences by training a shallow neural network to predict *k*-mer likelihoods given their context. Each sequence in this model is treated as a series of “sentences” produced by sliding windows, with each sentence broken into distinct *k*-mers. Word2vec algorithms are used to create these models, optimizing an objective function via stochastic gradient descent (SGD) to fit *k*-mer vectors, which correspond to *k*-mer co-occurrence likelihoods. They also utilized the pre-trained models which were applied to TF binding loci and core promoter regions across different species, including maize, sorghum, and rice, shedding light on the significance of sequence elements and their interactions.

**iv) Wimtrap** was constructed as a novel, simplified software tool that aims to identify *cis*-regulatory elements (CREs) and target genes of plant TFs in specific organs or conditions. This method uses an XGBoost-based approach (an ensemble ML) (**Figure 5B**) implemented in the R programming language, enabling the development of decision rule models using experimental TF ChIP-chip/seq data. Different layers of genomic features are integrated in the predictive models: the position on the gene, DNA sequence conservation, chromatin state, and other CRE imprints. Wimtrap takes only positive data as an input in the form of the coordinates of the TF peaks. Chromatin characteristics were given high weight here to achieve better accuracy. A five-fold cross-validation was employed to report the model’s performance. The established features form the basis for the pairwise correlations between them. The feature with the highest mean absolute correlation with the other features is eliminated if two features exhibit a correlation greater than 95%. This method is mainly focused on plants, especially on *A. thaliana*, the model species for plant molecular biology, also for which data are the most abundant.

### Deep learning-based TFBRs identification software tools

While the traditional ML approaches are still very relevant and extensively utilized, it is important to note that DL, a subset of ML, has attracted a lot of attention and gained popularity recently owing to its exceptional performance in a variety of applications. This has greatly motivated the research community to apply them to molecular biology.

#### 1) CNN-based TFBRs identification software tools

The CNN is a type of feed-forward neural network with convolution estimation and depth architecture. It has been utilized effectively in identifying images, video analysis, natural language processing, and other fields with good results **[112,113]**. CNN typically works by inputting image information and passing it through a combination of convolutional layers, non-linear layers, pooling layers, and complete connection layers to get the final output result. As shown in **Figure 5C**, the convolutional layer is largely responsible for feature extraction via the convolutional kernel scan, whereas the pooling layer is primarily responsible for feature selection and information filtering. Because of its strong feature extraction capacity, CNN has been thought to have the capacity to identify the motif in the process of searching for TFBRs. Below are a few tools which are based on CNN architecture:

**i) DeepBind,** implemented in Python, was the first method to use CNNs to identify DNA/RNA-binding sites from high-throughput data (**Figure 5C**), such as PBM, SELEX, ChIP-seq, and CLIP-seq data **[114]**. Unlike traditional ML algorithms, DeepBind eliminates the need for manual feature engineering by automatically learning to extract key characteristics from input sequences and identifying hidden spatial features and associations. This spatial pattern detection is a significant strength of CNNs, yielding superior results in motif discovery. In DeepBind’s architecture, raw nucleotide sequences are first converted into a one-hot encoding (OHE) matrix. The data then passes through a convolutional layer, which uses motif detectors to scan the OHE matrix and extract features. The output array is processed by a rectification layer that applies an absolute value operation to the extracted features. Next, the pooling layer enhances the robustness of these features and reduces overfitting. The features are then fed into a fully connected (FC) layer, where a probability score is calculated, representing the likelihood of protein binding. The training pipeline is designed to automatically adjust these parameters, minimizing the need for manual intervention and improving accuracy.

**ii) DESSO (DEep Sequence and Shape mOtif)**: This framework, implemented in Python, accepts both the OHE matrix of DNA sequences and DNA shape features to identify sequence and shape motifs **[115]**. Its multi-layer CNN architecture enables the extraction of more complex motif patterns compared to existing methods (**Figure 5C**). The first CNN layer contains multiple convolutional filters, or motif detectors, which detect low-level features from ChIP-seq peaks. High-valued features are then extracted using a max pooling layer followed by a FC layer. To reduce bias, a large set of background sequences was selected from the human genome, considering factors like GC content, no overlaps with positive peaks, and promoter and repeat overlaps. The framework uses these background sequences and the query sequence matrix to compute p-values for TFBR candidates using a binomial distribution. The optimal motif instances are identified based on the lowest p-values. The rectified linear unit (ReLU) activation function, known for better convergence and preventing the vanishing gradient problem, is employed in this model.

**iii) TSPTFBS (trans-species prediction of transcription factor binding sites)** is another CNN-based approach (**Figure 5C**) designed to identify TFBRs. It used DAP-seq experimental dataset for 265 TFs in *A. thaliana*, and has been implemented in Python programming language. The architecture contains one convolution, ReLU, max pooling, and three fully connected layers, which can identify the TF-DNA binding intensity for a given TF across any particular DNA sequence. It takes input in the form of an OHE matrix of the input sequence provided. In the initial phase of training, a CNN model was trained with the training set to learn the best parameters and hyper-parameters including different filter numbers. Finally, they also evaluated the trained models for the biological interpretability on the independent test set. This tool also works on transfer learning strategies to perform cross-species TFBRs identification.

**iv) SeqConv** is designed for analyzing ChIP-seq data from 104 TFs in *Zea mays*. Its architecture is adopted by the DeFine sequence-based deep learning model, optimized for identifying TF binding to specific DNA sequences **[116]**. SeqConv processes raw DNA sequences by generating their reverse complements to mimic the DNA double strand, as TFs recognize both strands at specific positions. These sequences are then converted into OHE, allowing the CNN to simultaneously model both the forward sequence and its reverse complement. The forward and reverse-complement sequences are fed into convolution layers that share the same filters, ensuring consistent feature extraction from both strands. SeqConv applies two pairs of max pooling and average pooling layers to the resulting features, which are then combined into a single vector. This combined vector is passed through a batch normalization layer before entering the fully connected layer. The final output is generated by a hidden layer with 64 nodes, leading to the output layer with a sigmoid activation function, which predicts the ChIP-seq binding probability.

**v) BPNet** models the relationship between cis-regulatory sequences and TF binding profiles at a base-pair resolution. The architecture created to model continuous base-resolution binding profiles as a function of DNA sequence, which was used to model ChIP-nexus/exo and ChIP-seq resolution binding profiles. It employed nine dilated convolutional layers with residual skip connections and exponential dilation to learn progressively complex predictive patterns within a 1-kb receptive field. They used only fully convolutional model without any pooling layers, especially maximum pooling layers. Lastly, it used a multiscale loss function to assess overall read counts using a mean-squared error loss and profile shape predictions using a multinomial negative log-likelihood loss. To extract the sequence features that were important for TF binding from the trained BPNet model, they extended DeepLIFT to quantify the contribution of each base within an input sequence. They further used TF-MoDISco to discover and summarize recurring sequence patterns into consolidated motifs from sequences of all bound regions from their associated base-resolution contribution scores.

**vi) AgentBind (DeepSEA (deep learning–based sequence analyzer))** employs a CNN framework that operates entirely based on sequences developed for genomic sequence by learning large-scale chromatin-profiling data, including TF binding, DNase I sensitivity and histone-mark profiles. The model includes three major characteristics: I) Integrating sequence information from a wide sequence context, II) Learning sequence code at multiple spatial scales using a hierarchical architecture, and III) Multitask joint learning of diverse chromatin factors that share predictive features. The model architecture includes three convolutional layers, two pooling layers, and one FC layer. The convolutional layers use kernels with sizes of 320, 480, and 960, respectively. The input sequence undergoes alternating convolution and pooling operations. The high-level convolutional layers capture broader spatial contexts, while the low-level layers extract more detailed feature information. The final FC layer integrates these sequence features across the entire sequence.

#### 2) Complex CNNs-based TFBRs identification software tools

In order to solve the problem of the vanishing/exploding gradient, complex CNN architectures are designed for DL models (**Figure 5D&E**). Some examples of complex CNN architectures are: ResNet (Residual Network) and DenseNet (Densely Connected Convolutional Network). ResNet utilizes skip connections, forming residual blocks, by connecting layer activations to subsequent levels, thus facilitating the stacking of these blocks to compose the network architecture. DenseNet, characterized by its tightly connected architecture, incorporates feature sharing and arbitrary interlayer connections within its convolutional neural network structure. Below, we discuss some tools that incorporate such architectures to identify TFBRs:

**i) MAResNet** is a method that integrates bottom-up and top-down attention mechanisms with a feed-forward network called ResNet, implemented in Python. This multi-scale attention mechanism is designed to extract rich and representative sequence features for identifying relevant patterns and motifs in DNA sequences (**Figure 5D**). The architecture employs two key branches: the soft mask branch and the trunk branch. The soft mask branch, which includes a feed-forward sweep and a top-down feedback step, rapidly collects and integrates global information, functioning as a feature selector that enhances important features to capture relevant patterns and motifs while suppressing noise. The trunk branch is used for feature extraction via CNN layers. MAResNet’s architecture processes DNA sequences using a binary OHE matrix. The sequence data is passed through convolutional, batch normalization, and ReLU layers, followed by three attention modules distributed across residual blocks (stages 1, 2, and 3). These attention modules adaptively generate attention-aware features, with the first stage employing multi-scale attention using convolution kernels of various sizes (3, 5, and 9), and later stages using 64 convolution kernels with a size of 13. The average pooling layer further reduce the size of the feature map. Finally, the dropout and the softmax function were used at the end of the network model

**ii) TSPTFBS 2.0:** The updated version of TSPTFBS, utilizing DenseNet, was constructed on a large-scale dataset of 389 plant TFs **(Figure 5E)**. DenseNet, which builds on the ResNet concept, connects each layer to all preceding layers, enhancing feature map reuse and reducing layer interdependence. This architecture effectively reduces the vanishing gradient problem, which is challenging in deep networks. TSPTFBS 2.0’s architecture includes an average pooling layer, four DenseBlocks, usually convolutional layers, three TransitionLayers, and two convolution layers (64 filters, kernel size = 3). Each DenseBlock consists of multiple layers with deep connections between them, ensuring that every layer is linked to all previous layers. The DenseBlocks maintain constant feature map sizes, allowing for effective concatenation, while TransitionLayers use 1×1 convolution and 2×2 pooling to downsample feature maps and reduce spatial dimensions. The DenseLayers within DenseBlocks are arranged with specific layer counts, “*Li”*, (Li = 6, 12, 24, 16). The final output from the last DenseBlock is flattened and passed through a fully connected layer to produce the final output value.

#### 3) RNN-based TFBRs identification software tools

RNN is a form of artificial neural network (ANN) intended to analyze data sequences (**Figure 5F**). Unlike classic feed-forward neural networks, which are well-suited for fixed-size input data, RNNs are specialized for sequences of various lengths. This above-described characteristic makes RNN particularly helpful for time-series data, natural language processing (NLP), speech recognition, as well as image and video analysis. RNN is meant to use sequential information from input data by using cyclic connections between building blocks such as the perceptron, long short-term memory (LSTM), or Gated recurrent unit (GRU). Some of the tools developed based on this philosophy to detect the regulatory sites are:

**i) KEGRU** is a Python-based implementation of an RNN algorithm, designed for identifying TFBRs using a Bidirectional Gated Recurrent Unit (BiGRU) network with *k*-mer embedding (**Figure 5F**). This model represents input DNA sequences as low-dimensional vectors through word embedding, capturing the distributional nature of *k*-mers. In KEGRU, *k*-mer sequences are initially extracted from DNA with specific lengths and stride windows. These sequences are converted into *d-*dimensional vectors using word2vec, where “*d*” represents the vector space dimensionality. The BiGRU network then processes these embeddings to learn and extract features from *k*-mers and identify TFBRs. The network incorporates *k*-mer embedding representation to refine its internal state and recognize complex relationships. The final output is generated by a dense layer with a sigmoid activation function, and performance is evaluated by comparing predictions with actual target labels using a loss function. KEGRU involves two categories of hyperparameters: model-related and data-related, with 12 parameter settings derived from different combinations of optimizers and GRU units. The tool operates in two main steps: model training and statistical analysis of the results, which are categorized by cell type, hyperparameter settings, and test indicators during data statistical analysis.

#### 4) Transformer-based TFBRs identification software tools

Traditionally, NLP tasks, such as machine translation and language modeling, relied on RNNs and CNNs to process text sequences. However, these models come with significant limitations, including challenges in handling long-term dependencies and high computational costs. The emergence of Transformer-based models addresses these constraints by leveraging self-attention mechanisms, enabling the model to focus on different parts of the input sequences, even when they are widely separated. While Transformers can be considered a form of RNN architecture, their notable improvement in performance, attributed to self-attention with multiple learning heads, elevates them to a league of their own in the realm of DL architectures. Bidirectional Encoder Representations from Transformers (BERT) is a pre-trained language model that utilizes the Transformer architecture to produce high-quality text representations. It represents a methodology for learning language representations using a transformer, particularly the encoder component.

**i) DNABERT** is a BERT-based tool designed for genomic sequence analysis using a Transformer architecture **(Figure 5G)**. It serves as a bidirectional encoder specifically developed to capture global, transferable information from DNA sequences, accounting for both upstream and downstream nucleotide contexts over extended ranges. Implemented in Python, the pre-trained DNABERT model learned basic syntax and semantics of DNA via self-supervision and fine-tuning by using task-specific labeled data, utilizing large datasets to build general-purpose representations before adapting to task-specific data with minimal architectural changes. Unlike standard BERT, DNABERT uses *k*-mer representation instead of individual nucleotide tokens, integrating bases into *k*-mers to provide richer contextual information. These *k*-mer tokens are embedded into numerical vectors, forming a matrix representation. During pre-training, DNABERT applies masked language modeling by masking *k*-length stretches of *k*-mers within DNA sequences to simulate regions where TF binding sites may occur. The model identifies the masked *k*-mer based on its surrounding context, enhancing understanding of token interconnections. DNABERT processes input through 12 Transformer blocks following an embedding layer, using outputs from the final hidden states for sentence-level classification and masked token outputs for token-level classification. DNABERT’s applicability extends to other mammalian species, as demonstrated by fine-tuning pre-trained models on the human genome and testing their performance on 78 mouse ENCODE ChIP-seq datasets **[117]**.

#### 5) Hybrid DL-based TFBRs identification software tools

Hybrid neural network model is a combination of various DL architectures as shown in **Figure 5H&I**. Most of the existing ones have used CNNs and RNNs. In genomics, it is a standard practice to combine CNNs with RNNs, leveraging CNNs to extract features from DNA or protein sequences and RNNs to capture sequential relationships in the data. This hybrid approach finds application in diverse tasks such as protein interaction prediction, genomic sequence categorization, and DNA-protein interaction identification. In the following section, we delve into a range of hybrid DL-based TFBRs identification software tools:

**i) WSCNNLSTM** proposed a combinatorial model using multiple-instance learning (MIL) with a hybrid DL network (**Figure 5H**), combining convolutional and RNNs to account for the spatial and sequential characteristics of DNA sequences. The model aims to learn the weakly supervised information of genomic sequences, as bound DNA sequences may have multiple TFBRs. The proposed framework may be considered in two sections: in the first one, this tool divides sequences into several overlapping instances using a sliding window, and then uses *k*-mer encoding to transform all instances into image-like inputs of high-order dependencies. In the second one, to capture the forward and backward local dependencies between motif features, they added a bi-directional recurrent layer after the convolutional layer in the weakly supervised framework. The convolutional layer captures motif features, while the recurrent layer captures long-term dependencies between these features. The convolutional layer scans for motifs using a fixed number of kernels, the max-pooling layer retains local best values, and the dropout layer prevents overfitting by randomly zeroing outputs. The bidirectional recurrent layer, composed of LSTM units, captures forward and backward long-term dependencies. Finally, the softmax layer computes a probability distribution over two labels representing bound or non-bound sequences.

**ii) deepRAM:** The purpose behind developing the deepRAM tool was to explore various combinations of DL algorithms, such as CNN/RNN based models (depicted in **Figure 5C, F&H**), to identify DNA/RNA-binding specificity. Implemented in Python, the architecture of deepRAM involves representing input sequences in two ways: OHE or *k*-mer word representation computed using word2vec. The DL architecture in deepRAM includes CNN-only models, renowned for detecting spatial patterns and motifs. RNN-only models excel at capturing sequential dependencies within sequences. Hybrid CNN-RNN models combine the strengths of both by capturing sequence-specific motifs and their context. CNNs are applied to biological sequence data by integrating one-dimensional convolution. RNN-only models utilize three types of RNN units: LSTM units, GRU units, and Bi-GRU. In all three types of models, the final module consists of one or two fully connected layers that integrate information from the entire sequence. This is followed by a sigmoid layer at the output of neural networks, calculating the probability that the input sequence contains a DNA/RNA-binding site.

**iii) AgentBind (DanQ):** DanQ is a hybrid framework combining convolutional and bi-directional long short-term memory (Bi-LSTM) recurrent neural networks to identify non-coding functions from sequences. In the DanQ model, the convolutional layer captures regulatory motifs, while the Bi-LSTM layer learns local dependencies between these motifs, aiming to establish a regulatory ‘grammar’ that enhances accuracy. The inclusion of the recurrent layer after max pooling helps capture the regulatory grammar influenced by physical constraints that affect the spatial arrangements and frequencies of motif combinations, which is particularly relevant for tissue-specific elements like enhancers. After the Bi-LSTM layer, DanQ features a dense layer with ReLU followed by a multi-task sigmoid output layer, similar to the DeepSEA model. Lastly, they used Grad-CAM **[118]**, a post-analysis technique for neural networks, to calculate importance scores for each nucleotide in the contextual regions and to identify sequence features that identify transcription factor binding regions.

**iv) PlantBind** is an attention-based multi-label neural network method designed for TFBRs identification. This integrated model comprises two separate CNN networks and an LSTM-Attention block (depicted in **Figure 5H**), which extract and learn relevant information. Implemented in Python, PlantBind supports simultaneous analyses for multiple TFs, combining data processing, embedding, feature extraction, and multi-label output modules. The initial module utilizes OHE to transform DNA sequences for handling by CNN, along with 14 DNA shape features for each input sequence. This yields two heterogeneous data types: a sequence matrix and a shape matrix. Two CNN models embed these matrices into the same shape, enabling them to be combined into one embedding matrix. In the feature extraction module, the embedded representation undergoes processing by an LSTM layer to extract shared underlying features across all TFs. The attention mechanism then allows the model to focus on relevant regions of the embedding vector for each specific TF. The multi-label module consists of two fully connected layers that generate multiple binding sites simultaneously. The framework is trained using cross-entropy loss enhanced by online hard example mining (OHEM) and focal loss (FL). In summary, PlantBind learns to associate sequence context with TF types and utilizes the attention mechanism to extract sequence content contributing most to identification. Its integrated multi-label framework enables learning of underlying associations between different TFs.

**(v) DeepSTF**, a novel multi-module hybrid DL architecture that combines CNN, enhanced transformer encoder structure, and Bi-LSTM (**Figure 5I**). It uses a combination of DNA sequence and shape for TFBRs prediction. They used five DNA shape features which are: helix twist (HelT), minor groove width (MGW), propeller twist (ProT), rolling (Roll), and minor groove electrostatic potential (EP). DNA shape information is incorporated into the DeepSTF model, which improves its performance by increasing the accuracy of TFBRs. The model extracts rich DNA shape features and integrates them with higher-order sequence characteristics by utilizing the enhanced transformer encoder structure, CNNs, and Bi-LSTM. The model is able to learn key characteristics and capture various dependencies as a result to this integration, which eventually raises the accuracy rate of TFBRs. Furthermore, the enhanced transformer encoder structure’s multi-headed attention mechanism helps the model produce a higher-level representation of the input data that is more thorough, comprehensible, and efficient at identifying important underlying features while suppressing unnecessary ones.

**(vi) PTFSpot** represents a cutting-edge deep-learning approach based on transformers, designed to learn from the structures of TFs and the co-variability of their binding regions (**Figure 5I**). This method aims to create a universal model for TF-DNA interactions, effectively detecting TFBR without the limitations of TF- and species-specific models. Leveraging binding data for 436 TFs and their corresponding 3D structural information, the algorithm constructs a Transformer-DenseNet based universal model to learn several hidden features. PTFSpot uses the interdependent variability of TF structures and binding regions to determine the most likely binding locations for any given TF, regardless of other TF or species specific models. Its universal TF-DNA interaction model enables it to identify probable binding regions for even unknown TFs and species.

### Performance evaluation strategies for TFBRs identification software tools

In this study, to evaluate the performance of all the considered tools for TFBRs identification, we compared all of these 19 tools across three different datasets for the comparative performance assessment as described above (Datasets “A”, “B”, and “C”). The default parameters were considered for Dataset “A”, as all of these software tools had already optimized their hyper-parameters during the training and model building processes for the classification task. Classifier evaluation relies on analytical reasoning, particularly in distinguishing between missed true instances (false negatives (FN)) and erroneously identified non-binding regions (false positives (FP)). To assess this, we consider the count of true binding regions (true positives (TP)) and correctly identified non-TFBR (true negatives (TN)). The ratio between the number of correctly classified samples, i.e., TP and TN, and the overall number of samples, termed as accuracy, a key metric in this evaluation process. The accuracy for the positive class can be seen as the fraction of positive instances that are correctly recovered by the model, also called the true positive rate (TPR) or sensitivity (Sn). The specificity (Sp) metric is used for evaluating a model’s ability to identify the TN from total negatives. Besides these, the precision metric quantifies the accuracy of positive predictions made by a model. Precision is defined as the ratio of true positive predictions to the total positive predictions made by the model. The recall is determined as the ratio of the number of positive samples accurately classified as positive to the total number of positive samples. The F1-score combines the precision and recall of a classifier into a single metric by taking their harmonic mean. The performance of the models were also assessed by computing the area under the receiver operating characteristic (AUROC) curve (AUC). A ROC curve depicts the trade-off between the true positive rate (TPR) and false positive rate (FPR) at various decision thresholds. Finally, six different metrics for performance were considered (Sn, Sp, F1-Score, Accuracy, Matthew’s correlation coefficient (MCC), and AUC). MCC is a reliable statistical measure which produces a high score only if the model obtains good results in all the four confusion matrix categories (TP, TN, FN, and FP). We calculated Acc, Sp, Sn, F1-score, AUC and MCC, as given in the following equations:

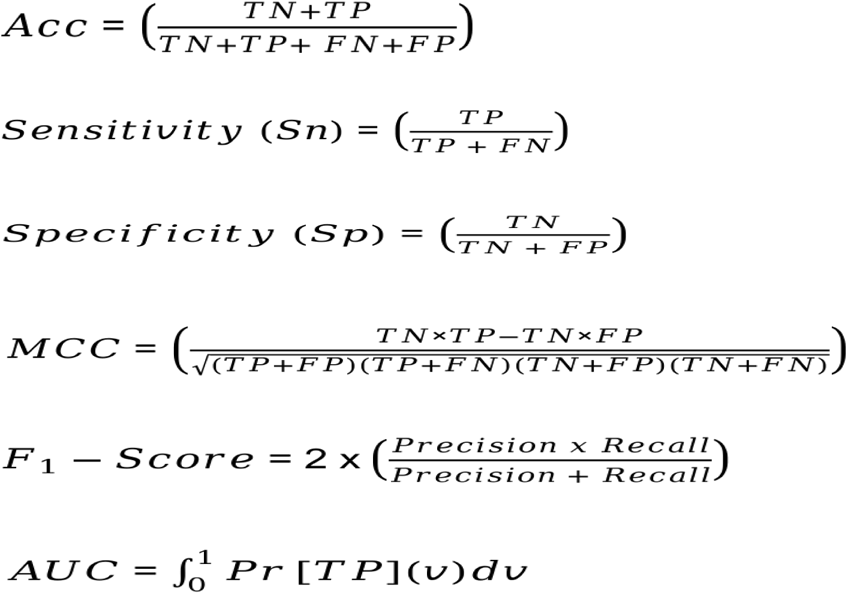

Where:

TP = True Positives, TN = True Negatives, FP = False Positives, FN = False Negatives, Acc = Accuracy, and AUC = Area Under Curve.

### Dataset for transcription factors binding motif identification

In our study on TFs binding motifs, we focused on comparing traditional motif identification techniques with DL methods. This analysis required a robust dataset, which we sourced from DAP-seq peak data. The curated and processed plant DAP-seq peak datasets for 10 specific TFs have been taken from Dataset “B”: AT2G28810, AT3G24120, AT5G02460, AT5G62940, AT5G66940, ATHB25, ATHB40, ATHB5, CEJ1, and MYB77. These TFs were selected based on the availability of high-quality and highest number of peaks, ranging from 9151 to 11,990, so that we could generalize the applicability of the motif finding solution of the software tools **Supplementary Table S2 Sheet 1**. By comparing these methods on the curated dataset, we aimed to assess their relative performance and identify scenarios which of the traditional or DL methods significantly perform. Hence, we have conducted a comprehensive analysis of TF binding motifs, comparing traditional motif identification, included HOMER **[119]**, CMF **[120,121]**, MEME **[122]**, MEME-ChIP **[122]**, and iFORM **[123]** and four of the most advanced tools DL based methods, deepRAM, DESSO, TF-MoDISco, and PTF-Vāc **[124]**.

### Characterization of software tools for transcription factor motif identification

To effectively identify transcription factor binding motifs, a variety of software tools have been developed, which include various traditional as well deep learning algorithms. These tools employ diverse algorithms and methodologies to analyze DNA sequences and pinpoint specific motifs associated with transcription factor binding. In this comparative assessment, we examine and characterize the most prominent software tools available for transcription factor motif identification. We explore their underlying algorithms, strengths, limitations, and the specific contexts in which they perform best. By providing a comprehensive overview, this characterization aims to guide researchers in selecting the most appropriate tools for their specific needs in motif discovery and analysis.

#### 1) Traditional motif discovery software tools

Most of these traditional approaches may be considered as unsupervised approaches as they work from labeled data and classification problem, while mainly performing clustering to raise a TF specific profile directly from binding data, and then doing scanning against it. The motif discovery algorithms are classified into four classes of enumerative, probability, nature inspired and combinatorial ones and each one has many sub-classes. Of these classes, probability approach overcomes many weak points of the enumerative approach like speed, dealing with long motifs and big data, numbers of required parameters and degenerated positions, and can find weak motifs. Some of the methods based on probability based algorithms are discussed below.

**i) HOMER:** it uses a motif discovery algorithm designed for regulatory element analysis in genomics, focusing solely on DNA rather than proteins. This algorithm employs differential motif discovery, analyzing two sets of sequences to pinpoint regulatory elements specifically enriched in one set compared to the other. It uses zero or one occurrence per sequence scoring (ZOOPS) along with hyper-geometric or binomial enrichment calculations to determine motif enrichment. Additionally, HOMER attempts to account for sequence bias within the dataset. Originally designed for ChIP-seq and promoter analysis, this versatile tool can be applied to various nucleic acid motif finding tasks.

**ii) CMF: C**ontrast **m**otif **f**inder (CMF) is a *de novo* motif discovery tool that find differentially enriched motifs in two sets of sequences and provide a non-discretized estimation of PWMs which are corrected for false positives. CMF generates two sets of motifs, one predominantly found in experimental dataset and the other predominantly found in control sequences. For each identified motif, CMF provides a list of identified binding sites within both input sequence sets and calculates a likelihood ratio (LR) score for each identified site.

**iii) MEME: Multiple EM for Motif Elicitation (MEME)** is a web-based service available on MEME suite as well can be downloaded and installed locally. MEME is a tool for *de novo* motif discovery, designed to identify un-gapped motifs in unaligned DNA or protein sequences. MEME algorithm builds upon the expectation maximization (EM) algorithm. It chooses starting points from all sub-sequences within the input dataset. It eliminates the assumption that the shared motif appears in every sequence. Once a motif is discovered, MEME removes its occurrences and continues searching for additional shared motifs in the dataset.

**iv) MEME-ChIP:** It is a web-based service developed for analyzing various genomic datasets such as ChIP-seq, CLIP-seq, and other selected genomic regions, accessible through the MEME suite can be downloaded and installed locally. Its functionalities encompass *de novo* motif discovery, motif enrichment analysis, motif location analysis, and motif clustering, offering a comprehensive understanding of enriched DNA or RNA motifs within input sequences. Notably, MEME-ChIP employs two distinct approaches for *de novo* motif discovery: a weight matrix-based method for precise identification and a word-based method for heightened sensitivity. This dual strategy utilized by MEME-ChIP involves two distinct motif discovery algorithms, namely MEME and discriminative regular expression motif elicitation (DREME) **[122]**, to uncover sequence motifs within a given set of nucleotide sequences. Following motif discovery, it employs a motif enrichment analysis algorithm known as central motif enrichment analysis (CentriMo) **[122]** to identify the enrichment of known functional motifs associated with TFs or RNA-binding proteins (RBPs) in the sequences. To facilitate result interpretation, MEME-ChIP further applies a clustering algorithm, grouping the discovered and enriched motifs based on their similarity to each other. This comprehensive approach enables researchers to gain valuable insights into the regulatory elements present in the analyzed sequences.

**v) iFORM:** Find Occurrence of Regulatory Motifs (iFORM) is a tool for scanning DNA sequences with TF motifs represented as PWMs. It integrates five traditional motif discovery programs through Fisher’s combined probability test. These algorithms involve motif instances identified by five classical algorithms, namely FIMO **[122]**, Consensus **[125]**, STORM **[126]**, RSAT **[127]**, and HOMER, utilizing Fisher’s method as these algorithms were selected for their widespread usage in various applications and the broad applicability of their scoring methods as motif scanners.

#### 2) Deep learning motif discovery software tools

Deep learning motif discovery tools leverage neural network architectures to learn complex patterns directly from the data. These tools, such as deepRAM, DESSO, TF-MoDISco, and PTF-Vāc can capture intricate relationships within sequences without relying heavily on predefined motifs. Here, it should be pointed out that PTF-Vāc utilized an *ab-initio* motif discovery by deep co-learning on variability in binding regions and TF structure. They are often more flexible in capturing subtle motifs and can potentially discover novel motifs that may not be well-characterized by traditional methods. These motif finding tools are briefly discussed below.

**i) deepRAM:** deepRAM uses convolutional filters for feature extraction in motif identification, emphasizing local pattern recognition. Despite the effectiveness, deepRAM mainly focus too much on individual filters, leading to redundant motif sequences and a limited ability to capture distributed motif representations.

**ii) DESSO:** This approach uses convolutional filters to identify low-level features, followed by max pooling and fully connected layers for high-level feature extraction. Motif detectors learn patterns from DNA sequences and background sequences, which are selected to eliminate biases. However, its reliance on a binomial distribution for p-value calculations can be a limitation, potentially affecting the accuracy in representing motif occurrences, especially in non-uniform distributions or cases where there are dependencies among motif instances.

**iii) TF-MoDISco:** This method uses contribution scores to assess positional importance, providing insights into the significance of individual bases. It identifies high-importance windows known as “*seqlets*”, which are then clustered into motifs after post-processing. While per-base importance scores help incorporate information from all network neurons, the model may assign disproportionately high importance to specific positions based on prevalent motifs in the training data. This can lead to biases and potentially overlook less frequent but biologically relevant motifs.

**iv) PTF-Vāc:** PTF-Vāc is the first of its kind *ab-initio* motif discovery tool, which claims to be not sensitive to volume of input data to report a binding region specif motif. It was developed on Deep Encoders-Decoders system, and is a universal model of deep co-learning that accounts for variability in binding sites and TF structure, making it entirely free from limitations of data volume species and TF specific models. Its capability extends to accurately detecting the binding motifs of unseen TF families and species, while also enabling the definition of credible motifs from its TFBR report. PTF-Vāc utilizes the purely refined informative regions obtained from PTFSpot to transform the extended binding regions suggested by PTFSpot into key elements, specifically focusing on potential binding sequence/motif components, such as potential binding sequence/motif components. It additionally applies an importance score using Grad-CAM to parallely substantiate the identified important TFBS/Motif candidate. It has effectively decoupled the challenge of motif identification from that of discovering binding regions, particularly within plant-specific systems.

## Results

### Performance benchmarking using transcription factor ChIP-seq data

Evaluating peak calling software tools is crucial for ensuring the accuracy and reliability of data analysis in genomics research. Accurate identification of genomic regions where TFs bind to DNA is essential for understanding gene regulation and other biological processes. TFBR models and subsequently motif discovery are built using the same. Therefore, any noise and wrong data selection at this level is supposed to influence all the computational models for downstream analysis. Therefore, identifying peaks is one of the initial steps in analyzing these experimental ChIP-seq data. Peak calling consists of two sub-problems: identifying candidate peaks and testing candidate peaks for statistical significance. Therefore, we analyzed different peak calling tools which produce varying results, and thorough evaluation helps one may choose the most reliable software tools for their specific needs, ensuring reproducibility and consistency in peak finding. Moreover, such evaluation drives the development of better algorithms, minimizes errors, and sets benchmarks, thereby advancing computational tools and supporting robust TFBRs’ and motifs’ discoveries. After enlisting 28 peak callers (**Supplementary Material S1 Table 2**) and execution of these software tools, we were capable to pick only six peak calling methods, which utilized various algorithms to identify candidate peaks from our input ChIP-seq datasets. The reason to run only six software tools was that the remaining ones were not maintained properly and are out of usage to the users in public domain.

Despite variations in algorithm, the extent to which these methodological variations interpret into their significant performance differences remains unclear. Therefore, benchmarking peak calling tools is challenging due to the absence of a comprehensive list of true positive genomic regions linked to experimental conditions. To address this, we selected five publicly available ChIP-seq datasets for five TFs, namely MYB62, bZIP9, WRKY94, HB33, and EREB147, with a control or negative dataset, which is used to distinguish true signal peaks from background noise in ChIP-seq experiments. After processing of the data, we used this data as an input for the considered tools and compared the final peak calling results on the basis of: 1) the total number of identified peaks after peak calling, 2) the total number of identified peaks overlapped with experimental dataset, 3) the true positive and false positive rates, 4) the overlapping between identified peaks among these six software tools (**Figure 6**), and 5) the estimated reproducible and stringent peaks obtained from IDR to measure reproducibility between replicates or different peak finders which further provides a rigorous assessment of peak consistency.

**Figure 6:**
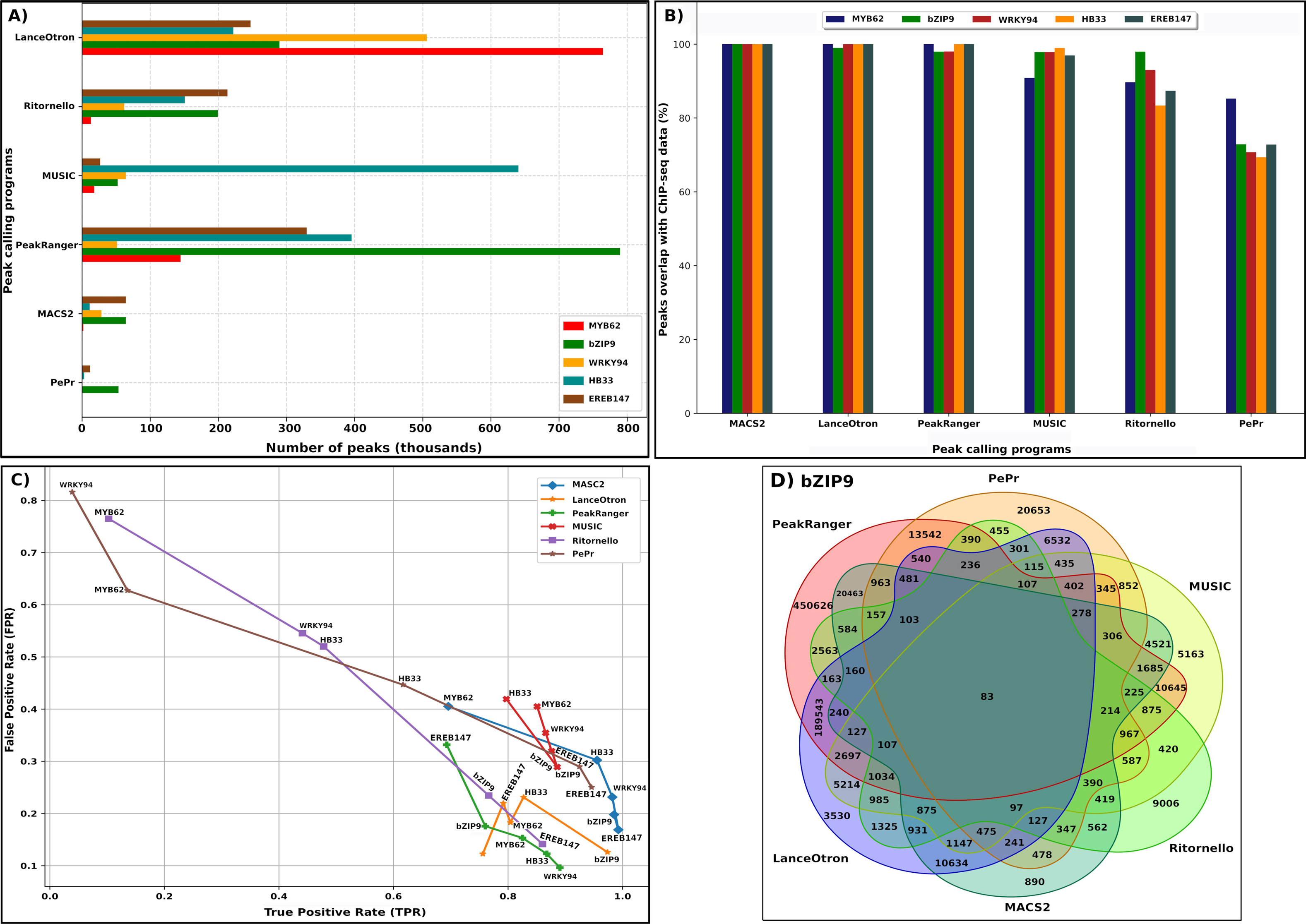
This figure depicts the diversity in peak identification across various programs applied to the TF binding experimental dataset. **(A)** It delineates the total peaks detected, **(B)** the percentage of overlapping peaks with the experimental data, **(C)** true positive and false positive rates, and **(D)** the overlap between six peak calling tools specifically for bZIP9, offering insights into their performance and reliability

For the first analysis, their are notable differences in performance among the examined algorithms across these five TF datasets, as illustrated in **Figure 6**. It can be clearly visualized from **Figure 6A** that LanceOtron emerged as the top performer, consistently identifying the correct peaks, exceeding two lakhs across all the five TFs. PeakRanger ranked second, detecting peaks exceeding one lakh for four TFs except the WRKY94, for which 51,121 number of peaks were recovered. MUSIC also exhibited strong performance, particularly for HB33, identifying peaks exceeding six lakh. Conversely, PePr and MACS2 consistently identified fewer than sixty thousand peaks for all the TFs. The differences in performance can be attributed to variances in default stringency levels and each tool’s capability to precisely detect peaks. However, the total number of peaks after peak calling may not solely guarantee the reliability of obtaining all the true binding events of the peaks for a particular TF. Therefore, we further overlapped the output of the software tools with that of the experimental dataset. As shown in **Figure 6B**, it was found that the output peaks identified by MACS2 for all the five TFs was 100% overlapping with the experimental datasets. On the other hand, for LanceOtron, the output peaks for four TFs scored 100% except for bZIP62. Similarity, for PeakRanger, 100% peaks for MYB62, HB33 and EREB147 overlapped with their ChIP-seq dataset. However, peaks retrieved by PePr for all the five TFs were less then 82% overlapping with the experimental dataset. This analysis suggests that the peaks generated by these software tools may be influenced by the statistical models to assess the peaks, some of which were not capable to correctly retrieve possible true binding peaks and rather captured more of the negative data signals from background control or negative data samples.

Therefore, to obtain all the true binding events from the peaks output with less noise and bias, which could provide more credibility to the generated experimental data and their subsequent downstream analysis for TFBRs and motif identification, we also evaluated these software tools on the basis of true positive rate (TPR) and false positive rate (FPR). We found MACS2, LanceOtron, and PeakRanger as the most credible ones. As shown in **Figure 6C**, it was found that MACS2 showed the TPR values greater than 95% for four TFs, namely bZIP9, WRKY94, HB33, and EREB147, respectively, except for MYB62, for which TPR value of 69.56% was reported. In contrary to overall TPR analysis, MACS2 retrieved significant peaks with the FPR values between 16% to 40% for all the five TFs. On the other hand, both LanceOtron and PeakRanger showed TPR values greater than 80% for three TFs. Here, LanceOtron obtained 80.42%, 97.36%, and 82.70% of TPR for MYB62, bZIP9, and HB33, respectively, while PeakRanger achieved 82.52%, 89.05%, and 86.75% of TFR values for MYB62, WRKY94, and HB33, respectively. LanceOtron showed FPR values between 12% to 23%, while FPR values for PeakRanger was between 9% to 33%. For other method like Ritrornello, which retrieved TPR values less then 76% for four TFs (MYB62, WRKY94, HB33, and bZIP9) except TPR value of 86% for EREB147, while the FPR values ranged from 14.16% (EREB147) to 76.5% (MYB62). Similarly, for PePr, which output the peaks with TPR values less then 61% for three TFs, namely WRKY94 (3.91%), MYB62 (13.56%), and HB33 (61.65%), while highest TPR values for bZIP9 (92.46%) and EREB147 (94.58%). However, PePr gave the peak signals with FPR values ranging from 25.02% (EREB147) to 81.60% (WRKY94).

Further, from the true binding regions obtained after peak calling, we also found the number of unique and overlapped peaks between all the true peaks obtained from these software tools. For the sake visualized peaks for bZIP9 (shown in **Figure 6D**). Here, PeakRanger and PePr have highest number of unique peaks with a total number of 4,50,626 and 20,653, respectively. Also the greatest number of overlapped peaks were between LanceOtron and PeakRanger, with a value of 1,89,543. For the common peaks between the three pairs of MUSIC and PeakRanger, LanceOtron and MACS2, PeakRanger and MACS2, a total of 10,645, 10,634, and 20,463, respectively. The maximum number of overlapped peaks (1,685) were obtained for MACS2, PeakRanger and MUSIC. However, minimum number of overlapped peaks (115) were reported between the software tools which were intersected with Ritrornello, LanceOtron, MUSIC, and PePr (**Figure 6D)**. This was the reason that we got only 83 peaks common between all these six software tools. Further details are given in **Supplementary Material S2 Figure 1**.

An ENCODE analysis pipeline had recommended using Irreproducible Discovery Rate (IDR) framework to reduce the inherent noise in ChIP-seq experimental data **[95]**. In our IDR analysis of peak reproducibility across four compatible software tools, the results revealed varying levels of peak consistency. Two of the studied software tools (PePr and MUSIC) generated output which was not compatible for IDR analysis. Specifically, MACS2 identified 70% of peaks as reproducible, while LanceOtron had 64.14% reproducible peaks. The third best tool, PeakRanger, yielded 59.72% of peaks as reproducible. Ritrornello showed a lower reproducibility rate of 46.87%. These percentages represent the proportion of peaks that were consistently detected and ranked across replicates, highlighting differences in peak reproducibility among the tools used. Also, in terms of reproducibility the top three best performing tools retained the same rank.

### Evaluation of software tools for TFBRs identification

Once the peak regions are identified as the binding regions, such data works as input to develop models of TFBRs. TFBR discovery is considered most important among all these three stages of computational challenges towards TF-DNA interaction modeling. Thus it is highly desirable to assess current status of this part of problem solving and propose the recommendations. To access the performance of the TFBRs identification software tools we summarized the entire TFBRs study from dataset collection to performance evaluation of all the 19 tools in **Figure 7**. By comparing their performance by means of metrics, such as sensitivity, specificity, and computational resources required, one can determine the most reliable and effective software tools for identifying TFBRs. We considered 38 tools for this part of the study ( **Supplementary Material S1 Table3)** and also faced many issues while trying to run these software tools. Some of the tools were not properly maintained by the providers while some were not upgraded as per the latest updates of OS and libraries, and hard were hard to fix for their issues. Hence, we were not able to run them on our dataset and had to discard them. We have tried to point out and fix these related issues and provided corrected scripts at out GitHub page (https://github.com/SCBB-LAB/Comparative-analysis-of-plant-TFBS-software), so that the users who are not an expert in this field could use them now. Some of the related issues and bugs are described below along with their solutions (**Supplementary Material S3)**.

**Figure 7:**
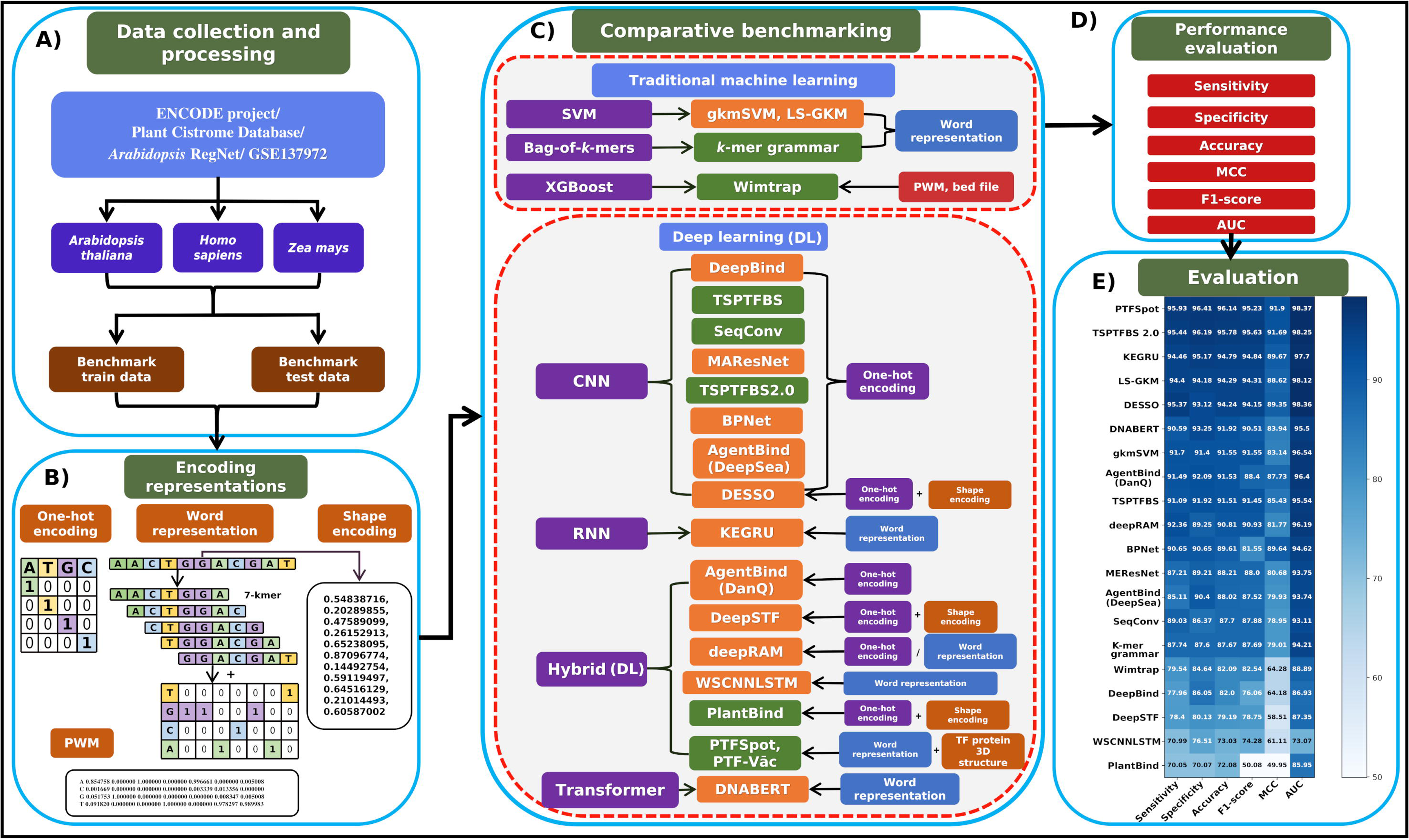
Overall layout of the present study. The image illustrates the outline of the entire evaluation flowchart to assess various models to identify TFBRs. **(A)** Illustrates data collection and processing to construct training and independent test datasets, **(B)** Shows the various encoding schemes used for input datasets used by the evaluated tools, **(C)** Description of the tools and their respective algorithms, **(D)** and **(E)** The evaluation criteria and the performance analyses parts.

### Bugs infest most of the existing tools

Throughout our evaluation of available software tools, we encountered numerous dependency issues and programming errors across C, C++, Python, and R implementations. These errors, ranging from “LogicError”, “KeyError”, “IndentationError”, “MemoryError”, “ImportError”, “AttributeError”, and many more, posed significant obstacles to the usability of these tools for biologists. **Table 2** provides a categorical overview of different types of errors faced during the extensive and exhaustive debugging of these software tools done in this study. To address these issues, we carefully debugged each tool, identifying and rectifying errors as they arose during execution detailed described in **Supplementary Material S3**. One prevalent issue was “ImportError” and “ModuleNotFoundError”, indicating difficulties locating or importing specific modules, observed in tools such as DeepBind, KEGRU, *k*-mer grammar, WSCNNLSTM, SeqConv, DESSO, and PlantBind. Similarly, “FileNotFoundError” emerged when accessing missing files or directories, affecting KEGRU, SeqConv, TSPTFBS, PlantBind, and TSPTFBS 2.0, necessitating appropriate file path adjustments. Runtime errors like “TypeError” and “KeyError” were also noted for tools like KEGRU, WSCNNLSTM, and SeqConv, which were resolved by data type conversions and dictionary key modifications. Additionally, “logicError” occurred in KEGRU, signaling flaws in code logic. For one of the software tools, AgentBind, we faced issues due to poorly defined data input formats and inadequate documentation on its GitHub repository. The input format description were not clearly outlined, leading to difficulties in integrating and processing data. To successfully execute BPNet, we encountered several issues due to its dependencies and the requirement for specific versions of packages. The tool’s functionality was heavily reliant on specific versions of various libraries and packages as listed in **Supplementary Material S4**. The stringent version requirements led to compatibility problems and difficulties in setting up the BPNet.

Furthermore, obtaining comprehensive performance metrics proved challenging for all software tools. Performance metrics are crucial for optimizing, selecting, and using models effectively. To address this, we developed dedicated functions or code segments to accurately calculate performance metrics, incorporated necessary calculations for evaluating model performance. The corrected scripts for each tool are now made available at https://github.com/SCBB-LAB/Comparative-analysis-of-plant-TFBS-software, which will now facilitate their use by the community.

### Hyper-parameter optimization and impact on performance

Optimal performance of DL networks relies on parameter and hyper-parameter tuning. While parameters are optimized automatically during learning, hyper-parameters, which are predefined settings, are not optimized automatically and significantly influence performance. The hyper-parameters include hidden units, batch size, optimizer choice, and learning rate, among others. In this study, all the 19 software tools (developed for animal and plant species) were analyzed on their default hyper-parameters on Dataset “A” providing consistent results across software tools. However, on Dataset “B” performance was evaluated after hyper-parameter optimization. Hence, we conducted hyper-parameter optimization specifically constructed for plant-specific TFBRs identification. During this process, models integrated knowledge from training instances and fine-tuned their parameters on validation and test datasets to minimize classification errors. Hyper-parameter adjustment during training aimed to optimize the model’s performance and enhance its ability to generalize. This optimization served as an objective assessment to monitor the model’s development and prevent overfitting **[128]**. Hyper-parameters, crucial for defining the model’s architecture, are typically set before finalizing an ML model. Two main categories of optimization techniques exist: manual search and automated search methods. Manual search involves manually testing different hyper-parameter settings, while automated approaches such as grid search **[129]**, random search algorithms **[130]**, and Bayesian hyper-parameter search **[131]** aim to overcome the limitations of manual search.

We performed hyperparameters optimizations which returned improved results for many of these tools while for some not much change was noticed. For instance, as listed in **Supplementary Material S5 Table 1**, DeepBind’s optimal hyper-parameters on Dataset “B” were {“Learning rate”: 0.001, “Batch size”: 128, “Learning steps”: 20000, “Dropout”: 0.50}, showing minimal performance variation from default settings as shown in **Figure 8A&B**. Similar results were observed for gkmSVM and LS-GKM, where the default parameter {“C”:1} yielded optimal performance. KEGRU achieved improved performance with the tuned hyper-parameters {“Batch size”: 200, “*k*mer length”: 5, “RNN unit”: 70, “Dropout”: 0.5, “Activation function”: softmax, “Optimizer”: Adam, “Loss”: Binary cross entropy} after optimization on Dataset “B” as can be observed from **Figure 8A&B**. Notably, deepRAM’s automatic hyper-parameter calibration enhanced average accuracy above 90% on Dataset “B”. For deepRAM, each of the 40 calibration sets, the best hyper-parameter settings was: {“Batch size”: 128, “Learning steps”: 5000, “Optimizer”: Adagrad, “Dropout”: 0.15, “Activation function”: ReLU}. As it can be seen from **Figure 8A&B**, these automatic hyper-parameter calibration upgraded the average accuracy on deepRAM of plant specific dataset (Dataset “B”) above 90%. For DESSO, an improvement of 9.13% accuracy was observed on Dataset “B”, with optimized hyper-parameters, including {“Activation function”: ReLU, “Hidden layers units”: 100, 50, “Dropout layer”: 0.4, “Activation function”: Sigmoid, “Optimizer”: SGD, “Batch size”: 64, “Epochs”: 50}. Similarly, TSPTFBS (plant-based) achieved an increase in accuracy on Dataset “B” with the optimized parameters. For TSPTFBS, we trained their model on Dataset “B” with optimized hyper-parameters. As shown in **Figure 8B**, we attained an increment in accuracy with a difference of 4.86%. DNABERT’s fine-tuned model exhibited improved accuracy compared to its default settings at: {“*k*mer length”: 6, “Learning rate”: 2e-4, “Epochs”: 5, “Batch size”: 1024, “Weight decay”: 0.001, “Optimizer”: AdamW, “Adam epsilon”: 1e-6, “beta1”: 0.9, “beta2”: 0.98}. We attained an increment in average accuracy by 2.38% from the default hyper-parameter. MAResNet also benefited from hyper-parameter adjustments: {“Optimizer”: SGD, “Weight decay”: 5e-4, “Momentum”: 0.9, dropout = 0.75, “Learning rate”: 0.004, “Batch size”: 256, “Loss”: Cross-entropy, “Activation function”: Softmax}, on Dataset “B”, providing a 2.6% accuracy improvement. For AgentBind, the default hyperparameters were: {booster: gbtree, alpha: 0.001, lambda: 10, eta: 0.05}. From the results, it’s evident that dataset refinement and hyper-parameter adjustment significantly impact model performance.

**Figure 8:**
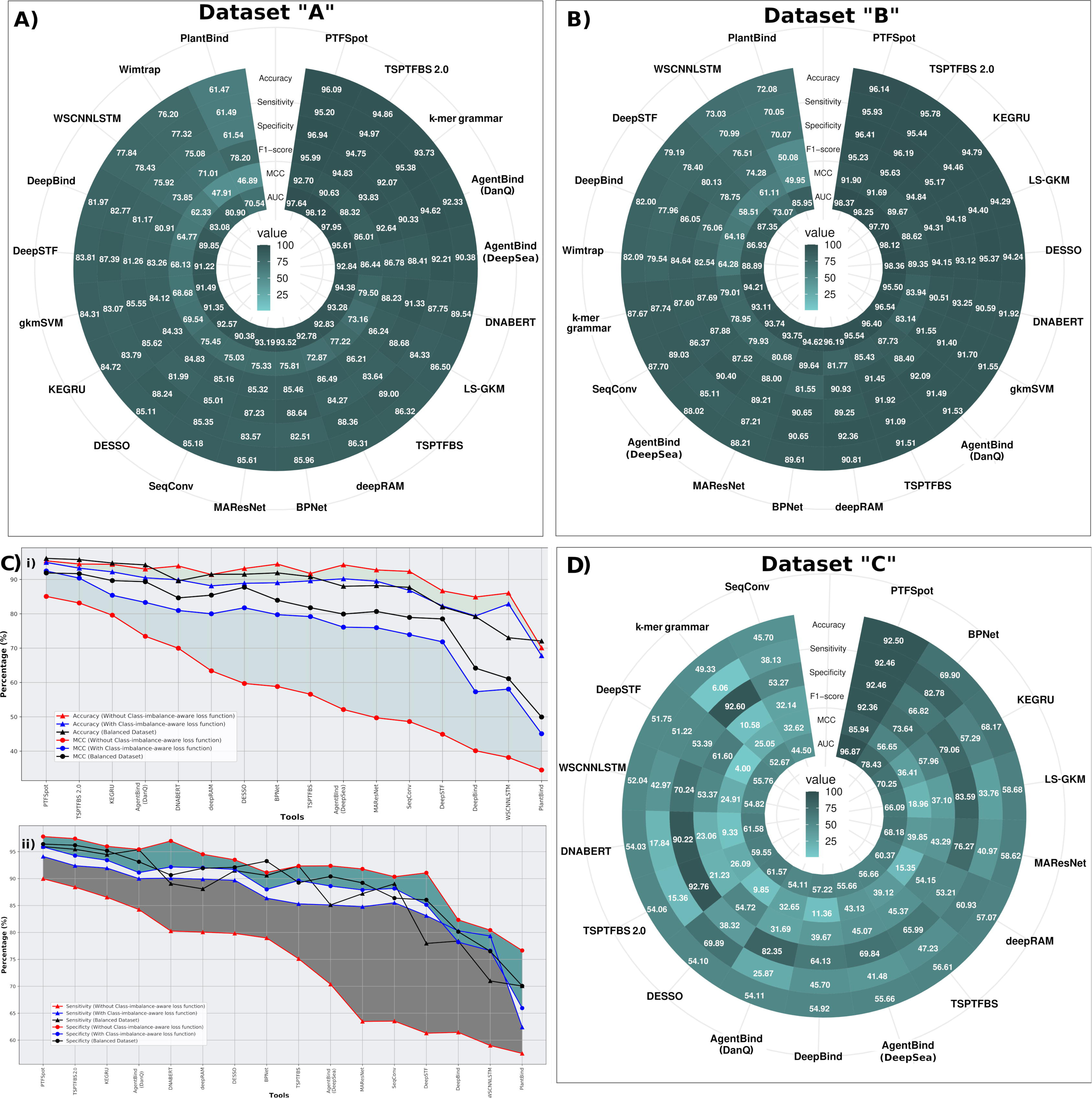
Performance and benchmarking of the traditional machine and deep learning software tools. Comparative evaluation with different combinations of test datasets. **(A)** Dataset “A”, **(B)** Dataset “B” used to train and test, and for comparative evaluation, **(C)** Impact of using class imbalanced datasets on models. **i)** The performance in terms of MCC and accuracy fall drastically when such datasets are used. Use of class-imbalance-aware loss function mitigates such issues, but yet remains lower performing than the models built with balanced datasets. **ii)** performance in terms of sensitivity and specificity. **(D)** Dataset “C”. For every such combinations, the performance metrics (sensitivity, specificity, F1-score, MCC and accuracy) are given in the form of circular heatplot. The **plot A** and **plot B** clearly indicate that PTFSpot consistently outperforms all the existing tools for all the performance metrics based on Dataset “A” and Dataset “B”. For Dataset “C”, as shown in **plot C,** is an independent test dataset used to compare the performance of all the listed tools for cross-species TFBRs identification. For Dataset “C”, again PTFSpot outperformed very well for the cross-species TFBRs identification. It clearly shows how poorly other software tools perform when applied in cross-species manner. None of the existing tools were found suitable for practical application for finding TFBRs across different species.

### Performance across different validated datasets

A comprehensive benchmarking study evaluated the performance of these 19 software tools based on ML and DL approaches across three datasets: “A”, “B”, and “C”. Datasets “A” and “B” were used for both model training and testing, while Dataset “C” specifically tested tools’ capabilities for cross-species TFBRs identification.

### Most of the reported performances were found correct

The analysis of model performance on Dataset “A” demonstrated the effectiveness of *in-silico* models in accurately identifying TFBRs, aligning with their stated claims **Supplementary Table S3**. On Dataset “A”, PTFSpot emerged as the top performer across all performance metrics, with an average accuracy of 96.09% and an average MCC value of 0.9270. Leveraging from DenseNet and Transformer architecture, PTFSpot (plant-based) showcased a narrow gap of only 1.74% between sensitivity and specificity. TSPTFBS 2.0 achieved the second-highest performance, with an average accuracy of 94.86% and an average MCC value of 0.9063, evaluated on a plant-specific dataset of 386 DAP-seq TFs. *k-*mer grammar secured the third position, with an average accuracy of 93.73% and an average MCC value of 0.8832, applied to core promoter regions, ChIP-seq peaks, and open chromatin regions. *k-*mer grammar classifies regulatory regions based on sequence features, such as *k-*mers by differential weighting key words. Additionally, the gap between sensitivity and specificity was 3.31%. For AgentBind, its two different architectures, DanQ (RNN/LSTM) and DeepSea (CNN), attained the accuracy values of 92.33 and 90.38%, respectively. However, the gap between sensitivity and specificity was 4.29 (DanQ) and 3.8% (DeepSea). DNABERT followed the same, achieving an average accuracy of 89.54% and an average MCC value of 0.7950, while exhibiting a notable difference of 3.1% between sensitivity and specificity. TSPTFBS, which was trained and evaluated for 265 TFs for *A. thaliana,* achieved an average accuracy of 86.32%. LS-GKM, evaluated for 322 for humans TFs, attained an average accuracy of 86.50%. However, TSPTFBS edged out LS-GKM with an average MCC value of 0.7722 compared to LS-GKM’s average MCC value of 0.7316. BPNet, attained 85.96% accuracy with a gap of 6.13% between sensitivity and specificity. BPNet scored 0.7581 MCC score. Two other tools, MAResNet and DESSO, were also evaluated for 690 human ENCODE ChIP-seq experimental datasets. They achieved very similar average accuracy of 85.61% and 85.11%, respectively. The difference could be observed for sensitivity, where DESSO had a higher success rate to identify TFBRs compared to MAResNet, which had a higher average sensitivity value of 88.24%. They exhibited similar ranges of average MCC values of 0.7533 and 0.7545 for MAResNet and DESSO, respectively. We further observed that two more different tools with distinct architectures, namely KEGRU evaluated for 125 human ENCODE ChIP-seq experimental datasets and gkmSVM evaluated for 467 human ENCODE ChIP-seq experimental datasets showed a very thin line of difference in their average accuracy(84.72% and 84.31%, respectively). While both the software tools had sensitivity and specificity difference of 1.83% and 2.48%, respectively.

On Dataset “A”, the performance exhibited by the 19 tools on their respective datasets aligns closely with the reported or calculated performance metrics. Notably, PTFSpot demonstrated exceptional performance, surpassing all other tools. Second to it was TSPTFBS 2.0, which also utilized the DenseNet in its architecture. DenseNet’s multi-layered structure enables efficient feature extraction and propagation, addressing challenges such as the vanishing gradient problem. The success of these architectures underscores their potential in TFBRs identification, highlighting their significance in regulatory genomics. Additionally, *k*-mer grammar, designed specifically for plant-based datasets, performed impressively as the third-best tool by leveraging *k*-mer frequencies for information capture. DNABERT and LS-GKM, designed specifically for identifying human TFBRs, also demonstrated robust performance. DNABERT, a proficient language model, effectively learns semantic and syntactic patterns inherent in DNA sequences. Overall, the performance results across all software tools exhibited notable similarity, validating the consistency of reported results with the findings in **Figure 8A** and **Supplementary Table S4 Sheet 1**.

### Animal specific tools remodeled for plants perform well

A thorough evaluation of all the 19 software tools was done on plant-specific Dataset “B”, comprising 265 DAP-seq experimental TFBRs data for *A. thaliana* (**Supplementary Table S5**). Comprehensive performance details are presented in **Figure 8B**, and **Supplementary Table S4 Sheet 2**. Once again, PTFSpot (plant-based) emerged as the top performer, achieving an impressive average accuracy of 96.14% and the high MCC value of 0.9190. Notably, PTFSpot demonstrated a remarkable balance between sensitivity and specificity, with a mere 0.48% gap. Again, TSPTFBS 2.0 secured the second position, with an average accuracy of 95.78% and MCC value of 0.9169. KEGRU obtained the third-best performance with an average accuracy of 94.79% and MCC value of 0.8967, showcasing a commendable balance between sensitivity and specificity (0.71% difference). LS-GKM achieved an average accuracy of 94.29% and MCC value of 0.8862. DESSO followed closely with an average accuracy of 94.24% and MCC value of 0.8935. DNABERT attained an average accuracy of 91.92% with MCC value of 0.8394, showcasing a 2.66% difference between sensitivity and specificity. Additionally, gkmSVM and TSPTFBS, for human and plant TFBRs identification respectively, performed equally well with average accuracy values of 91.55% and 91.51%, respectively. However, gkmSVM exhibited a slightly better balance between sensitivity and specificity, with a 0.3% gap compared to AgentBind (DanQ)’s 0.6% and TSPTFBS’s 0.83%. Nevertheless, AgentBind (DanQ) and TSPTFBS achieved higher average MCC values of 0.8773 and 0.8543 compared to gkmSVM’s MCC value of 0.8314. Among the plant-specific tools, SeqConv, *k*-mer grammar, and Wimtrap achieved average accuracy values of 87.7%, 87.67%, and 82.09% respectively. They obtained average MCC values of 0.7895, 0.7901, and 0.6428 respectively, with *k*-mer grammar displaying a notable balance between sensitivity and specificity (0.14% gap).

Among the top nine best-performing software tools, which achieved an average accuracy scores exceeding 90%, six were originally designed for human ChIP-seq data. While, TSPTFBS and its updated version, TSPTFBS 2.0, and PTFSpot were initially developed for plant TFBRs identification. Results from Dataset “B” demonstrate the efficacy of animal-based tools on plant-specific datasets, exhibiting higher average accuracy values and MCC scores with minimal gaps between sensitivity and specificity. It is worth noting that some of the top-scoring animal-based software tools may perform well for plants if retrained with plant-specific data. Among these top-performing software tools, KEGRU, and DESSO utilize DL algorithms, while LS-GKM and gkmSVM are based on SVM, and DNABERT employs the Transformer BERT algorithm.

On the sidelines, before closing this section, a small analysis was also performed to check the impact of using class imbalanced datasets. The reason is an argument that in actual condition imbalanced distribution of the positive and negative instances happens. Logically, a good learner will not get affected as well as very large datasets also mitigate such issues. Besides this all, a balanced dataset discourages any skew in learning due to imbalanced dataset. Therefore, in this part of study we used a skewed dataset where the negative instances were increased in the ratio of 1:4 of positive vs negative instances, and all these compared tools were retrained an evaluated. We found that training with such imbalanced datasets only negatively influenced all these tools and their models and their performance reduced. In this reduction, the sensitivity was more affected while specificity was slightly affected for the positive side. With the use of class-imbalance-aware loss function, their performance could be improved, but yet remained lesser than the performance observed with the training on the balanced datasets. Therefore, it is not recommended to use imbalanced datasets in designing such models as the model performance gets negatively affected. **Figure 8C** illustrates the findings for this part of analysis.

### Most of the tools have limited practical utility due to poor cross-species performance

Consequently, the search for reliable algorithms for TFBRs discovery in plants is of utmost importance, with an essential requirement for cross-species functionality. In the actual application of cross-species TFBRs identification, we found a huge performance gap as discussed below. Hardly a few tools have reported their practical applications to identify cross-species TFBRs, such as DNABERT (human-based), *k*-mer grammar, TSPTFBS, TSPTFBS 2.0, SeqConv, Wimtrap, PlantBind, and PTFSpot. However, the criteria they followed to identify common TFBRs between related species were quite different. DNABERT fine-tuned the human pre-trained model genome model on 78 mouse ENCODE ChIP-seq experimental datasets. They reported that these pre-trained models on human datasets significantly performed on the mouse datasets, demonstrating the robustness and application of DNABERT even across different genomes. When we analyzed its performance for cross-species TFBRs identification, it did not stand to its promise.

For the tool *k*-mer grammar, they developed models for maize and used them to study the regulatory environment of closely related species. They trained the models to preferentially capture conserved features (those shared throughout lines or related species). They examined models that were trained on TF binding loci (*Zea mays* KN1) and tested across the core promoters in two species (sorghum and rice). Using the concept of transfer learning, TSPTFBS demonstrated a practical application of the CNN models, trained for 265 TFs from *A. thaliana*. They tested it across other species. They gathered ChIP-seq datasets for 10 TFs from three different plant species, including four TFs from *Oryza sativa* (NAC6: GSM2748278, BZIP23: GSM2152800, ERF48: GSE93081, MADS29: GSE42201), four TFs from *Zea mays* (ARF5: GSM1925000, O2: GSE63991, P1: GSE38587, KN1: GSE39161), and two TFs from *Glycine max* (GLYMA.06G314400: GSE101672, GLYMA.13G317000: GSE101672). These TFs were screened on the basis of two requirements: (a) Its protein sequence must be very similar to at least any of the *Arabidopsis* TFs (as determined by BLASTP with an E-value of 1.0e^-5^), (b) or its TF motif must be highly similar to at least one *Arabidopsis* TF (as determined by TomTom with a p-value of 1.0e^-3^). Ironically, both these conditions laid down by the authors actually go against the concept of cross-species variability, and no way truly reflect the cross species application but evade answering it, as will transpire in the sections below, plants exhibit enormous genomic variability.

For TSPTFBS 2.0, which used ChIP-seq datasets for 15 TFs for six additional species were retrieved from ChIP-Hub, covering *Eucalyptus grandis*, *Hordeum vulgare*, *Physcomitrella patens*, *Arabis alpina*, *Chlamydomonas reinhardtii*, and *Solanum lycopersicum*. The fourth one, SeqConv, simply applied their applicability by transferring a *Z. mays* ChIP-seq pre-trained model to *A. thaliana* one DAP-seq TF dataset and reported an accuracy over 80%. For PlantBind, in order to test the application of the pre-trained model (*A. thaliana*) on datasets of *Zea mays*, they gathered ChIP-seq datasets (Zm00001d031717, Zm00001d042907, Zm00001d024644, and Zm00001d029875) for four maize TFs. They chose the comparable counterparts from maize based on their strong similarity to at least one *Arabidopsis* TF motif, determined by TomTom with an E-value cutoff of 1.0e^2^. As observed with TSPTFBS 2.0, this again ensured incorrect assessment of cross-species applicability while clearly ignoring the variability. On the contrary to these software tools, Wimtrap looked at how well pre-trained models could be applied to different conditions or species. Only the pre-trained models on flowers of *A. thaliana*, ripening fruits of *S. lycopersicum*, seedlings of *O. sativa*, and seedlings of *Z. mays* were used to identify the binding sites in the *A. thaliana* seedlings, with the highest AUC of 86% on ripening fruits of *S. lycopersicum*. Only, PTFSpot appeared truly designed on the concept of cross-species transfer learning on the basis of DNA sequence and TF protein three dimensional structure/sequence co-variability, while being completely free from such similarity and TF specific model based conditions. PTFSpot was tested across wide range of species and TFs, including those for which no model ever existed.

In our assessment for cross-species performance and applicability of these tools, we listed common TFs among different plant species where limited ChIP-seq and DAP-seq datasets are available. And this number becomes much lower when one looks for common TFs across the species. We succeeded in getting only five common TFs, namely LHY1, MYB62, MYB81, MYB88, and WRKY25 between *A. thaliana* and *Z. mays* **(Supplementary Table S1 Sheet 3)**. This dataset was considered as Dataset “C”, which was used to assess the cross-species performance of these tools. We used the pre-trained models for these common TFs, trained on *A. thaliana* DAP-seq dataset from Dataset “B”, and tested all 19 software tools on the five common TFs from *Z. mays*. However, here we wish to point out that gkmSVM, PlantBind and Wimtrap were found unfit for the cross-species TF model evaluation for these common TFs. The main reason is the fact that gkmSVM has an inbuilt function for their model evaluation, which does not provide the facility to load data for independent evaluation. For PlantBind, as we can also observe in **Figure 8A&B**, the performance on Dataset “B” was below an average accuracy of 75%. Therefore, we did not consider its pre-trained model on Dataset “B” for cross-species TFBRs identification. On the other hand, for Wimtrap, the PWM for the respective TFs were required to perform the task of sub-sequence scanning in the test file for data input along with the TF peak files. This requirement may not work for many species as species-specific motif information is mostly lacking in plants, and in fact, this is one of the major quests of TFBRs discovery in plants. A limited amount of information on *A. thaliana* TFs PWMs is available, but almost nothing is available for other plant species, including *Z. mays.* This is a big limitation, as actual TFBRs annotations are done on newly assembled genomes, where motif information is rarely present. Therefore, we had to discard these tools for Dataset “C” based cross-species evaluation.

As we further tested the rest of the 16 tools’ applicability in cross-species identification on an independent test dataset, we observed that common TFs trained models showed a very poor performance on Dataset “C” except PTFSpot as illustrated in **Figure 8D** and **Supplementary Table S4 Sheet 3**. Among all the software PTFSpot attained an average accuracy of 92.50% with MCC value of 0.8594. For the other software tools, named BPNet and KEGRU, the average accuracy values of 69.90% and 68.17% were noted. This may be noted that both, BPNet and KEGRU were primarily developed for animal system. However, as listed in **Supplementary Table S4 Sheet 3**, KEGRU worked well for the two TFs, namely MYB62 and MYB81, with accuracy values of 97.04% and 90.28%, respectively. A gap of 2.35% and 2.91% between sensitivity and specificity was observed for MYB62 and MYB81, respectively, with MCC scores of 0.9410 and 0.8058, respectively. It can be seen from **Figure 8D** that most of the tools displayed strong over-fitting. 10 out of the total of 14 software tools merely attained an average accuracy value between 50% and 60%. MAResNet, had scored for MYB88 with 67.78% accuracy, scoring sensitivity and specificity values of 75.90% and 59.66%, respectively. In addition to it, LS-GKM for MYB88, deepRAM for MYB62, TSPTFBS for MYB88, DESSO for MYB62, DNABERT for MYB88, and TSPTFBS 2.0 for MYB88 scored accuracy values of 66.70%, 64.72%, 64.17%, 60.85%, 62.49%, and 61.83%, respectively. In overall, the cross-species benchmarking results indicated that the effectiveness and applicability of all considered tools in identifying TFBRs across plant species were surprisingly low except for PTFSpot. Thus, PTFSpot appears to be the most reliable tool for cross-specific and practical application to identify TFBRs in plants. PTFSpot successfully identified TFBRs across other plant species and even with unseen TFs, while doing parallel learning upon the structural variations and corresponding alterations in binding region partners.

TFs and their binding sites vary across species, with changes in TF structure leading to differences in binding regions **[11,26–31]**. This variability causes most existing tools, except PTFSpot, to fail in cross-species scenarios. Our study analyzed peak data from 10 common TF families across five plant species (*A. thaliana, Brassica napus, Glycine max, Hordeum vulgare*, and *Triticum aestivum*). Using Alphafold2 **[132]** to model 3D structures due to the lack of experimentally obtained structures for most plant TFs, we found significant structural differences, with RMSD values above 0.401Å (**Supplementary Table S6 Sheet1** and **Supplementary Material S2 Figure 2**). For example, the WRKY TF family showed an RMSD of 1.285Å between *A. thaliana* and *Brassica napus*, while the homeobox TF family had an RMSD of 1.472Å between *A. thaliana* and *Glycine max* (**Figure 9** and **Supplementary Table S6 Sheet1**). This highlights the substantial variability in TF structures across species, a factor that pre-trained models often overlook, affecting their cross-species applicability. PTFSpot successfully addresses this variability, creating a universal model **[66]**.

**Figure 9:**
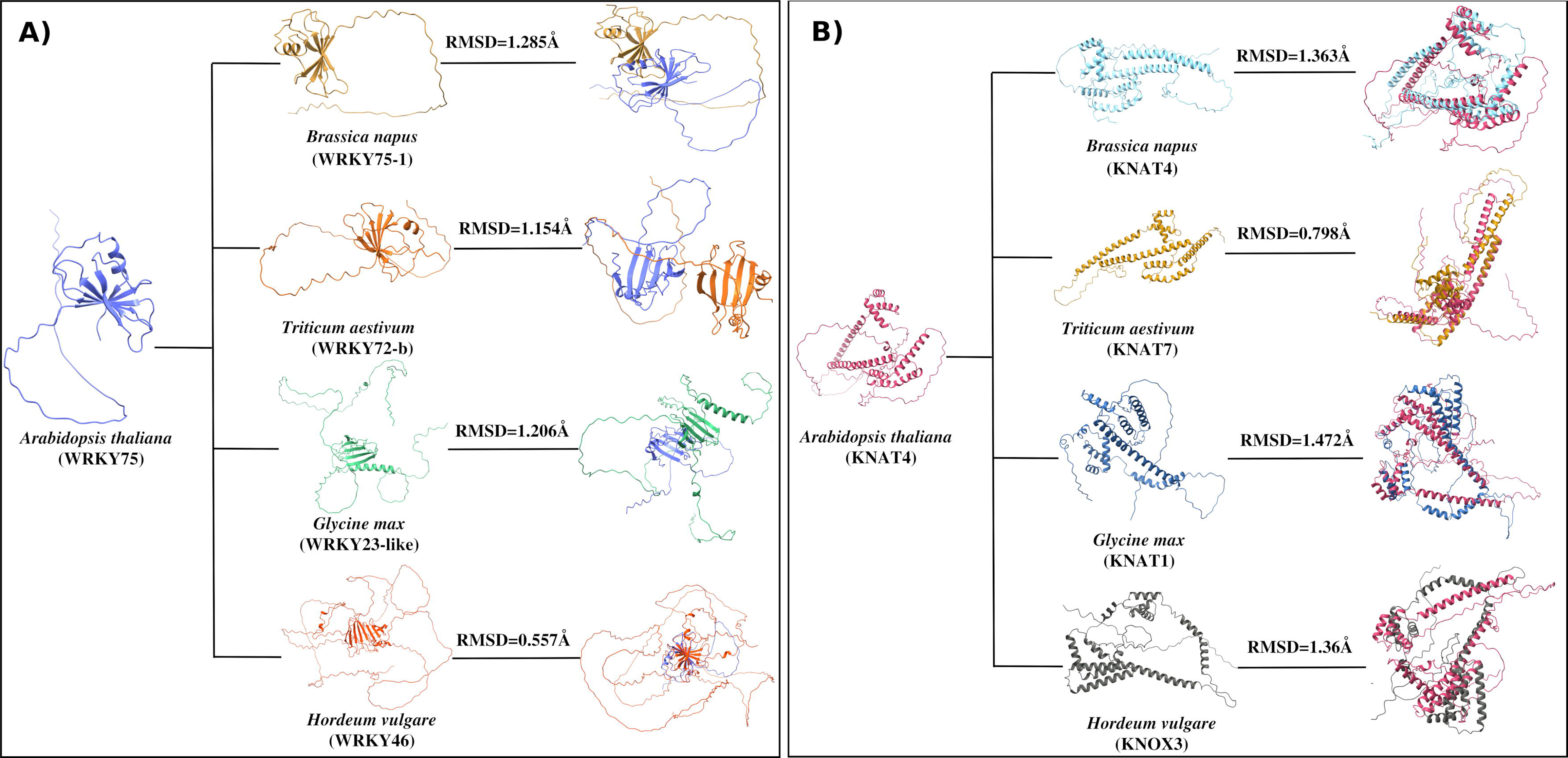
3D structure based co-variation in transcription factors families’ structure across five plant species (*A. thaliana* vs *Brassica napus*, *A. thaliana* vs *Triticum aestivum*, *A. thaliana* vs *Glycine max*, *A. thaliana* vs *Hordeum vulgare*). (A), (B) The corresponding superimposed TF structures with the RMSD value indicating the structural variations across TF families, named WRKY and ethylene, respectively.

### Need to redefine the transcription factors binding sites (TFBS) as binding region marker motifs

As described above, the problem of transcription factor binding region discovery is closely knitted with motif discovery, where the enriched motifs are assumed to be the binding site. While the fact is that both the problem are separate. Earlier methods would use such enriched motifs identified from the binding experiment data to scan them across the genomes. Later on, with advent of novel machine learning based approaches, it has reversed to finding TFBRs and proposing such motifs using enrichment tools or feature weights analysis. This part defines the final leg of computational challenges where the algorithms can be clearly viewed into two different groups: A) Traditional clustering based unsupervised approaches which apply statistical enrichment and probabilistic models, and B) Deep-Learning guided motif discovery.

In this direction study of total nine most popular and recent tools was conducted, which included five traditional motif identifiers, namely HOMER, CMF, MEME, MEME-ChIP, and iFROM. Besides this, four recent and DL based tools were also involved (deepRAM, DESSO, TF-MoDISco, and PTF-Vāc). These tools were studied for their performance based on the following three major criteria: 1) the influence of input sequence data on binding regions discovery, 2) overlap and similarity with experimentally reported motifs, and 3) coverage of overall experimental binding data provided for any TF (**Figure 10**). The proposed motifs by each such tool were compared with the 10 TFs experimentally reported motifs from the JASPAR database **[133]** and Plant Cistrome Database. We used the available DAP-seq dataset for ten different TFs to scan and report their binding sites across these sequences.

**Figure 10:**
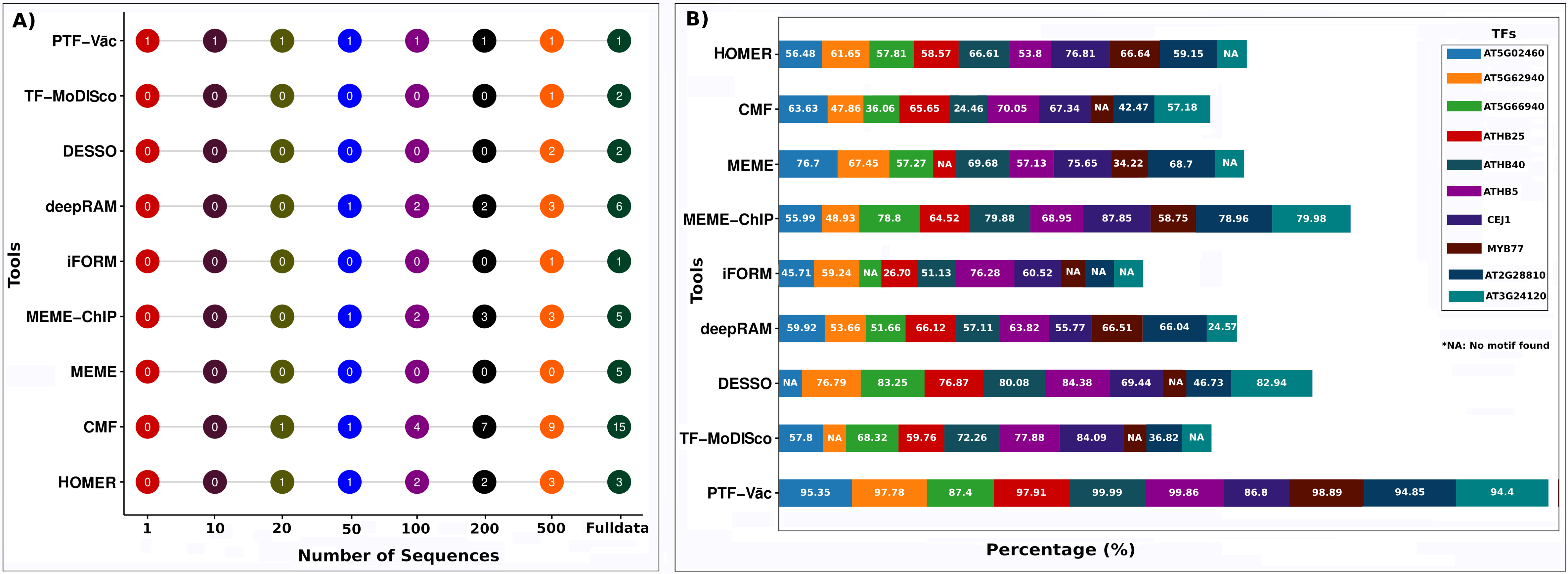
Comparative performance of nine different tools for TFBS motif discovery. This involved five traditional motif identifiers (HOMER, CMF, MEME, MEME-ChIP, and iFROM) and four deep learning-based based tools (deepRAM, DESSO, TF-MoDISco, and PTF-Vāc). It evaluates their performance based on two major criteria: **(A)** the impact of input sequence data on binding motif discovery, and **(B)** overall binding data coverage for 10 different TFs. PTF-Vāc stands out for its ability to discover motifs irrespective of input data size and achieve high data coverage of identified motifs. It displayed no sensitivity towards input data volume also.

The effect of input sequence data size on the discovery of the TFBR motifs were evaluated for the performance and also measured the considered software tools for every considered input size (binding region data). It was found that, as claimed by PTF-Vāc also, every tool was sensitive towards the volume of input data taken and their results varied with the input data sizes ( **Figure 10A**). Some of the tool, for instance, HOMER and CMF required minimum 20 sequences, while deepRAM and MEME-ChIP required at least 50 input sequences to identify motifs. Multiple motifs were identified by tools like CMF, MEME-ChIP and deepRAM when 100 sequences were provided. TF-MoDISco, DESSO, iFORM, and MEME only discovered motifs with more than 200 sequences. In contrast, PTF-Vāc was able to discover the motifs regardless of the number of input sequences. This pattern was observed for all the TFs studied here. Additionally, as observed earlier (PTF-Vāc), MEME, iFORM, DESSO, and TF-MoDISco often produced motifs for a given TF when provided with a sufficient number of input sequences. For instance, as shown in **Figure 10A**, MEME produced motifs when number of input sequences raised to full dataset for ATHB5. On the other hand, iFORM, DESSO, and TF-MoDISco, both started to produce motifs when number of input sequences were above 200. While taking the case of TF ATHB5 as an illustrative example in **Figure 10A**, it is clear that for the varying input data size, only PTF-Vāc was not sensitive towards the volume of input data while others displayed a variable range of sensitivity towards the volume of input data and identified motifs and their numbers.

In the second phase of the present study, as showcased by PTF-Vāc work, here too assessment was made for the overlap and similarity between the motifs discovered by various software tools and experimentally reported motifs for 10 TFs (**Supplementary Table S7 Sheet 1)**. For these TFs the length of their reported motifs ranged from 11 to 21 base pairs. The best performer among all was PTF-Vāc which gave an average similarity of 96.61% for the compared motifs for all the 10 TFs. Next it was MEME-ChIP which had 73.78% average similarity, and deepRAM had 68.75% average similarity for all the compered 10 TFs (**Supplementary Table S7 Sheet 1**). **Supplementary Table S7 Sheet 1** provides a list of motif similarity between discovered motifs and experimentally reported motifs for different software tools on all these 10 TFs. Among all the tools, HOMER and deepRAM exhibited minimal overlap with the experimentally reported motif for MYB77 motifs showing 40% and 45% overlap, respectively. Motifs discovered for AT5G62940 and AT5G02460 by iFORM matched with reported with 25.5 and 30 similarity. For DESSO two motifs for AT5G62940 and AT5G66940 showed 80% similarity with their reported motifs. However for deepRAM, the proportion of similarity for all motifs ranged from 66.7 to 74.55%. For MEME, AT5G02460 and AT5G66940 the similarity of 62.5% was reported. While, CMF, iFORM, DESSO, and TF-MoDISco, did not discover any motif similar to reported one for MYB77. The same was noticed for AT3G24120 for which TF-MoDISco, iFORM, MEME, and HOMER were unable to identify any motif. Moreover, 80% similarity was reported for ATHB40 (MEME and TF-MoDISco), and ATHB5 (MEME-ChIP). In general it was observed that most the tools were reporting different motifs for the considered TFs which were not significantly closer to the reported motifs, except PTF-Vāc and MEME-ChIP. Most of the tools, as also mentioned above, gave multiple motifs due to large input size. Due to this some of their reported motifs were found common among themselves, while they disagreed with each other for a good number of reported motifs (**Supplementary Table S8 Sheet1-3**).

While assessing for the 3^rd^ and final criterion, this was observed that motifs reported by the most of the existing tools did not cover the binding data sufficiently and were representative of just a part of the complete binding data, clearly suggesting their failure to propose any reliable universal representative/marker for the TF-DNA interactions. An assessment of the overall binding data coverage for all 10 TFs by all the compared software tools was performed (**Figure 10B**). From the analysis, it was found that as reported already by PTF-Vāc, it exhibited the highest data coverage for the majority of TFs, with coverage ranging from at least 94.4% to 99.99% for eight TFs (AT3G24120, AT5G02460, AT5G62940, ATHB25, ATHB40, ATHB5, AT2G28810, and MYB77). The lowest data coverage by PTF-Vāc identified motifs were 86.8% and 87.4% for CEJ1 and AT5G66940 TFs, respectively (**Figure 10B**). From this figure, it can be clearly seen that HOMER, MEME, iFORM, DESSO, and TF-MoDISco were not capable to discover motifs for many of the TFs. Even for the reported motifs by these tools, the experimental binding data was poorly covered by many of them. For example, HOMER’s eight TFs, CMF’s all nine TFs, MEME’s six TFs, MEME-ChIP’s five TFs, iFORM’s five TFs, deepRAM for all the 10 TFs, DESSO’s two TFs, TF-MoDISco’s four TFs, showed the data covered by their discovered motifs less then 70% of the complete experimental binding data for each TF.

Whether the probabilistic traditional approaches or most of the recent deep-learning based approaches, there appears to be huge scope of improvement. Motif finding by them would gather much sense if biologically relevant information are also infused, instead of mainly performing the statistical and enrichment exercises. Sparsity in such motifs is another important factor which they need to deal. Yet, as transpires from from the **Figure 10** and above mentioned results for three parameters of benchmarking, PTF-Vāc, emerges as the best solution for relevant motif discovery in plants, maintaining huge lead over the second best, MEME-ChIP, and the third one, DESSO. MEME-ChIP is a meta-program which involves three different algorithms (MEME, DREME, and CentriMo) to finally propose a motif which make it take note of positional conservation even for a single nucleotide and shorter stretches, making it able to deal with sparsity for some extent. Good performance of DESSO can be attributed to the infusion of biologically important feature of DNA Shape through deep-learning, which the probabilistic models other studied tools largely miss. A first of its kind a*b-inito* capability for motif discovery using generative and explainable AI while learning from co-variability of TF and its binding preferences forwarded by PTFSpot **[66]** appears to be the major reason for the enormous lead observed for PTF-Vāc. It is also the only tool which can identify binding sites and motifs in complete independence from TF and species models, and even for single sequence, which none of the above mentioned tools are capable of. It also makes it suitable choice for scenarios where novel genome has to be annotated or a user has to scan the promoters of only few selected genes as PTF-Vāc is immune to data-volume sensitivity to propose motifs/TFBS.

## Discussion

Transcription is a fundamental biological process that defines the identity and function of a cell, guiding its response to environmental signals and determining its future behavior. This process is the core to life that it not only establishes the current state of a cell but also influences its trajectory, shaping the cell’s fate through dynamic changes in gene expression. Central to this all are transcription factors and their binding spots across the DNA which regulate downstream gene’s expression. Thus, finding these binding regions of various TFs is one of very important question. Using various binding experiments specific to TFs these interactions are identified. However it is a very costly affair in terms of time, efforts and money, and quite an impractical approach to evolve general models and theories. Therefore, one essentially requires software tools in this area of research.

Computationally, the major challenge is to evolve a universal model of TF-DNA interaction, in center of which stands the finding of TFBRs. Developing TFBR models require dealing experimentally reported binding data from ChIP and PBM based approaches. While after developing TFBRs, many have interest in determining the TFBS motifs. As already discussed in the introduction and results parts that initially such TFBS motifs alone were attributed as binding sites of TF during the early and traditional approach, which is not rightly placed approach as binding of TF depends upon a long range of sequence scanning by TFs and flanking region’s properties **[66]**. Therefore, finding TF-DNA interaction has three primary computational stages to pass through, and all three are intricately related with each other where finding TFBR is the main challenge, preceded by computational approaches for rightful peak-calling from the experimental binding data, followed by computational approaches to identify TFBS in the identified TFBRs. Thus, the present study has systematically carried analysis of all these three stages of software tools involved, their working approaches, and how credible each have been, presenting a clear guideline for all working in the field to systematically plan their studies based on the findings and recommendations made here. Many of these tools were found not well maintained and nor runnable by the scientific community and biologists. This study has also tackled such issues, raised the flags accordingly, fixed them and has released the corrected codes of these tools in public domain which the community now can use.

The first stage of computational challenge belonged to generating a reliable experimental binding dataset from the ChIP based experiments for which various peak-calling tools are used. We studied several peak-calling tools, which read the signal intensity arising from the different base labels for corresponding genomic bins, followed by statistical interventions to report the credible peak regions, the binding regions. It was found that the tools which apply background signal normalization and local signal statistics tend to perform more reliably than those who don’t apply that. Also, consideration of Irreproducible Discovery Rate (IDR) improves the reliability of the identified peaks. In overall performance, it was found that MACS, LanceOtron, and PeakRanger emerged as the best ones to work with. Their common agreements among themselves was high as well as they had least false peak identifications rates. Also, in terms of response to IDR consideration also, they maintained the same ranks. Also, the tools at this stage don’t need to be specific for plants and are able to work for all species.

The output of the previous stage of software tools, the peak regions, reflecting the binding regions in the range of 70-200 bases long sequences, become the input to the second and most important stage of the challenge, raising the computational models of TFBRs. These models are backbone to computationally identify TF-DNA interactions, but same time also attract the attention over a large number of factors affecting the performance, ranging from the approach (ML or DL), kind of data used to train the system, importance given to properties beyond the motif regions, and above all, consideration of variability across the plant genomes.

Most of the existing tools were found having of bugs creating difficulties in being run by the users. The study while analyzing these tools, faced these issues and fixed them, rendering them usable by the community. Barring a few, most of these tools have applied a similar approach towards building the dataset. Positive datasets don’t create issues, but inappropriate negative datasets mislead a lot. We observed that majority of these tools have centered around the motifs and used them as the marker of positive datasets and involved negative datasets devoid of them completely. Also, sequences were picked random in many cases, many instances of which were found overlapping with positive instances. Both these factors have significantly compromised the quality of the models raised by these tools. A motif alone can’t be claimed as TFBS as flanking regions conditions and context mainly determine the binding of the TF. A motif may be significantly enriched in the binding data, but it may also be present in non-binding regions. Thus, a negative data must have instances with the motifs found also in the positive instances, ensuring a strong, near to the real dataset to learn from. Care about these factors have been taken by tools like PTFSpot which has resulted into its superlative and consistent performance across all types of datasets taken in this study. At the stage of TFBR discovery it is highly beneficial to use such tools which don’t base them on motifs as priority.

The next notable factor was the impact of the learning algorithms used. There was a clear performance difference due to this factor, where the advanced deep-learning based tools were consistently found better performing. The best performers were strong deep-learning tools like PTFSpot and TSPTFBS 2.0 which employ Transformers and enormous variability across the species in their promoters DenseNets which have capabilities to learn the context and counter the vanishing gradients while going very deep into learning (**Figure 8B**). The RNN based tools scored thereafter, which are better capable than CNNs in learning the local sequential patterns, but can’t learn over long ranged relationships like transformers. The lowest performers were CNN and ML approach. We found that most of the CNN based tools hardly try to present more evolved data encoding and mainly focused upon directly learning from plain reading of the four bases. CNNs are good in spatial pattern recognition, but lack in deciphering context, and become more weak if encoding is not done with attempt to present more informative input. Best example of this observation is the performance difference between the two versions of AgentBind. Its RNN version (DanQ) performed better than its CNN version (DeepSea).

The least performing tools were shallow learners, the ML tools, which unlike the deep-learning algorithms, can’t decode the hidden features and have limited learning abilities. They just learn directly on the input data features provided by the person developing it, and don’t make any attempt to go further and try extracting more information and hidden relationships. Thus, designing an efficient ML model is subject to the expert knowledge developing it. However, there was a strong exception to this observation in the form of LS-GKM, a SVM based method which outperformed many Deep-Learning tools for TFBR detection. But it appears to be more attributable to the kind of features used than the machine learning approach. Unlike most of the Deep-learning tools which this simple SVM based tool outperformed, it did not orient itself to any motif, nor did limit itself to single base one-hot encoding to learn from, but relied upon decoding the various *k*-mers based representations, which could capture multiple motifs syntax. For all these *k*-mers, the authors have developed dedicated kernels for learning. LS-GKM thus clearly becomes an example to learn that how incorporation of biologically important features can boost the performance of even a simple machine learner and outperform much advanced tools.

Though LS-GKM used *k*-mers to represent a regulatory region, presenting the fact that consideration of features from long regions make more biological sense and thus enhance performance of the software tool drastically, it could not capture the inter-relationships between these *k*-mers, which is equally insightful biological information to detect regulatory regions. Learning on DNA Shape data near to motif site alone gave a good advantage to tools like DESSO, which we found contributing to it performance by 4%. We observed that all such tools benefited a lot when binding region characterization moved beyond motif discovery, and looked into context, surroundings beyond the motifs. All of the top performing tools from PTFSpot to AgentBind displayed that commonality which returned drastically improved performance than the previous generation of tools. The same we witnessed with simple machine learning approach of LS-GKM which outperformed more advanced deep-learning models due to such approach as already discussed above. This has a strong biological reasoning. Contrary to long held belief that a short stretch of motif is enough for TF-DNA interactions, in actual binding of a TF involves scanning long DNA sequence having characteristic signals to invoke halting of the scanning TF, not the binding motif along **[134,135]**.

The most important aspect emerging for this study is the matter of universality and practical use of these tools, where barring PTFSpot, almost all of these studied tools appeared failing to deliver. Value of any such tool is only when it could be used to annotate newly reported plant genomes for TFBRs. Which means that the developed models of the software tools for TF-DNA interaction should be universal and hold true across all the species. As transpires from the **Figure 8D**, barring PTFSpot, all the studied tools performed very poorly when applied in cross species manner. Even if the one attaining highest accuracy after PTFSpot, BPNET, could achieve just ∼70% accuracy. In fact, on this cross species analysis, all the studied software tools barring PTFSpot, can’t be even ranked for accuracy performance as on closely looking one find immense imbalance and gap between sensitivity and specificity values for all these tools, very well reflected by extremely poor and almost same range of MCC values for them. This clearly underlines that most of these developed tools are of no use for real applications like annotating new genomes. PTFSpot stands out as the only solution.

But why only PTFSpot works and rest all failed in true terms of practical applicability of cross-species use? The answer is “considerations for genomic variability” which is much profound in plants. Barring PTFSpot, none of the existing software tool tried to address and capture this most important factor for universal reach. All of them were found working with species specific models developed separately for each TF. Even within a species, if the TF model is not available, these tools can’t help in finding the TFBRs for such TFs. Ironically, most of the tools which claimed to be plant specific, have implemented hardly anything specific for plants except considering a plant specific datasets. This is why, as transpires from the study above, when the same plant specific datasets were used to train the software tools developed on animals, those tools not just came at par of the plant specific tools, but sometimes even outperformed the plant specific tools due to their better learning algorithms. Apart from it, between Datasets “A” and “B”, notable differences in tool performance were observed. This reason could be attributed to variations in how tools constructed their negative datasets and hyper-parameter optimization strategy. It was evident that existing tools generally overlooked the significance of the negative dataset, often using dinucleotide shuffle and random genomic sequences where similarities to positive instances existed. Few tools attempted to devise more robust methods for constructing higher-quality negative sets **[66]**, which would better reflect real-world scenarios. Consequently, a considerable difference in performance was noted, with most tools performing notably better on Dataset “B”.

Unlike animals system, where majority of focus has been around human as the prime model organism, in plants the prime model organism has been *Arabidopsis thaliana*, a plant with very simple genome and which does not capture the levels of complexity and diversity plant genomes witness **[11,26–31]**. While the fact is that, compared to animals, plants display enormous variability across its genome affecting almost every genomic element, be it protein coding genes or promoters or transcription factors and their binding sites **[20,23,136,137]**. **Figure 11** very lucidly illustrates the variability degree differences between animal genomes and plant genomes.

**Figure 11:**
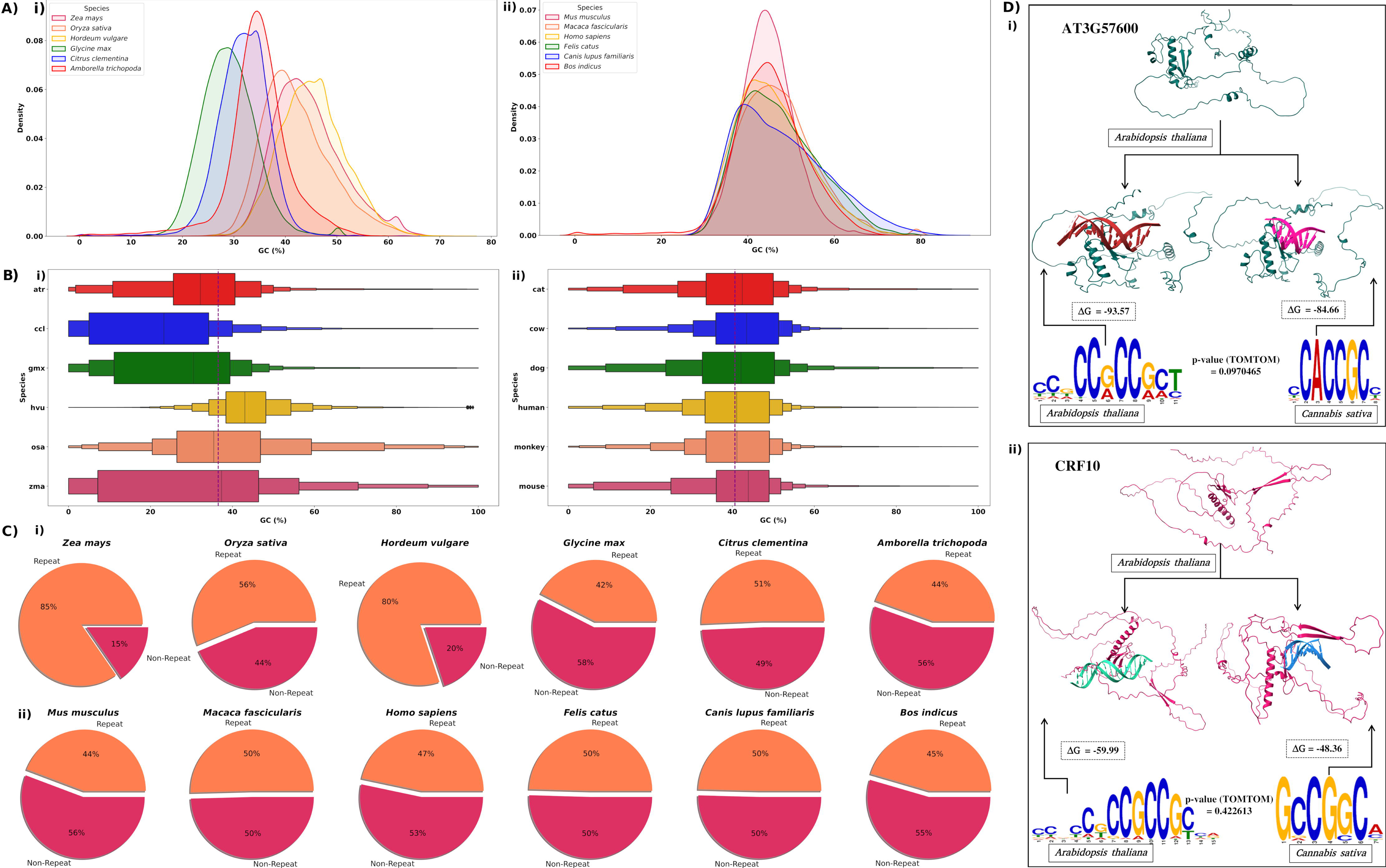
Variability across plants and vertebrates genomes. **(A)** Total GC content distribution of promoter sequences in **(i)** plants, and **(ii)** vertebrates genomes. **(B)** Total GC content distribution of repeat regions in **(i)** plants, and **(ii)** vertebrates genomes. **(C)** Total repeat content variations across **(i)** plants, and **(ii)** vertebrates genomes. **(D)** The TF structure and its binding site preference varies across the species, which is much profound across plants than animals. For the similarly distanced species in plants and animals, the TFBS motif also exhibited significant variations in plants than animals. This variation affects the stability of TF-DNA interactions in plants significantly.

Animals display almost same range of GC content in their 2kb upstream promoter regions, while plants display enormous variability across the species in their promoters **(Figure 11A i-ii)**. Plants are enormously enriched for complex repeats while displaying lots of variability across the types of repeats and even within the types, when compared to animals **(Figure 11B i-ii)**. Animal genomes appear to be almost static with their repeat contents, their composition, and types, plant are not. Therefore, it is very likely that the TFs across the plant species would display complex repeats while displaying y such variability and in turn would display variability in their binding preferences, which had become the background of PTFSpot. An analysis for five TFs and their reported binding motifs between phylogenetically similarly distanced species in animals (Human vs Mouse) and plants (*Arabidopsis thaliana* vs *Cannabis sativa*) **[138,139]**, further supported this view. Between human and mouse, the TFBS did not vary a lot, nor their binding preferences were much lowered. But in the plants, variation in TFBS motifs were so much substantial that their TOMTOM scores became insignificant as well as their binding preferences lowered drastically. **Table 3 and 4** and **Supplementary Material S2 Figure 3&4** highlights this finding while discussing TFBS variance and structural and docking analysis for these species and their TFs.

On the similar lines, in **Figure 9** and **Figure 11D** above, while considering the two plant TFs for representation purpose, it was clearly illustrated that how much the structure of a TF varies as one walk across the various plant species, displaying substantial RMSD values when structurally compared. Therefore, it becomes mandatory to consider genomic variability to raise any relevant TF-DNA interaction detection software tool. And unfortunately, barring PTFSpot, almost all of the existing tools have missed this highly important factor and rendered themselves of almost no use in plant TF-DNA interaction research. PTFSpot captured this variability by learning upon the changes in binding preferences with TF sequence and structure. Unlike rest of the tools, it essentially involves TF sequence and structure also for which the binding regions have to be detected. It considers AlphaFold2 generated structure to measure the relative structural standings and derive its binding preference maps accordingly. Thus, for now, for cross-species universal and realist application, one can rely only of PTFSpot to detect TF-DNA binding regions in plants.

The final stage of these challenges of detecting TF-DNA interactions is reporting of the TF binding motifs, which has been regularly called as transcription factor binding sites (TFBS). This study, first of all, objected calling such motifs as TFBS and finds roots of such practice in the fact that the initial and traditional practice of computational detection TF-DNA interactions worked with smaller binding data where defining the binding preferences were restricted to short motifs. These motifs were reported using unsupervised statistical learning and clustering approaches, most popularly Gibb’s sampling and Expectation Maximization (EM), generating position weight matrices as TFBS model. Now, as transpires from above results and discussion sections, it is very clear that such motifs also occur a lot in the non-binding regions also, and TF-DNA interactions are dependent on many contextual factors spread across a good length of sequence. In such scenario, these TFBS should be called as preferred or marker motifs only which needs to be dealt separately as purely motif finding problem instead of calling it TFBS discovery. The second and recent phase of development in this area has addressed the same issue where most of the machine and deep-learning algorithms now focus mainly on TFBR discovery. Once they are found, then they define such motifs across them. However, they also apply similar approach in defining the motifs through enrichment or apply a newer approach of convolution layer based feature extraction to define such motif. Details of their different approaches are already given in the methods section of the article. The study found that despite being fresh in their approach of motif discovery, barring PTF-Vāc, most of the deep-learning algorithms were performing a bit lower than the traditional approach of MEME-ChIP. MEME-ChIP is a meta-program which has incorporated three different algorithms to tackle shorter and sparser motifs, which makes it capable to deliver better than the rest of the deep-learning tools studied here for such motif discovery. Also, the deep-learning tool’s motif finding stage is closely attached to their TFBR discovery stage, their motif discovery is largely influenced by the level of noise and false identifications in the their previous stages, which is not the matter with MEME-ChIP like tool which works standalone. However, MEME-ChIP, completely lacks any infusion of biological information except assumption of some relevant motif across the binding data. Also, lots of user defined parameters are required there and it becomes more like a statistical analysis package. Barring PTF-Vāc, all these tools were found immensely influenced by the kind and volume of binding/TFBR data being provided. None of them was able to work below 50 instances of binding regions and their number of different reported motifs linearly grew with increment in the volume of the input data. Interestingly, their reported motifs differed also among themselves, as well as did not cover the full experimental binding data significantly while sometimes even missed to identify any motif for some TFs (**Figure 10**). They also grossly disagreed with experimentally reported motifs for many TFs. Therefore, inconsistency with motif discovery tools was their one of the prime issues which also raises question of their reliability. PTF-Vāc stands out because it is completely unique in its treatment of identifying motifs by applying generative AI while using the wisdom of PTFSpot in its background to ensure that it doesn’t get any noise input data for any given TF while retaining biologically relevant information also. It also learning from the flanking regions and contexts to determine the most suitable motif candidates. Unlike most of the tools, it is also not sensitive towards input data volume and can work with same reliability level for single sequence or thousands of sequences. Its identified motifs went closest to the experimentally reported motifs and also covered the experimental binding data the highest. Since it is based on PTFSpot’s universal modeling approach, it can work for any TF and species, seen or unseen before. It applies encoder-decoder generative system on PTFSpot’s learning to speak the most important potentially relevant motif, where explainable AI is also applied to provide additional confidence score through Grad-CAM approach. Explainability has been the issue with deep-learning approaches, and this the hottest area of AI research at present also. The Deep-learners are usually called black boxes because of lack in transparency about their functioning which makes them hard to explain. This happens due to existence of several hidden features which are derived from the input layers and have mathematical explanations for derivation than their exact biological sense. Explainable AI is the latest movement to counter this issue with deep-learning algorithms.

This study has dealt with all these three major stages of computational challenges in identifying TF-DNA interactions in a highly extensive and comprehensive manner, giving a clear guideline about how to approach the problem of TF-DNA interactions in the present scenario. Also, the findings made in this study will prove itself very important for the development of future generations of TF-DNA interaction studies where factors like epigenomic regulation and long distanced interactions stand almost unattended and may be incorporated further. These tools can tell about the binding regions across a genome. But a more interesting question now arises, specially when we see tools like PTFSpot capable of bypassing DAP-seq like experiments, that is it possible now to tell when these TFs would interact at any given binding region/site? It is a worth millions of dollars question indeed!

## Conclusion

The present work has comparatively analyzed the three important parts involved in computational detection of transcription factors binding regions (TFBRs) in plants: peak calling, TFBR detection using peak calling data, and TFBS motifs detection using TFBR information. In doing so, wide range of algorithms and software tools have been evaluated for each phase. It was found that most of the existing tools developed for plants, baring a few, are not sufficiently reliable to identify TFBRs. In fact, it was found that some of the algorithms developed primarily using animal binding data when retrained upon the plants binding data performed far superiors. Yet, none of these approaches, except PTFSpot, was found capable to work in cross-species manner. PTFSpot was found capable to efficiently work in cross-species manner because it has learned covariability between TF structure and its preferred binding regions. Further to this, it was found that the existing approaches of reporting TFBS across the TFBRs has some serious limitations, including sensitivity towards volume of input data to report motifs, not sufficient coverage of experimental binding data by the reported motifs, and difference from experimentally reported motifs. *Ab-initio* TF specific motif discovery appears a solution to such situation. At present, there is only one such tool capable to perform this task, recently released PTF-Vāc, which has been built upon PTFSpot, learning from covariability between the TFBR and TF structure, capturing the most informative and biologically relevant signatures. It employs Transformers based encoder-decoders system. The tools which were having technical glitches have been categorized, corrected, and the corrected version have been made available to the community. The present study is expected to work as a guide towards understanding the software tools and steps involved towards successful discovery of plant TFBRs, which will help the plant biologists to carefully plant their study.

## Authors’ contributions

Jyoti carried out the most of the study. Jyoti, and Ritu carried out the benchmarking study. SG assisted in the study with analysis and evaluation. RS conceptualized, designed, analyzed, and supervised the entire study. Jyoti, Ritu, SG, and RS wrote the MS.

## Acknowledgments

The study was carried out under the aegis of The Himalayan Centre for High-throughput Computational Biology (HiCHiCoB), a BIC supported by the Dept. of Biotechnology, Govt. Of India. Jyoti is thankful to UGC for her fellowship. Ritu is thankful to CSIR for her fellowship. SG is thankful to DBT, India for financial support as project associateship. Jyoti, SG, and Ritu are thankful to Academy of Scientific and Innovative Research (AcSIR) for their Ph.D. enrollment. The authors are thankful to Umesh Bhati for his assistance in MS writing. All authors are thankful to the Director, CSIR-IHBT, for his kind support for this study.

## Declaration of competing interest

The authors declare that they have no competing interests.

## Ethical approval

Not required.

## Ethics statement

Not applicable.

## Data and code availability

All the secondary data used in this study were publicly available and their due references and sources have been provided in **Supplementary information**. All data and information generated/used, methodology related details etc. have also been made available in the Supplementary data files provided. All the corrected running scripts for each tool have also been made available at Github at https://github.com/SCBB-LAB/Comparative-analysis-of-plant-TFBS-software.

## Supplementary information

**Supplementary Table S1:** Detailed description of Dataset “A”, Dataset “B”, and Dataset “C”.

**Supplementary Table S2:** Dataset description of motif finding software tools for 10 DAP-seq transcription factors.

**Supplementary Table S3:** Performance metrics of all the the 19 software tools on Dataset “A”, which comprises of the datasets used by 19 software tools.

**Supplementary Table S4:** Comparative evaluation details of all the tools on Dataset “A”, Dataset “B”, and cross-species Dataset “C”.

**Supplementary Table S5:** Performance metrics of all the the 19 software tools on Dataset “B”, a DAP-seq dataset for 265 plant transcription factors.

**Supplementary Table S6:** 3D structure based co-variation in transcription factors families’ structure across five plant species (*A. thaliana* vs *Brassica napus*, *A. thaliana* vs *Triticum aestivum*, *A. thaliana* vs *Glycine max*, *A. thaliana* vs *Hordeum vulgare*).

**Supplementary Table S7:** Comparative evaluation details of all the motif identification tools on *A. thaliana* DAP-seq dataset.

**Supplementary Table S8:** Comparison of the motifs identified by software tools with each others’ identified motifs and with reported motifs of *Arabidopsis thalaina* and *Zea mays*.

**Supplementary Material S1:** List of the existing experimental techniques, list of peak calling software tools, some published tools for transcription factor binding regions (TFBRs) identification, as well as databases available for both human and plant research.

**Supplementary Material S2: Figure 1** represents number of peaks overlap in four TFs binding regions between six software tools and **Figure 2** illustrates 3D structure based co-variation in transcription factors families’ structure across five plant species (*A. thaliana* vs *Brassica napus*, *A. thaliana* vs *Triticum aestivum*, *A. thaliana* vs *Glycine max*, *A. thaliana* vs *Hordeum vulgare*). **Figure 3** and **4** show the the co-variation in the structure of the transcription factors and their corresponding binding sites across plants (*A. thaliana* and *Cannabis sativa*) and (*Homo sapiens* and *Mus musculus*) species. TF structure and its binding motif comparison with experimental validated binding motif of TFs and its 3D structure and the docking analysis within and across other TFs.

**Supplementary Material S3:** Analysis, reporting, and solutions for bugs/errors encountered in all benchmarking software tools.

**Supplementary Material S4:** A complete list of dependencies for successful running of BPNet environment.

**Supplementary Material S5:** Hyper-parameters optimization to optimize the performance of TFBRs identification software tools.

## Notes

https://github.com/SCBB-LAB/Comparative-analysis-of-plant-TFBS-software

### Competing Interest Statement

The authors have declared no competing interest.

### Summary of Updates

This version contains revised text, figures and tables.

