## Supplementary Material S1 for "Comprehensive analysis of computational approaches in plant transcription factors binding regions discovery"

**Tables**

**Table 1:** List of experimental methods to identify transcription factors binding regions.

| **S. No.** | **Experimental method** | **Description** | **In vivo or**  **in vitro** | **DNA ligand** | **Unique features** | **Year** |
| --- | --- | --- | --- | --- | --- | --- |
| **1** | **ChIP (Chromatin immunoprecipitation)-ChIP [1]** | Chromatin immunoprecipitation (ChIP) and microarray technology recognizes transcription factor (TF) or other DNA-binding protein binding sites throughout the genome. | *In vivo* | Genomic | Optimized cross-linking, sonication, and antibody selection | 2000 |
| **2** | **PBM (protein binding microarrays) and variants [2]** | Fluorescent labeled antibodies that are specific to the TF are utilized to find the protein-DNA interactions directly to double-stranded DNA microarrays. | *In vitro* | Synthetic and randomized | High-throughput | 2004 |
| **3** | **ChIP-seq (chromatin immunoprecipitation followed by sequencing) and variants [3]** | An effective method for mapping genome-wide transcription factor binding sites (TFBSs) *in vivo* by combining ChIP with next-generation sequencing (NGS). | *In vivo* | Genomic | High-throughput and most widely used protocol for TFBS mapping in vivo Endogenous TF or fused with an epitope tag | 2007 |
| **4** | **SELEX-seq (systematic evolution of ligands by exponential enrichment followed by sequencing) [4]** | It is a powerful method used to generate high-affinity nucleic acid ligands, such as aptamers with high affinity and specificity for a wide range of targets screened by random single-stranded nucleic acid sequences. | *In vitro* | Synthetic and randomized | The versatility and evolutionary nature of SELEX make it a valuable tool for generating novel nucleic acid ligands. | 2007 |
| **5** | **ChIP-exo [5]** | Optimizes the resolution and specificity of protein-DNA binding site identification through the combination of chromatin immunoprecipitation with exonuclease digestion. | *In vivo* | Genomic | Employed for studying the interactions between distinct DNA-binding proteins, such as TFs as well as additional regulatory proteins, and proteins within different species. | 2011 |
| **6** | **ChIPmentation [6]** | Automates and enhances the ChIP-seq workflow by integrating sequencing-compatible adaptors into the chromatin immunoprecipitation step by step. | *In vivo* | Genomic | Avoids adaptor dimers sequencing which become apparent when adaptors self-ligate without DNA fragments are less frequent. | 2015 |
| **7** | **ChIP-nexus (Chromatin Immunoprecipitation followed by Nucleotide-resolution Exonuclease digestion and Sequencing) [7]** | A more sophisticated version of the ChIP-seq method, known as, offers higher resolution and accuracy for mapping protein-DNA interactions. It incorporates high-throughput sequencing, exonuclease digestion, and ChIP. | *In vivo* | Genomic | Provides a thorough perspective of the *in vivo* binding landscape of TFs by achieving higher resolution compared to standard ChIP-seq and enhanced robustness compared to ChIP-exo. | 2015 |
| **8** | **(amp)DAP-seq (DNA affinity purification followed by sequencing) [8]** | Recombinant TFs are used in the DAP-seq method to affinity-purify genomic DNA fragments, which are then submitted to NGS in order to determine the cistrome, or collection of genomic binding sites, of the TF; A PCR amplification step is introduced to the AmpDAP-seq methodology, which modifies the DAP-seq method by eliminating DNA methylation patterns prior to affinity purification. This modification enables it possible to identify the binding sites for TFs, including certain plant TFs, that prefer to bind to unmethylated DNA. | *In vitro* | Genomic | Enables the development of cistrome and epicistrome maps for hundreds of TFs of an organism at minimal cost and with great throughput. | 2016 |

**Table 2:** List of peak calling software for ChIP-seq data analysis.

| **S.No.** | **Tool** | **Algorithm** | **Approach** | **Published year** | **Resolution** | **Use of variability of local signal** | **Software applicability** |
| --- | --- | --- | --- | --- | --- | --- | --- |
| **1** | **SISSRs (Site Identification from Short Sequence Reads) [9]** | Site Identification from Short Sequence Reads (SISSRs) algorithm | Sliding window | 2008 | Yes | No | Yes |
| **2** | **USEQ [10]** | Collection of algorithms and software for peak calling | Implemented several different methods for peak calling | 2008 | No | No | N/A |
| **3** | **CisGenome [11]** | Two-pass algorithm | Implemented a modular design, use sliding window for peak detection | 2008 | No | Yes | Yes |
| **4** | **F-Seq [12]** | F-Seq density estimation algorithm | Kernel density estimation | 2008 | Yes | Yes | Yes |
| **5** | **FindPeaks [13]** | Used directional reads module for identifying peaks | Implemented a modular architecture | 2008 | No | No | N/A |
| **6** | **QuEST (Quantitative Enrichment of Sequence Tags) [14]** | Construct profiles and use shifting method | Statistical framework-Kernel Density Estimation approach | 2008 | Yes | Yes | N/A |
| **7** | **MACS (Model-based analysis**  **of ChIP-Seq) [15]** | MACS algorithm (use shift and sliding window algorithm) | Model-based Analysis of ChIP-Seq | 2008 | Yes | Yes | Yes |
| **8** | **GLITR (GLobal Identifier of Target Regions) [16]** | GLITR algorithm | Used ChIP-Seq Peak Finder framework | 2009 | No | No | N/A |
| **9** | **PeakSeq [17]** | Two-pass strategy | Two-pass strategy | 2009 | Yes | Yes | Yes |
| **10** | **SICER [18]** | Scoring scheme | Spatial clustering approach | 2009 | No | Yes | Yes |
| **11** | **SIPeS (Site Identification from Paired-end Sequencing) [19]** | SIPeS algorithm | Used dynamic fragment pileup value for peak calling | 2010 | Yes | Yes | N/A |
| **12** | **Sole-Search [20]** | Sole-Search program | Implemented several different analysis steps for peak calling | 2010 | No | Yes | N/A |
| **13** | **Hpeak (Hidden Markov model (HMM)-based Peak-finding algorithm) [21]** | HMM-based algorithm | Hidden Markov Model (HMM) | 2010 | No | Yes | N/A |
| **14** | **BayesPeak [22]** | BayesPeak algorithm | Used Hidden Markov model (HMM) for finding peaks | 2011 | No | Yes | N/A |
| **15** | **PeakRanger [23]** | Same algorithm as PeakSeq for identifying broad regions. Summit-valley-alternator algorithm | Build the read coverage profile | 2011 | Yes | Yes | Yes |
| **16** | **SeqSite [24]** | Two-step strategy: detect tag-enriched regions and then pinpoint binding sites in the detected regions | Poisson model | 2011 | Yes | Yes | N/A |
| **17** | **T-PIC (Tree shape Peak Identification for ChIP-Seq) [25]** | Tree shape Peak Identification for ChIP-Seq (T-PIC) algorithm | Tree-based statistics | 2011 | No | No | Yes |
| **18** | **W-ChIPeaks [10]** | PELT algorithm and BELT algorithm | Statistical methods control false discovery rate | 2011 | No | No | N/A |
| **19** | **ZINBA (Zero-Inflated Negative Binomial Algorithm) [26]** | Zero-Inflated Negative Binomial Algorithm (ZINBA) | Statistical framework | 2011 | No | Yes | N/A |
| **20** | **GEM (Genome wide Event finding and Motif discovery) [27]** | Genome wide event finding and motif discovery (GEM) | Probabilistic model | 2012 | Yes | Yes | Yes |
| **21** | **GLMNB (Negative binomial generalized linear model) [28]** | Sliding window | Generalized Linear Model with Negative binomial distribution | 2012 | Yes | Yes | Yes |
| **22** | **DROMPA (DRaw and observe Multiple enrichment profiles and annotation) [29]** | Sliding window | Two-step procedure, DROMPA peak-calling program | 2013 | No | No | N/A |
| **23** | **NEXT-peak (the normal-exponential two-peak) [30]** | NEXT-peak algorithm | Normal-exponential two-peak (NEXT-peak) model | 2013 | Yes | No | Yes |
| **24** | **BroadPeak [31]** | Maximal-segment algorithm, Gibbs sampling algorithm, Ruzzo-Tompa algorithm | Probabilistic model | 2013 | No | No | N/A |
| **25** | **PePr [32]** | Sliding window approach | Negative binomial distribution | 2014 | Yes | Yes | Yes |
| **26** | **MUSIC [33]** | Multiscale decomposition, and the multi-mappability profile | Linear regression and Poisson distribution model | 2014 | Yes | No | Yes |
| **27** | **Ritornello [34]** | Deconvolution of multi-binding events | digital signal processing (DSP) and statistical techniques | 2017 | No | Yes | Yes |
| **28** | **LanceOtron [35]** | Peak scoring algorithm | Deep learning (CNN) | 2022 | Yes | YEs | Yes |

**Table 3.** List of some published tools for transcription factor binding regions identification.

| **S. No.** | **Software** | **Year** | **Species** | **Algorithm** | **Dataset** |
| --- | --- | --- | --- | --- | --- |
| **1** | **Ppromotif [36]** | 2014 | *Arabidopsis thaliana* | Probabilistic modeling | AGRIS database |
| **2** | **DeepBind [37]** | 2015 | Human | CNN | ChIP-seq |
| **3** | **DeepSEA [38]** | 2015 | Human | CNN | ChIP-seq |
| **4** | **gkmSVM [39]** | 2016 | Human | SVM | ChIP-seq |
| **5** | **Basset [40]** | 2016 | Human | CNN | DNase-seq |
| **6** | **DanQ [41]** | 2016 | Human | CNN and Bi-LSTM | ChIP-seq, DNase-seq |
| **7** | **LS-GKM [42]** | 2016 | Human | SVM | ChIP-seq |
| **8** | **DeeperBind [43]** | 2016 | Human | CNN+LSTM | PBMs |
| **9** | **DeepSNR [44]** | 2017 | Human | CNN and DeepCNN | ChIP-exonuclease |
| **10** | **Dilated [45]** | 2017 | Human | Dilated CNN | ChIP-seq |
| **11** | **TFimpute [46]** | 2017 | Human | CNN | ChIP-seq |
| **12** | **KEGRU [47]** | 2018 | Human | Bi-GRU | ChIP-seq |
| **13** | **DeFine [48]** | 2018 | Human | CNN | ChIP-seq |
| **14** | ***k*-mer grammar [49]** | 2019 | *Zea mays* | bag-of-*k*-mers | ChIP-seq, MNA-seq |
| **15** | **FactorNet [50]** | 2019 | Human | CNN+BiLSTM | ChIP-seq |
| **16** | **DESSO [51]** | 2019 | Human | CNN | ChIP-seq |
| **17** | **WSCNNLSTM [52]** | 2019 | Human | Multi-instance learning and hybrid neural network | ChIP-seq |
| **18** | **deepRAM [53]** | 2019 | Human | CNN/RNN | ChIP-seq |
| **19** | **DeepTF [54]** | 2019 | Human | CNN+LSTM | ChIP-seq |
| **20** | **TbiNet [55]** | 2020 | Human | CNN, Bi-LSTM, and Attention mechanism | ChIP-seq |
| **21** | **FCNA [56]** | 2021 | Human | CNN | ChIP-seq |
| **22** | **DLBSS [57]** | 2021 | Human | CNN/RNN | PBM |
| **23** | **BPNet [58]** | 2021 | Human | CNN | ChIP–nexus |
| **24** | **SAResNet [59]** | 2021 | Human | Self-attention mechanism+residual network | ChIP-seq |
| **25** | **DNABERT [60]** | 2021 | Human | Transformer | ChIP-seq |
| **26** | **SeqConv [61]** | 2021 | *Zea mays* | CNN | ChIP-seq |
| **27** | **D-AEDNet [62]** | 2021 | Human | Deep Attentive Encoder-Decoder Neural Network | ChIP-exo and ChIP-seq |
| **28** | **TSPTFBS [63]** | 2021 | *Arabidopsis thaliana* | CNN | DAP-seq |
| **29** | **AgentBind [64]** | 2021 | Human | CNN, CNN+RNN | ChIP-seq |
| **30** | **D-SSCA [65]** | 2022 | Human | CNN+ attentation mechanism | ChIP-seq |
| **31** | **MAResNet [66]** | 2022 | Human | Top-down and bottom-up attentation mechanism+residual network | ChIP-seq |
| **32** | **FCNGRU [67]** | 2022 | Human | CNN+GRU | PBM |
| **33** | **Wimtrap [68]** | 2022 | *Arabidopsis thaliana* | XGBoost | ChIP-seq |
| **34** | **PlantBind [69]** | 2022 | *Arabidopsis thaliana* | CNN+Bi-LSTM | ChIP-seq |
| **35** | **FCNsignal [70]** | 2022 | Human | Encoder + Decoder + Skip Architecture | ChIP-seq and ATAC-seq |
| **36** | **TSPTFBS 2.0 [71]** | 2023 | *Arabidopsis thaliana* | DenseNet | ChIP-seq and DAP-seq |
| **37** | **DeepSTF [72]** | 2023 | Human | CNN, transformer encoder, and Bi-LSTM | ChIP-seq |
| **38** | **PTFSpot [73]** | 2024 | Plants | DenseNet + Encoder | ChIP-seq and DAP-seq |

**Table 4:** List of databases contain information on transcription factors.

| **S. No.** | **Database** | **Description** | **Year** | **Link** |
| --- | --- | --- | --- | --- |
| **1** | **TRANSFAC [74]** | A database which provides details on eukaryotic TFs, their binding locations, and the regulation of transcription. It presents information on regulatory networks, PWMs (position weight matrices), and TF binding sites. It covers TFs from a wide range of taxa, including animal and plants species. | 1996 | https://genexplain.com/transfac/ |
| **2** | **AGRIS [75]** | *Arabidopsis* Gene Regulatory Information Server (AGRIS) is a repository including TFs and gene regulatory networks in *Arabidopsis thaliana* It contains comprehensive information on the protein sequences, expression data, cis-regulatory components, and functional annotations of *Arabidopsis* TF families. | 2003 | https://agris-knowledgebase.org/ |
| **3** | **ENCODE [76]** | The Encyclopedia of DNA Elements (ENCODE) project contains abundance of information on TF binding sites and other regulatory components obtained from the human genome. It provides a comprehensive collection of ChIP-seq data, TF binding profile, and other functional annotations. | 2003 | https://www.encodeproject.org/ |
| **4** | **JASPAR [77]** | A widely used database for TF binding profiles and motifs. It contains curated TF binding site profiles for both animals, plants and several other species. | 2004 | https://jaspar.genereg.net/ |
| **5** | **PlantTFDB [78]** | An extensive database containing information on TFs for various plant species. Their phylogenetic analyses, functional annotations, protein sequences, gene expression data, and curated TF families are all covered. | 2008 | http://planttfdb.cbi.pku.edu.cn/ |
| **6** | **AnimalTFDB [79]** | Animal Transcription Factor Database contains detailed  annotations for each TF, including basic information, gene structure, functional domain, 3D structure  hit, Gene Ontology, pathway, protein–protein interaction, paralogs, orthologs, potential TF-binding sites and targets | 2012 | https://ngdc.cncb.ac.cn/databasecommons/database/id/8 |
| **7** | **Plant Cistrome database [80]** | The database serves as a valuable resource for plant researchers interested in transcriptional regulation and TF-DNA interactions. It provides a user-friendly interface, data visualization tools, and curated information that can aid in the exploration of plant cistromes and the discovery of novel regulatory elements. | 2016 | http://neomorph.salk.edu/PlantCistromeDB |
| **8** | **HOCOMOCO [81]** | The Human and Mouse Consensus database is a repository which contains profiles and motifs for both mouse and human TF binding sites. It contains insights on DNA motifs, gene regulation networks, and TF binding preferences. | 2018 | http://hocomoco11.autosome.ru/ |
| **9** | **PlantRegMap [82]** | Plant Regulatory Map is a database that combines regulatory interactions between TFs, their target genes, and multiple species of plants. For investigating gene regulation in plants, the repository provides detailed regulatory maps and network analysis tools. | 2020 | http://plantregmap.gao-lab.org/ |
