## Supplementary Material S2 for "Comprehensive analysis of computational approaches in plant transcription factors binding regions discovery"

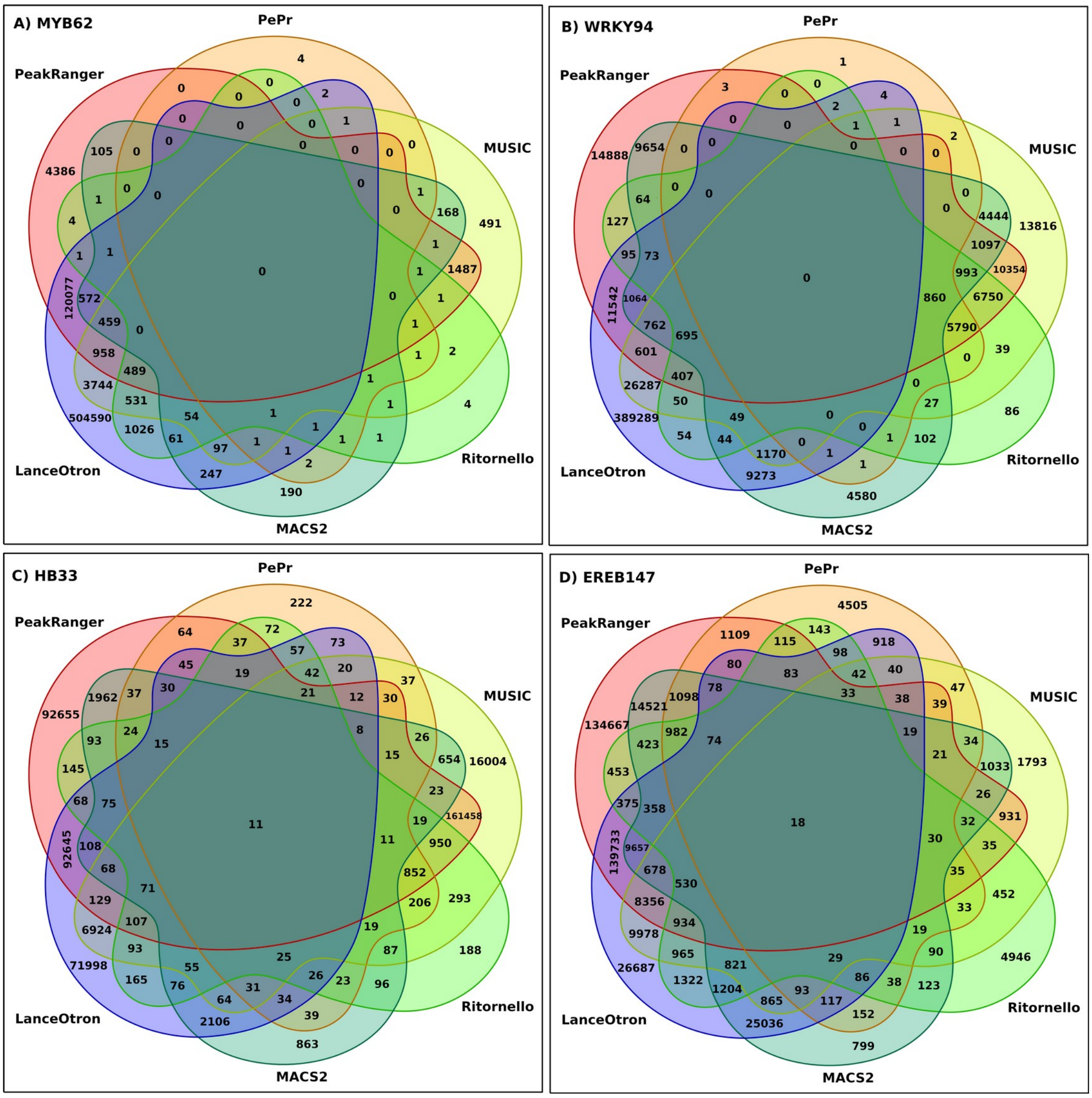


**Supplementary Figure 1: Number of peaks overlap in MYB62, WRKY94, HB33, and EREB147 TFs binding regions between six software tools:** For four transcription factors (TFs), the unique and common peaks between pairs, trios, quartets, and all six software tools are shown.

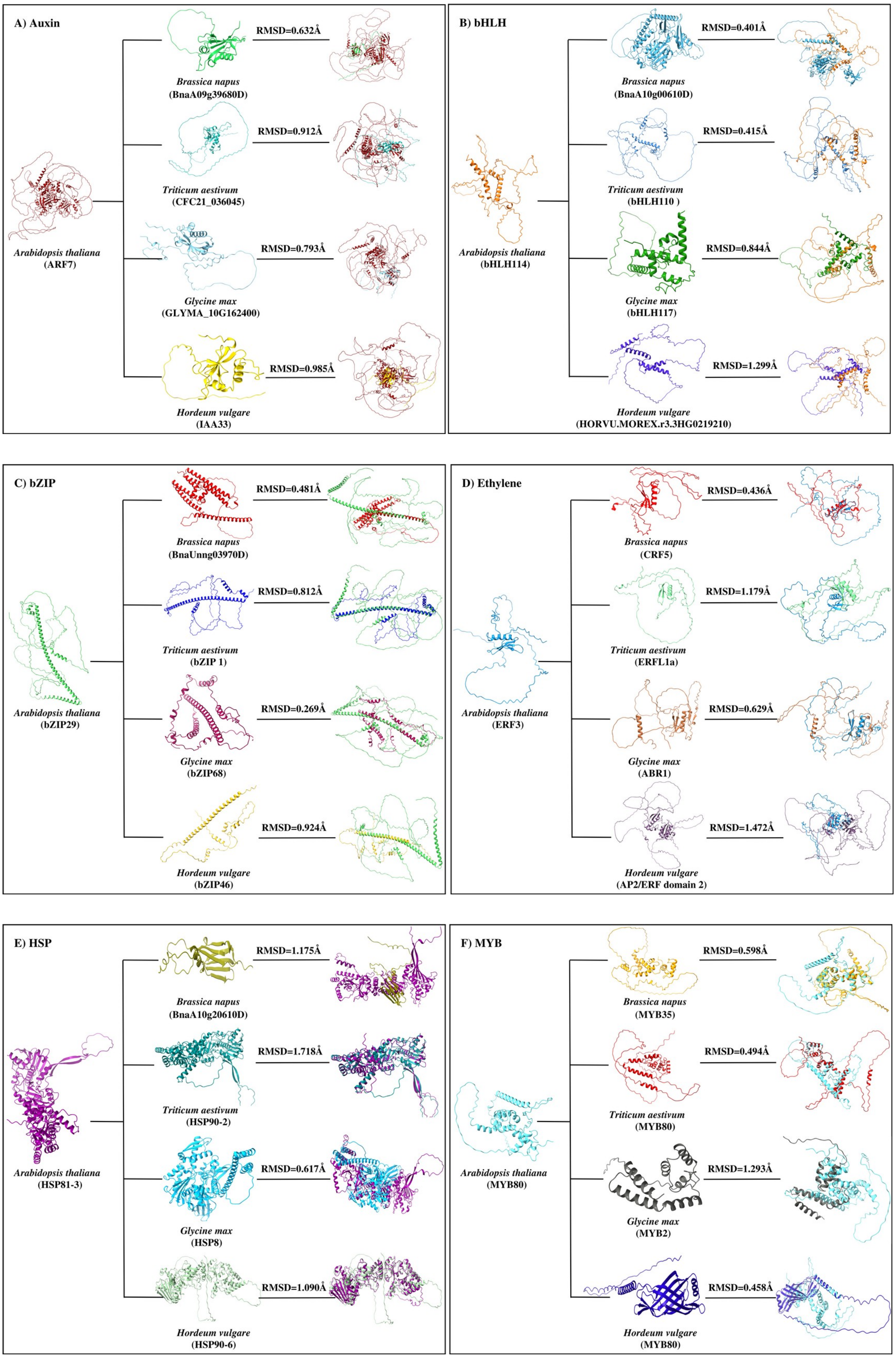

A) Auxin
RMSD=0.632Å
Brassica napus
(BnaA09g39680D)
RMSD=0.912Å
Triticum aestivum
(CFC21_036045)
Arabidopsis thaliana
(ARF7)
RMSD=0.793Å
Glycine max
(GLYMA_10G162400)
RMSD=0.985Å
Hordeum vulgare
(IAA33)
B) bHLH
RMSD=0.401Å
Brassica napus
(BnaA10g00610D)
RMSD=0.415Å
Triticum aestivum
(bHLH110 )
Arabidopsis thaliana
(bHLH114)
RMSD=0.844Å
Glycine max
(bHLH117)
RMSD=1.299Å
Hordeum vulgare
(HORVU.MOREX.r3.3HG0219210)
C) bZIP
RMSD=0.481Å
Brassica napus
(BnaUnng03970D)
RMSD=0.812Å
Triticum aestivum
(bZIP 1)
Arabidopsis thaliana
(bZIP29)
RMSD=0.269Å
Glycine max
(bZIP68)
RMSD=0.924Å
Hordeum vulgare
(bZIP46)
D) Ethylene
RMSD=0.436Å
Brassica napus
(CRF5)
RMSD=1.179Å
Triticum aestivum
(ERFL1a)
Arabidopsis thaliana
(ERF3)
RMSD=0.629Å
Glycine max
(ABR1)
RMSD=1.472Å
Hordeum vulgare
(AP2/ERF domain 2)
E) HSP
RMSD=1.175Å
Brassica napus
(BnaA10g20610D)
RMSD=1.718Å
Triticum aestivum
(HSP90-2)
Arabidopsis thaliana
(HSP81-3)
RMSD=0.617Å
Glycine max
(HSP8)
RMSD=1.090Å
Hordeum vulgare
(HSP90-6)
F) MYB
RMSD=0.598Å
Brassica napus
(MYB35)
RMSD=0.494Å
Triticum aestivum
(MYB80)
Arabidopsis thaliana
(MYB80)
RMSD=1.293Å
Glycine max
(MYB2)
RMSD=0.458Å
Hordeum vulgare
(MYB80)

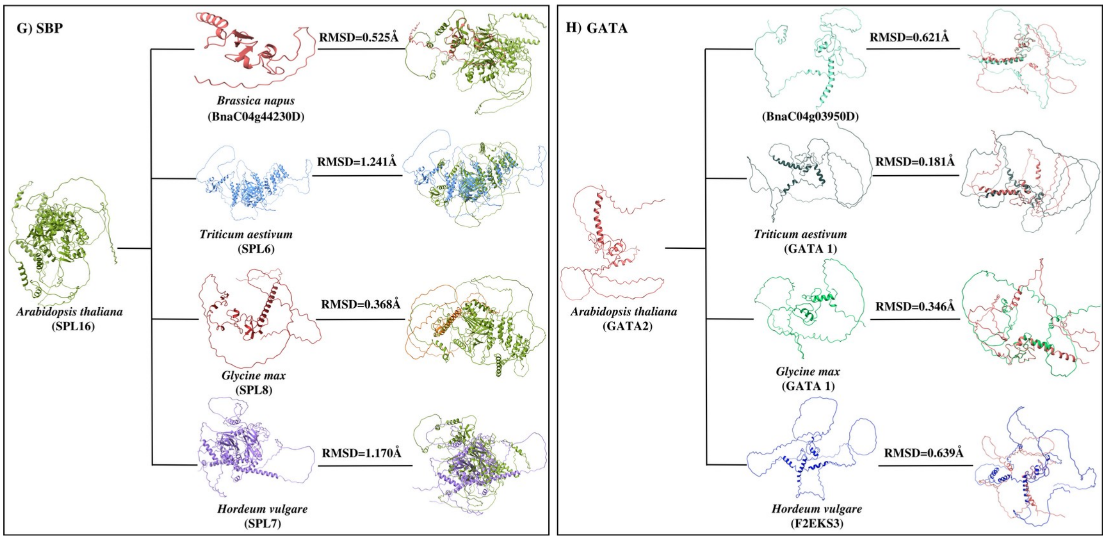


**Supplementary Figure 2: 3D structure based co-variation in transcription factors families' structure across five plant species (*A. thaliana* vs *Brassica napus*, *A. thaliana* vs *Triticum aestivum*, *A. thaliana* vs *Glycine max*, *A. thaliana* vs *Hordeum vulgare*).** (A), and (B) The corresponding superimposed TF structures with the RMSD value indicating the structural variations across TF families, named Auxin and bHLH, respectively. (C), and (D) The corresponding superimposed TF structures with the RMSD value indicating the structural variations across TF families, named bZIP and ethylene, respectively. (E), and (F) The corresponding superimposed TF structures with the RMSD value indicating the structural variations across TF families, named HSP and MYB, respectively. (G), and (H) The corresponding superimposed TF structures with the RMSD value indicating the structural variations across TF families, named SBP and GATA, respectively.

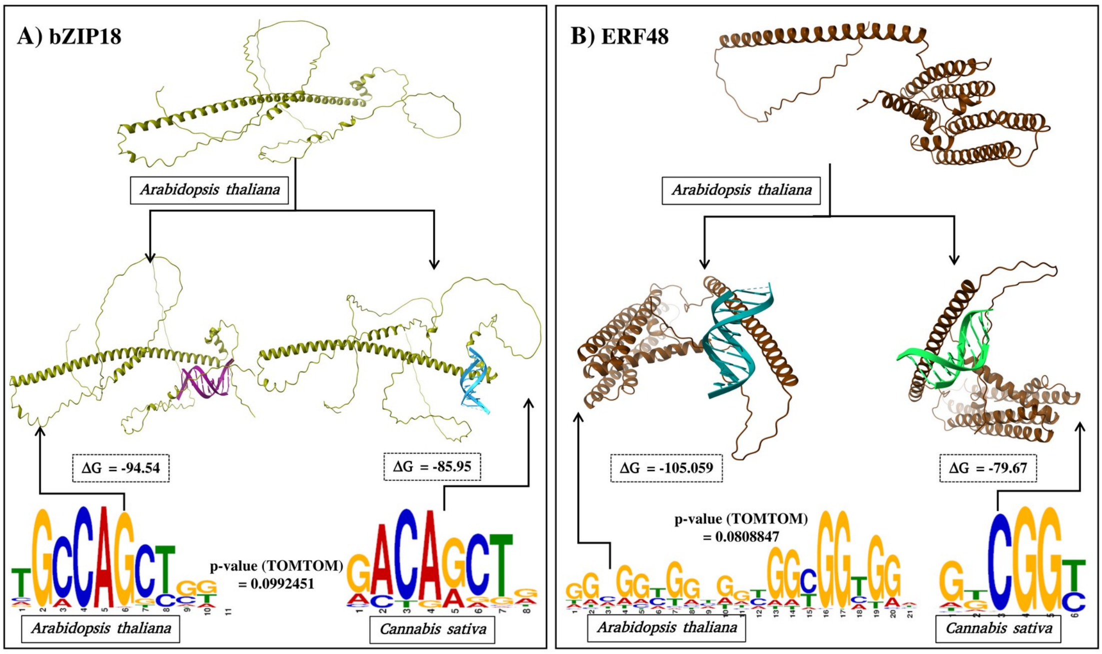

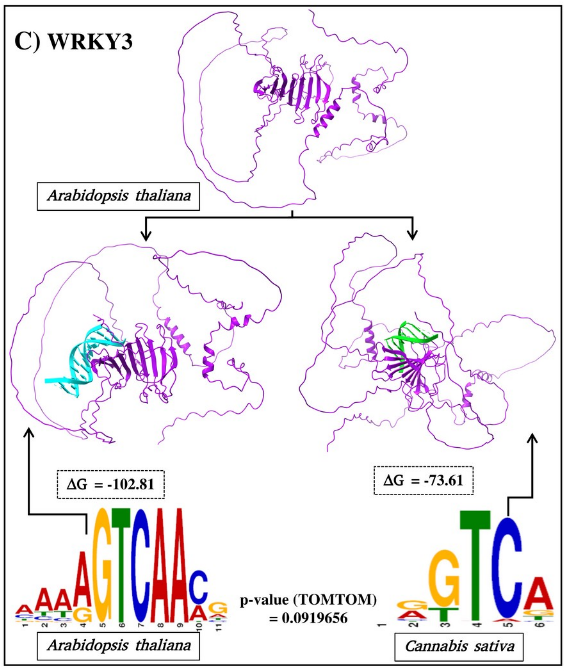


**Supplementary Figure 3:** Co-variation in the structure of the transcription factors and their corresponding binding sites across *A. thaliana* and *Cannabis sativa* species. TF structure and its binding motif comparison with experimental validated binding motif of (A) bZIP18, (B) ERF48, and (C) WRKY3 TFs and its 3D structure. The docking analysis shows the stability of the complexes when a TF was docked to its binding motif within the same species and when docked with other species for the same TF, there was less stability of the complexes. It is clearly evident that the same binding motif can not work across the species and it varies with species as well as the structure of the TF.

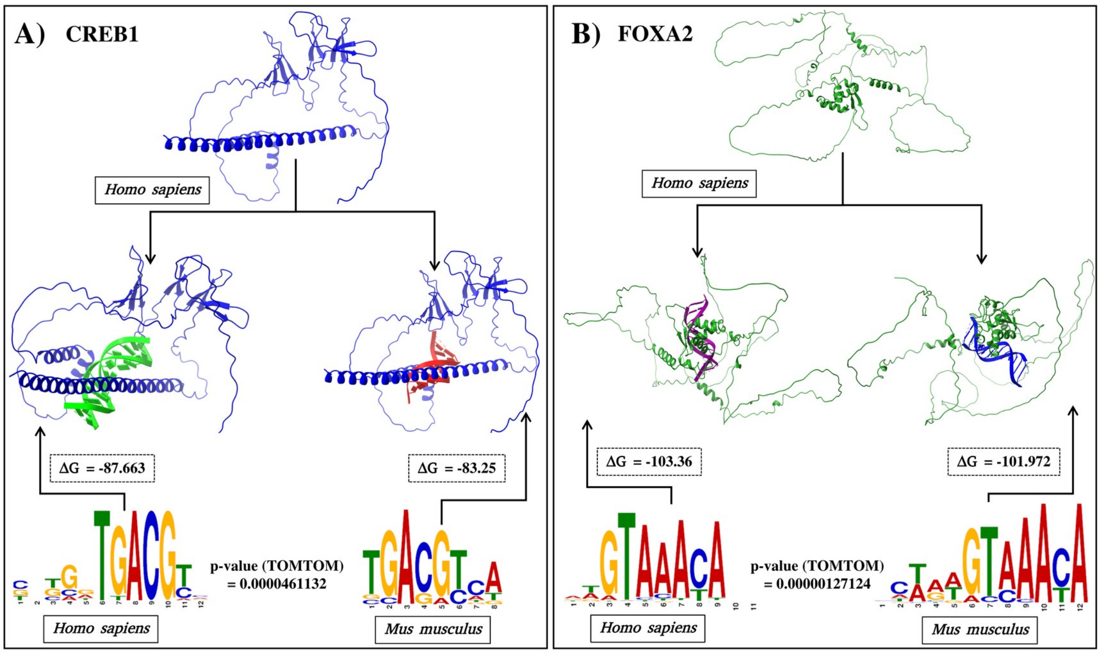

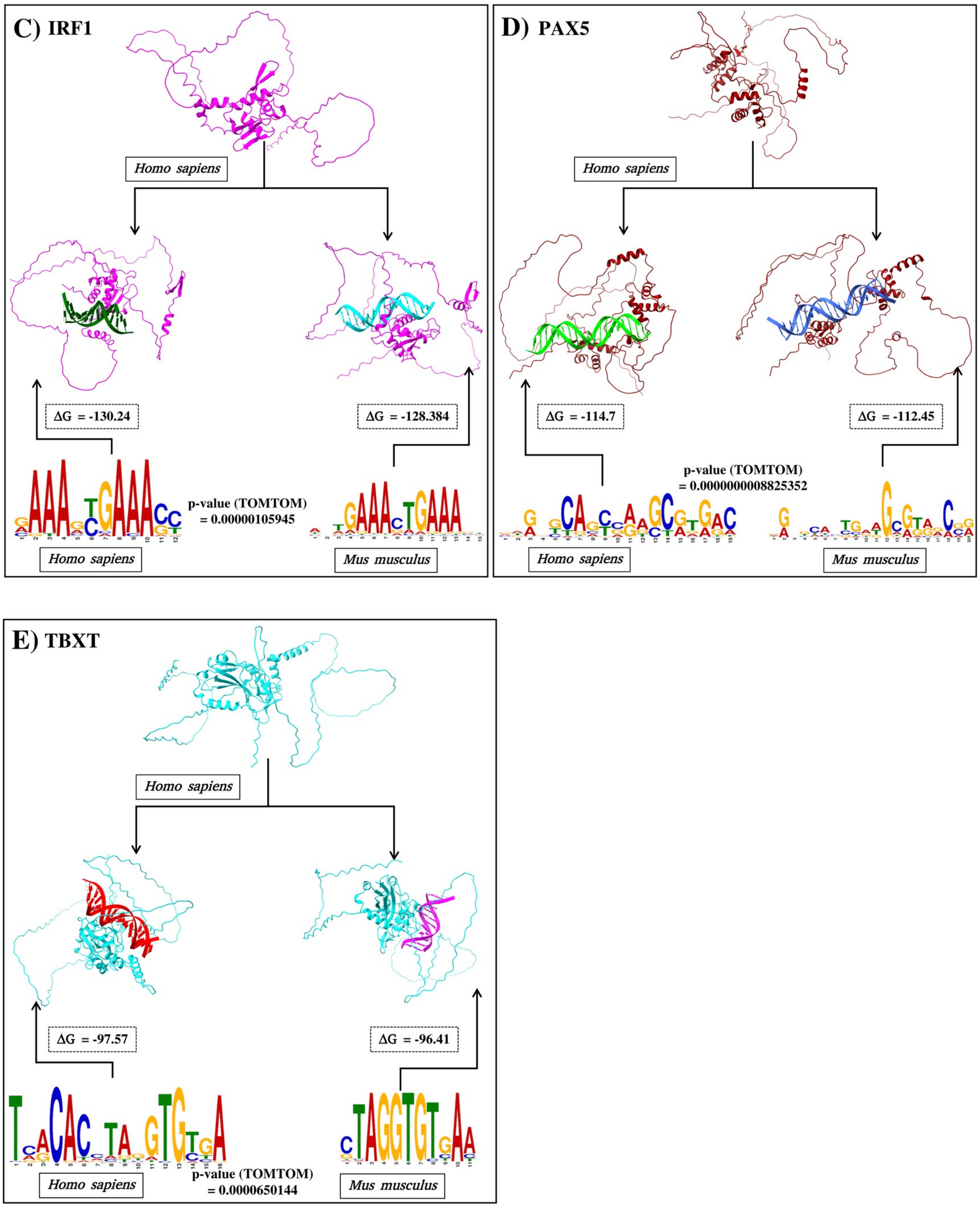


**Supplementary Figure 4:** Co-variation in the structure of the transcription factors and their corresponding binding sites across *Homo sapiens* and *Mus musculus* species. TF structure and its binding motif comparison with experimental validated binding motif of (A) CREB1, (B) FOXA2, (C) IRF1, (D) PAX5, and (C) TBXT TFs and its 3D structure. The docking analysis shows the stability of the complexes when a TF was docked to its binding motif within the same species and when docked with other species for the same TF, there was more stability of the complexes. It is clearly evident that the same binding motif can easily work across other species as the comparative motifs have a significant similar motifs as well as the structure of the TF.
