## Supplementary Material S3 for "Comprehensive analysis of computational approaches in plant transcription factors binding regions discovery"

### **Supplementary File**

**Error analysis and debugging**

**NOTE:** Before executing a machine learning (ML) or deep learning (DL) program, take into consideration the following instructions in order to minimize any inconvenience:

1. It is essential to ensure that your input data is properly formatted, labeled, and in the expected format to ensure smooth execution and obtain meaningful results. It is advised to visit our GitHub page at https://github.com/SCBB-LAB/Comparative-analysis-of-plant-TFBS-software and have insight into the input data formats, written in detailed description in README files of all the software, so that you can reduce inconveniences and make your ML and DL program more manageable.

2. Second, to use GPU acceleration, assure that you have all the necessary software and dependencies installed, notably Python, the appropriate libraries (such as TensorFlow, PyTorch, scikit-learn, keras, etc), and any specific GPU drivers or CUDA libraries.

3. To keep track of dependencies in order to avoid incompatibilities with other projects, create a virtual environment for the project you are working on, which is described in our GitHub README file. When you are dealing with multiple projects or programs that each have different package prerequisites and you want to ensure that each project has its own isolated environment with the necessary dependencies. Therefore, activating “Conda” enviornment can be exceptionally advantageous. It facilitates improved project management, organization, and reliability for your Python-based programs.

Now, we capable of what is listed below in order to identify and resolve errors in Python codes for each and every software.

============================================================

**1. DeepBind**

**Algorithm:** Convolution neural network (CNN)

**Resource type:** Standalone package

**See URL:** https://github.com/jisraeli/DeepBind

| **Dependencies** | **Script** | **Bugs** | **Problem** | **Solution** |
| --- | --- | --- | --- | --- |
| Python,  CUDA 7.0+  Visual C++ 2012 / GCC 4.8+ | /home/user/DeepBind/code/libs/smat/src$ make | make: *** [makefile:230: ../build/release/obj/SMAT_CUDA/reduce.o] Error 127 | CUDA compatiblity error | ‘DeepBind-with-Pytorch’ forked from ‘jisraeli/DeepBind’ |

**See URL:** https://github.com/MedChaabane/DeepBind-with-PyTorch forked from jisraeli/**DeepBind**

Therefore, the forked DeepBind model was implemented in PyTorch in above listed repository. Using PyTorch, they applied the model from the article titled “Predicting the sequence specificity of DNA and RNA-binding proteins by deep learning” into the practice. The publication additionally provides supplementary notes that explore the architecture in great detail.

**Resource type:** Standalone package

| **Dependencies** | **Script** | **Bugs** | **Problem** | **Solution** |
| --- | --- | --- | --- | --- |
| Python3, PyTorch | /home/user/DeepBind/deepbind.py | N/A | N/A | N/A |
|  |  | N/A | No performance metrics computation lines in program | A whole bunch of lines were added to calculate evaluation metrics |

**Command line:**

To execute the program, run the following command line in the Linux terminal:

- python3 deepbind.py deepbind_data/TF_name

**NOTE:** Here, “TF_name” refers to the specific TF file name, which could be either in FASTA or bed file format for the input sequences. For example, it could be for example “ABF2”. This naming convention will be consistent across all the methods detailed below.

**Output:**

The trained model and the best hyperparameters, named “TF_name_Model.pth” and “TF_name_best_hyperparameters.pth” respectively, are saved to the output location in the “model/” directory. Additionally, the performance metrics for model training on the test data are also located in the “model/” directory, specifically in the file named “TF_name_result.txt”.

**============================================================**

**2. gkmSVM**

**Algorithm:** Support Vector Machine (SVM)

**Resource type:** Standalone

**See URL:** https://www.beerlab.org/gkmsvm/gkmsvm-tutorial.html

| **Dependencies** | **Script** | **Bugs** | **Problem** | **Solution** |
| --- | --- | --- | --- | --- |
| R version 3.5 or later |  | Error in install.packages("ROCR", "kernlab", "seqinr") :  unable to install packages | Package installation error | Installed these packages individually,  “install.packages('ROCR')  install.packages('kernlab')  install.packages('seqinr')  install.packages('gkmSVM')” |
|  | 1.R | N/A | No script for performance metrics calculation | Provided a python program to calculate all the performance metrics, ‘evaluate.py’ in parant ‘gkmSVM’ directory. |

**Commands line:**

To enter the R environment, simply type “R” in the terminal. Then, you can run the following commands for your particular TF:

- library(gkmSVM)
- gkmsvm_kernel('example/TF_name_pos.fa','example/TF_name_neg.fa', 'output/TF_name_kernel_out')
- gkmsvm_trainCV('output/TF_name_kernel_out','example/TF_name_pos.fa','example/TF_name_neg.fa',svmfnprfx='output/TF_name', outputCVpredfn='output/TF_name_cvpred.out', outputROCfn='output/TF_name_roc.out', outputPDFfn = 'output/TF_name_ROC2.pdf')

**Output:**

To get the prediction scores of positive and negative TFBS files, run the following shell script lines in your terminal:

- awk '{print $3"\t"$2}' output/TF_name_cvpred.out | grep "^1" > output/TF_name_pos_gkmpredict.txt # prediction score for positive dataset
- awk '{print $3"\t"$2}' output/TF_name_cvpred.out | grep "^-1" > output/TF_name_neg_gkmpredict.txt # prediction score for negative dataset
- python3 evaluate.py -p output/TF_name

The prediction output files are saved in “output/” directory with “TF_name_cvpred.out” extension. Keep the “evaluate.py” python program in the same directory. Replace “TF_name” with the name of your TF to execute your program. Make sure you're in the correct directory where your program is located before running this command

**Output:** This will output all the performance metrics such as accuracy, MCC-values, F1 score, etc., in the “output/TF_name_result.txt” file located in the “result/” directory.

**============================================================**

**3. LS-GKM**

**Algorithm:** Support vector machine (SVM)

**Resource type:** Standalone

**See URL:** https://github.com/Dongwon-Lee/lsgkm

| **Dependencies** | **Script** | **Bugs** | **Problem** | **Solution** |
| --- | --- | --- | --- | --- |
| C/C++ | src/gkmtrain  src/gkmpredict | N/A | N/A | N/A |
|  |  |  | No script for performance metrics calculation | Provided a python program to calculate all the performance metrics, ‘evaluate.py’ in parant ‘gkmSVM’ directory. |

**Command line:**

To perform model training, prediction, and evaluation for TFBSs identification, run the following command line in the Linux terminal:

- bin/gkmtrain /user_path/lsgkm/example/TF_name_pos_train.fa /user_path/lsgkm/example/TF_name_neg_train.fa /user_path/example/result/TF_name_gkmtrain
- bin/gkmpredict /user_path/lsgkm/example/TF_name_pos_test.fa /user_path/lsgkm/output/TF_name_train.model.txt /user_path/lsgkm/output/TF_name_gkmpredict_pos.txt
- bin/gkmpredict /user_path/lsgkm/example/TF_name_neg_test.fa /user_path/lsgkm/output/TF_name_train.model.txt /user_path/lsgkm/output/TF_name_gkmpredict_neg.txt
- python3 scripts/evaluate.py -p output/TF_name

Replace “TF_name” with the name of your TF to execute your program. Make sure you're in the correct directory where your program is located before running this command

**Output:**

This will output all the performance metrics such as accuracy, MCC-values, F1 score, etc., in the “output/TF_name_result.txt” file located in the “result/” directory.

**============================================================**

**4. KEGRU**

**Algorithm:** Recurrent neural network (Bi-GRU)

**Resource type:** Source code and data are not publicly available

**Solution:** Source code was provided by co-author

| **Dependencies** | **Script** | **Bugs** | **Problem** | **Solution** |
| --- | --- | --- | --- | --- |
| Python3,  keras based architecture | home/user/kegru/dbrmodel7.py | line 58 | SyntaxError: Missing parentheses in call to 'print'. Did you mean print(...)? | Script changed from python2.7 to python3 version by “2to3 dbrmodel7.py -W” |
|  |  |  | ImportError: cannot import name 'pad_sequences' from 'keras.preprocessing.sequence' | From: “keras.preprocessing.sequence import pad_sequences”  To:  “from keras_preprocessing.sequence import pad_sequences” |
|  |  |  | ModuleNotFoundError: No module named 'keras_preprocessing' | conda install -c conda-forge keras-preprocessing |
|  |  |  | ModuleNotFoundError: No module named 'keras.layers.embeddings' | from keras.layers import Embedding |
|  |  | Lines 63, 64 | FileNotFoundError: [Errno 2] No such file or directory: '/home/szhen/szhen/DBLSTM/test2/init_pos_5gram_2stride' and ‘/home/szhen/szhen/DBLSTM/test2/init_neg_5gram_2stride' | Line63: pos_name = options.name + '_pos.fa'  Line64: neg_name = options.name + '_neg.fa' |
|  |  | Line 81 | TypeError: Word2Vec.__init__() got an unexpected keyword argument 'size' | Use "vector_size" instead of "size" |
|  |  | line 140 | TypeError: int() argument must be a string, a bytes-like object or a real number, not 'NoneType' | model.add(Bidirectional(GRU(int(80), return_sequences=True))) |
|  |  | line 318 | ValueError: Could not interpret optimizer identifier: None | model.compile(loss='binary_crossentropy', optimizer="Adam", metrics=['accuracy']) |
|  |  | line 65 | TypeError: Model.fit() got an unexpected keyword argument 'nb_epoch' | “epochs” not “nb_epoch” |
|  |  | line 513 | tensorflow.python.framework.errors_impl.PermissionDeniedError: /home/szhen; Permission denied | parser.add_option('-p', dest='file_path', default='pathtotrain data file', help='train data file path') |
|  |  | line 181 | KeyError: 'acc', ‘val_acc’ | ‘accuracy’, ‘val_accuracy’ |
|  |  | Line 184 | TypeError: can only concatenate str (not "NoneType") to str | str(curoutn), str(moname) and str(options.out) |
|  |  | Lines from 117 to 120 | embedding matrix was not being updated and resulted into very poor predictions | Added few lines to get *k-*mer in upper case |
|  |  |  | No performance metrics computation coding in program | A whole bunch of lines were added to calculate evaluation metrics |

**Command line:**

To execute the program, run the following command line in the Linux terminal:

- python3 kegru_train.py -n TF_name -p data

**Output:**

The “TF_name_result.txt” text file contain performance metrics for a particular TF in the “result/”directory.

**============================================================**

**5. *k*-mer grammar**

**Algorithm:** Linear regression

**Resource type:** Standalone

**See URL:** https://bitbucket.org/bucklerlab/k-mer_grammar/src/master/

| **Dependencies** | **Script** | **Bugs** | **Problem** | **Solution** |
| --- | --- | --- | --- | --- |
| Python3,  keras based architecture  sqlalchemy=1.4.39 | /home/user/k-mer_grammar/kgrammar_bag-of-k-mer_training_testing.py | Line 38 | ImportError: cannot import name 'joblib' from 'sklearn.externals' (/home/csir/miniconda3/lib/python3.10/site-packages/sklearn/externals/__init__.py) | import joblib |
|  |  | line 26 | ModuleNotFoundError: No module named 'sqlalchemy' | python3 -m pip install sqlalchemy |
|  |  | line 267 | AttributeError: 'TfidfVectorizer' object has no attribute 'get_feature_names'. Did you mean: 'get_feature_names_out'? | vcblry = tmpvectorizer.get_feature_names_out() |
|  |  | Lines from 230 to 237 | Stringent parameters were set for a specific length of the sequences and 5 columns were considered for data input | Added few lines to input sequences of any length and only two columns for specified sequence and label only |
|  |  |  | No performance metrics computation coding in program | A whole bunch of lines were added to calculate evaluation metrics |

**Command line:**

To execute the program, run the following command line in the Linux terminal:

- python3 kgrammar_bag-of-k-mer_training_testing.py 8 False TF_name

Replace “TF_name” with the name of your TF to execute your program. Make sure you're in the correct directory where your program is located before running this command.

**Output**

The resulting model file, named “kgrammar_bag-of-k-mers_LR_mode_full_TF_name_8.pkl”, will be saved to the output directory “output/”. Additionally, a database containing *k*-mer weights will also be included in the output directory. Results of the training, including accuracy and other metrics (such as the confusion matrix), will be saved to “TF_name_result.txt” located at “output/”.

**============================================================**

**6. WSCNNLSTM**

**Algorithm:** Weakly supervised learning framework combining multi-instance learning (MIL)

**Resource type:** Standalone

**See URL:** https://github.com/turningpoint1988/WSCNNLSTM

| **Dependencies** | **Script** | **Bugs** | **Problem** | **Solution** |
| --- | --- | --- | --- | --- |
| Python3,  Keras based architecture | sh encoding.sh | encoding.sh: 9: Syntax error: Bad for loop variable |  | Direct calling of encoding.py command line to terminal |
|  | /home/user/WSCNNLSTM/encoding.py | Line 17 | SyntaxError: Missing parentheses in call to 'print'. Did you mean print(int 'data shape: ',data.shape)? | Script changed from python2.7 to python3 version by “2to3 encoding.py -W” |
|  | /home/user/WSCNNLSTM/train_val_test.py | Line 37 | SyntaxError: Missing parentheses in call to 'print'. Did you mean print('working on %s now' % name)? | Script changed from python2.7 to python3 version by “2to3 train_val_test.py -W” |
|  | /home/user/WSCNNLSTM/models.py | Line 17 | print model.summary()  ^  SyntaxError: invalid syntax | Script changed from python2.7 to python3 version by “2to3 models.py -W” |
|  |  | Line 6 | ModuleNotFoundError: No module named 'ANDNoisy' | Module is imported from Noisy_and.py instead of the wrong module is imported |
|  | /home/user/WSCNNLSTM/Noisy_and.py | Line13 | print "a = {}".format(self.a)  ^  SyntaxError: invalid syntax | Script changed from python2.7 to python3 version by “2to3 Noisy_and.py -W” |
|  |  | Line 4 | ModuleNotFoundError: No module named 'keras.engine.topology' | from tensorflow.python.keras.layers import Layer |
|  | /home/user/WSCNNLSTM/models.py | Line 41 | Hyperparameter value is not set for dropout layer | Added few lines to input sequences of any length |
|  | /home/user/WSCNNLSTM/utils.py | Line 69 |  | Hyperparameter values set for dropout layer |
|  |  | Lines from 70 to 80 | Hyperparameter value not as per article | Hyperparameter Values set as per reported in article |
|  |  | Lines 99, 101, 127 | KeyError: 'val_acc' | Keywords Changed to val_accuracy and accuracy |
|  |  | Lines 126 | KeyError: 'acc' |  |
|  | /home/user/WSCNNLSTM/train_val_test.py | Lines 95, 96, 104 and 105 | KeyError: 'val_acc' | Keywords Changed to val_accuracy |
|  | /home/user/WSCNNLSTM/utils.py | Line 103 | del_num = range(len(history_all): 'range' object has no attribute 'pop' | del_num = list(range(len(history_all))) |
|  | /home/user/WSCNNLSTM/train_val_test.py |  | No performance metrics computation coding in program | A whole bunch of lines were added to calculate evaluation metrics |

**Command line:**

To execute the program, run the following command line in the Linux terminal:

- python3 encoding.py ./example/TF_name/positive.fasta ./example/TF_name/negative.fasta ./example/TF_name/datalabel.hdf5 -m ./mappers/3mer.txt -c 120 -s 10 --no-reverse -kmer 3 -run 'ws'
- python3 train_val_test.py -datalable .example/TF_name/datalabel.hdf5 -k 3 -run 'ws' -batchsize 300 -params 12

Replace “TF_name” with the name of your TF to execute your program. Make sure you're in the correct directory where your program is located before running this command.

**Output:**

The (“datalabel.hdf5”) file is saved in the same location “example/TF_name/” as the input file for the “encoding.py”. All the trained models on *k*-fold cross validation and prediction scores are saved to the output location “model/” directory. Additionally, the performance metrics for model training on the test data (“train_matrics.txt” and “test_matrics.txt”) are also saved in the “model/” directory.

**===========================================================**

**7. deepRAM**

**Algorithm:** CNN, RNN type (BiLSTM)

**Resource type:** Standalone

**See URL:** https://github.com/MedChaabane/deepRAM

| **Dependencies** | **Script** | **Bugs** | **Problem** | **Solution** |
| --- | --- | --- | --- | --- |
| Python3,  Pytorch based archetecture | /home/user/deepRAM/deepRAM.py | Lines 313, 429, 434, 440 and 445 | ValueError: Only one class present in y_true. ROC AUC score is not defined in that case. | Added line to shuffle the positive sample to generate negative samples |
|  |  |  | No performance metrics computation coding in program | A whole bunch of lines were added to calculate evaluation metrics |

**Command line:**

To execute the program, run the following command line in the Linux terminal:

- python3 deepRAM.py --train_data example/TF_name_train.txt --test_data example/TF_name_test.txt --data_type DNA --train True --evaluate_performance True --model_path model/TF_name_model.pkl --out_file model/TF_name_prediction.txt --out_file1 model/TF_name_result.txt --Embedding False --Conv True --RNN False --conv_layers 1

**Output:**

The trained models (“TF_name_model.pkl”) and prediction scores file (“TF_name_prediction.txt”) are saved to the output location in the “model/” directory. Additionally, the performance metrics for model training (“TF_name_result.txt”) are also saved in the “model/” directory.

**============================================================**

**8. DESSO**

**Algorithm:** CNN

**Resource type:** Standalone

**See URL:** https://github.com/OSU-BMBL/DESSO

| **Dependencies** | **Script** | **Bugs** | **Problem** | **Solution** |
| --- | --- | --- | --- | --- |
| Python3,  Keras based architecture | /home/user/DESSO/code/train.py | Line 68, 82 | ‘train’, ‘test’ for generating negative dataset for as per methods requirement | However, in line 121 of util.py, to provide negative dataset by the user ‘all’ statement is considered |
|  | /home/user/DESSO/code/libs/training.py |  | No performance metrics computation coding in program | A whole bunch of lines were added to calculate evaluation metrics |

**Command line:**

To execute the program, run the following command line in the Linux terminal:

- python3 train.py --start_index 0 --end_index 1 --peak_flank 100 --network CNN --feature_format Seq

**Note:** The argument “--peak_flank” is used to specify the length of input sequences. It can be adjusted to 50 for sequences of 100 base pairs in length, 100 for sequences of 200 base pairs, and vice versa.

**Output:**

For the argument “--feature_format Seq”, all the predicted sequence motifs filtered by convolution filters will be saved in the “output/encode_201/gc_match/TF_name/Seq/CNN/0/” directory.

**============================================================**

**9. TSPTFBS**

**Algorithm:** CNN

**Resource type:** Standalone

**See URL:** https://github.com/liulifenyf/TSPTFBS

| **Dependencies** | **Script** | **Bugs** | **Problem** | **Solution** |
| --- | --- | --- | --- | --- |
|  | /home/user/TSPTFBS/train.py | Lines 162-164 | File "Train.py", line 162  '''  ^  TabError: inconsistent use of tabs and spaces in indentation | Remove lines without  indentation |
| Python3,  Keras based architecture |  | Lines 159, 160 | FileNotFoundError: [Errno 2] No such file or directory: 'Example/ABF2.fasta' and ‘Example/neg.fasta’ | First of all move to Data directory and fetch ABF2.fasta file format from ABF2.bed file and then copy 'ABF2.fasta' and ‘neg.fasta’ to 'Example/' folder |
|  | /home/user/TSPTFBS/Predict.py | Line 73 to 78 | Input test prediction file format problem | Few lines are added to input test and label files |
|  |  |  | No performance metrics computation coding in program | A whole bunch of lines were added to calculate evaluation metrics |

**Command line:**

To execute the program, run the following command line in the Linux terminal:

- python3 Train.py TF_name
- python3 predict.py TF_name

**Output:**

The performance metrics text file output for model training (“TF_name_result.txt”) on test data is saved in the “model/” directory.

**============================================================**

**10. DNABERT**

**Algorithm:** Transformer

**Resource type:** Standalone

**See URL:** https://github.com/jerryji1993/DNABERT

| **Dependencies** | **Script** | **Bugs** | **Problem** | **Solution** |
| --- | --- | --- | --- | --- |
|  | /home/user/DNABERT/example/run_finetune.py | Line 37 | OSError: Model name 'PATH_TO_DNABERT_REPO/src/transformers/dnabert-config/bert-config-6/config.json' was not found in model name list. | path to "/home/user/DNABERT/src/transformers/dnabert-config/bert-config-6/config.json" is provided |
| Python3,  PyTorch based architecture |  | No | No performance metrics computation for coding in program | “example/compute_result.py” python script was edited to compute all performance metrics |
|  | /home/user/DNABERT/example/compute_result.py |  | Output of the prediction file was in “.npy” format and therefore actual labels were also converted to “.npy” format to compute all evaluation metrics | “example/compute_result.py” is edited to get evaluation metrics |

**Command line:**

To execute the program, run the following command line in the Linux terminal:

- cd example/
- python3 run_finetune.py --model_type dna --tokenizer_name=dna6 --model_name_or_path 6-new-12w-0 --task_name dnaprom --do_train --do_eval --data_dir TF_name --max_seq_length 100 --per_gpu_eval_batch_size=32 --per_gpu_train_batch_size=32 --learning_rate 2e-4 --num_train_epochs 2.0 –output_dir TF_name --evaluate_during_training --logging_steps 100 --save_steps 4000 --warmup_percent 0.1 --hidden_dropout_prob 0.1 --overwrite_output --weight_decay 0.01

**Note:** Here, the argument “--max_seq_length” is considered to specify the length of input sequences. It can be adjusted to 100 for sequences of 100 base pairs in length, 200 for sequences of 200 base pairs, and vice versa.

**============================================================**

**11. SeqConv**

**Algorithm:** CNN

**Resource type:** Standalone

**See URL:** https://github.com/shenwei19/SeqConv

| **Dependencies** | **Script** | **Bugs** | **Problem** | **Solution** |
| --- | --- | --- | --- | --- |
| Python3,  Keras based architecture | /home/user/SeqConv/src/prepare.py | No line number | can't open file 'prepare.py': [Errno 2] No such file or directory | Path to “src/prepare.py” provided |
|  |  | Line 10 | FileNotFoundError: [Errno 2] No such file or directory: 'fa/zm.fa' | - mkdir fa/ - cp zm.fa (Zea mays genome renaming and copy to fa/ directory) |
|  |  | Line 11 | Input file “in_file1” not found error | file1=open(in_file1).readlines() |
|  |  | Line 12 | FileNotFoundError: [Errno 2] No such file or directory: 'train/zm_tf_train.txt' | - The file named "zm_tf_train.txt" is in directory named "data". - Therefore, mkdir train/ and copy this file to train directory |
|  |  | Line 14 | TypeError: 'float' object cannot be interpreted as an integer | for i in range(len(file2)//2): |
|  | /home/user/SeqConv/src/generator.py |  | can't open file 'generator.py': [Errno 2] No such file or directory | src/generator.py |
|  |  | Line 6 | IndexError: list index out of range | input file path is not in proper way ,therefore path is given "train/zm_tf.narrowPeak" |
|  |  | Line 7 | FileNotFoundError: [Errno 2] No such file or directory: 'maize4.chrmsize' | file path is provided in the script  "data/maize4.chrmsize" |
|  |  | Line 18 | TypeError: 'dict_keys' object does not support indexing | chrm=random.choice(list(dic.keys())) -> to collect the keys fpr python3 |
|  |  | Lines 7, 25 | File “maize4.chrsize” and “train” directory not found error | Provide path to data/maize4.chrsize and make directory “train” |
|  | /home/user/SeqConv/src/SeqConv.py |  | python3: can't open file 'SeqConv.py': [Errno 2] No such file or directory | path to "src/SeqConv.py" directory |
|  |  | Line 12 | ImportError: cannot import name 'BatchNormalization' from 'keras.layers.normalization' (/home/user/.local/lib/python3.8/site-packages/keras/layers/normalization/__init__.py) | from tensorflow.keras.layers import BatchNormalization |
|  |  | Line 14 | ModuleNotFoundError: No module named 'keras.layers.advanced_activations' | from keras.layers import LeakyReLU |
|  |  |  | No performance metrics computation for coding in program | A whole bunch of lines were added to calculate evaluation metrics |

**Command line:**

To execute the program, run the following command line in the Linux terminal:

- python3 prepare.py TF_name_pos.bed GCF_000001735.3_TAIR10_genomic.fna TF_name_train.txt
- python3 generator.py TF_name_pos.bed
- python3 SeqConv.py TF_name

**Output:**

The performance metrics text file output for model training (“TF_name_result.txt”) on test data is saved in the “model/” directory.

**============================================================**

**12. MAResNet**

**Algorithm:** Top-down and bottom-up attention mechanisms with feed forward neural network (ResNets)

**Resource type:** Standalone

**See URL:** https://github.com/csbio-njust-edu/maresnet

| **Dependencies** | **Script** | **Bugs** | **Problem** | **Solution** |
| --- | --- | --- | --- | --- |
| Python3,  pytorch | /home/user/maresnet/train_on_cell_datasets.py | Line 35 | FileNotFoundError: [Errno 2] No such file or directory: 'Dataset/cell_dataset' | Need to download the global dataset (see the dataset section) to “Dataset/global_dataset” directory |
|  |  | Line 64 to 69 | performance metrics output stored in .pickle format | Format changed to “.csv” not to separately convert to “.pickle” to “.csv” format |
|  | /home/user/maresnet/train_test_api/test_api.py | Line 61 to 64 | performance metrics output stored in .pickle format | Format changed to “.csv” not to separately convert to “.pickle” to “.csv” format |
|  |  |  | No performance metrics computation for coding in program | A whole bunch of lines were added to calculate evaluation metrics |

**Command line:**

To execute the program and train the model on cell datasets, ensure you have three input files for each different TF data in the “example/TF_name” folder. Then, run the following command line in the Linux terminal:

- python3 train_on_cell_datasets.py

**Output:**

The models will be saved in the “checkpoint/” directory, while the prediction results and the log file will be saved in the “runs/” directory.

**============================================================**

**13. Wimtrap**

**Algorithm:** XGBoost

**Resource type:** Standalone

**See URL:** https://github.com/RiviereQuentin/Wimtrap

| **Dependencies** | **Script** | **Bugs** | **Problem** | **Solution** |
| --- | --- | --- | --- | --- |
| R package (R 4.0.4) | 1.R | N/A | N/A | N/A |

**Command:**

To enter the R environment, simply type “R” in the terminal. Then, you can run the following commands for your particular TF:

args <- commandArgs(trailingOnly = TRUE)

imported_genomic_data.seedlings <- importGenomicData(organism = "Arabidopsis thaliana",

genomic_data = c(

DHS = "example/DHS_athal_seedlings_normal.bed",

DGF = "example/DGF_athal_seedlings_7_days.bed",

CNS = "example/CNS_athal.bed"

))

**ABF2data.seedlings** <- getTFBSdata(pfm = "PFMs_athal.pfm", TFnames = "TF_name", organism = "Arabidopsis thaliana", genome_sequence = "GCF_000001735.3_TAIR10_genomic.fna", imported_genomic_data = imported_genomic_data.seedlings)

**ABF2model** <- buildTFBSmodel(TF_name_data.seedlings,

ChIPpeaks = c(TF_name = "TF_name.bed"),

model_assessment = TRUE)

**Output:**

The models and the performance metrics will be save in same parent directory.

============================================================

**14. PlantBind**

**Algorithm:** CNN+Bi-LSTM

**Resource type:** Standalone

**See URL:** https://github.com/wenkaiyan-kevin/PlantBind.git

| **Dependencies** | **Script** | **Bugs** | **Problem** | **Solution** |
| --- | --- | --- | --- | --- |
| Python3,  pytorch | /home/user/PlantBind/src/main.py |  | /home/user/anaconda3/bin/python: can't open file '/home/user/PlantBind/main.py': [Errno 2] No such file or directory | ‘/home/user/PlantBind/src/ main.py’ |
|  |  | Line 55 | RuntimeError: CUDA error: invalid device ordinal | python3 -m pip install --pre torch torchvision -f https://download.pytorch.org/whl/nightly/cu111/torch_nightly.html -U |
|  |  | Line 104 | RuntimeError(data_sampler = DistributedSampler(dataset)) | Comment out lines from 104-108 |
|  |  |  | RuntimeError: Unable to find a valid cuDNN algorithm to run convolution | Added “os.environ['TF_FORCE_GPU_ALLOW_GROWTH'] = 'true' ” to ‘/home/user/PlantBind/src/main.py’ |

**Command line:**

To execute the program, run the following command line in a Linux terminal:

- python3 -m torch.distributed.launch --nproc_per_node=2 src/main.py --inputs data_folder/ --length 101 --OHEM True --focal_loss True --batch_size 1024 --lr 0.01

Replace “TF_name” with the name of your TF to execute your program. Make sure you're in the correct directory where your program is located before running this command.

**Output:**

The following output files will be saved into the “output/” directory:

- “trained_model_101_seqs.pkl”
- “metrics_b.json”
- “metrics_m.json”
- “Gmean_threshold_b.txt”
- “Gmean_threshold_m.txt”
- “precision_recall_curve_Tfs_315.pdf”
- “roc_prc_curve_Tfs_315.pdf”

============================================================

**15. TSPTFBS 2.0**

**Algorithm:** DenseNet

**Resource type:** Standalone and Webserver

**See URL:** https://github.com/liulifenyf/TSPTFBS-2.0.git and <http://www.hzau-hulab.com/TSPTFBS/>

| **Dependencies** | **Script** | **Bugs** | **Problem** | **Solution** |
| --- | --- | --- | --- | --- |
| Python3,  keras | /home/user/PlantBind/src/densenet.py | Line 108 | FileNotFoundError: [Errno 2] No such file or directory: '/home/hlcheng/corn/sample/neg_sample.npy' | Modified and added many python functions for data input and train test the model. |

**Command line:**

To execute the program, run the following command line in a Linux terminal:

- python3.8 densenet.py TF_name

Replace “TF_name” with the name of your TF to execute your program. Make sure you're in the correct directory where your program is located before running this command.

**Output:**

The output model, named “TF_name_checkmodel.hdf5”, will be saved in the “TF_name_model/” directory. Additionally, the performance metric, named “TF_name_performance_metrics”, will be saved to the “TF_name/” directory.

============================================================

**16. DeepSTF**

**Algorithm:** CNN, transformer encoder, and Bi-LSTM

**Resource type:** Standalone

**See URL:** https://github.com/YuBinLab-QUST/DeepSTF.git

| **Dependencies** | **Script** | **Bugs** | **Problem** | **Solution** |
| --- | --- | --- | --- | --- |
| Python3,  Torch | /home/user/train.py | Error in “/home/user/DeepSTF/DeepSTF/models/DeepSTF.py” in line 44 | RuntimeError: Expected all tensors to be on the same device, but found at least two devices, cpu and cuda:0! (when checking argument for argument weight in method wrapper__native_layer_norm) | Adjusted all the functions written in the script “/home/user/DeepSTF/DeepSTF/models/DeepSTF.py” to the “/home/user/DeepSTF/DeepSTF/models/DeepSTF.py” to resolve all the errors related to “filenotfound” error to trained model saving. |

**Command line:**

To execute the program, run the following command line in a Linux terminal:

- python3 train.py TF_name

Replace “TF_name” with the name of your TF to execute your program. Make sure you're in the correct directory where your program is located before running this command.

**Output:**

The output model, named “TF_name_model.pth”, will be saved in the “model/” directory. The performance metric, named “TF_name_stats.txt”, will be saved to the “output/” directory.

**17. AgentBind**

**Algorithms:** CNN+BiLSTM (DanQ) and CNN (DeepSea)

**Resource type:** Standalone

**See URL:** <https://github.com/Pandaman-Ryan/AgentBind>

| **Dependencies** | **Script** | **Bugs** | **Problem** | **Solution** |
| --- | --- | --- | --- | --- |
| Python2.7,  Keras | AgentBind-Release-Version-DanQ/model | No description about data formation, how to run software to run on our own datsset | Data construction problem, how to run software to run on our own datsset | Provided a detailed pipeline to solve issues related to data format and model training and testing procedure |

**Command line:**

To execute the program, run the following command line in a Linux terminal:

- python2.7 model/train-transfer-learning.py --data_dir example/TF_name/training --valid_dir example/TF_name/validation --n_train_samples 8974 --n_valid_samples 2238 --train_dir TF_name_out --seq_size 1000 --checkpoint_dir TF_name_out --batch_size 128 --n_classes 2

Replace “TF_name” with the name of your TF to execute your program. Make sure you're in the correct directory where your program is located before running this command.

**NOTE:** Make sure to provide correct number of samples for ‘train_samples’ and ‘valid_samples’.

**Output:**

The output model, named “TF_name_model.pth”, will be saved in the “TF_name_out” directory. The performance metric, named “TF_name_stats.txt”, will be saved to the “TF_name_out” directory.

**18. BPNet**

**Algorithms:** CNN

**Resource type:** Standalone

**See URL:** https://github.com/kundajelab/bpnet

| **Dependencies** | **Script** | **Bugs** | **Problem** | **Solution** |
| --- | --- | --- | --- | --- |
| Python3.6,  conda or pip installation | bpnet | Compatibility with specific version python and approximately 275 packages of same specified versions. | Took too much time of approximately two and half day to install these libraries through conda environment | All the necessary steps and details of dependencies are provided in the GitHub ‘README.md’ file. |

**Command line:**

To execute the program, run the following command line in a Linux terminal:

- bpnet train --premade=bpnet9 examples/chip-nexus/dataspec.yml --override='seq_width=200;n_dil_layers=6' trained_model

**Output:**

The output model will be saved in the “trained_model/” directory.

**19. PTFSpot**

**Algorithm:** DenseNet **+** Transformer

**Resource type:** Standalone and webserever

**See URL:** https://github.com/SCBB-LAB/PTFSpot and https://scbb.ihbt.res.in/PTFSpot/index.php

| **Dependencies** | **Script** | **Bugs** | **Problem** | **Solution** |
| --- | --- | --- | --- | --- |
| Python3,  Torch | /home/user/train.py | N/A | N/A | N/A |

**Command line:**

Run following command line in Linux terminal to execute the program:

- python3 hyper_param.py file_for_tuning

**Output:**

The output model, “ptfspot.h5”, which is a hyperparameter-optimized trained model for the transformer, is saved in the “model/” directory. The performance metric, “TF_name_stats.txt” will be saved to “output/” directory.
