## Supplementary Material S4 for "Comprehensive analysis of computational approaches in plant transcription factors binding regions discovery"

**Supplementary Materials**

**List of dependencies for BPNet environment:**

1. argon2-cffi=20.1.0
2. async_generator=1.10
3. attrs=20.3.0
4. backcall=0.2.0
5. backports=1.0
6. backports.functools_lru_cache=1.6.1
7. bcftools=1.6
8. bcolz=1.2.1
9. bedtools=2.27.1
10. blas=2.20
11. bleach=1.4.2
12. blosc=1.21.0
13. bokeh=2.3.0
14. bzip2=1.0.8
15. ca-certificates=2020.12.5
16. certifi=2020.12.5
17. cffi=1.14.5
18. click=7.1.2
19. cloudpickle=1.6.0
20. contextvars=2.4
21. cudatoolkit=10.2.89
22. curl=7.71.1
23. cython=0.29.22
24. cytoolz=0.11.0
25. dask=2021.3.0
26. dask-core=2021.3.0
27. dataclasses=0.8
28. decorator=4.4.2
29. defusedxml=0.7.1
30. distributed=2021.3.0
31. entrypoints=0.3
32. fastparquet=0.4.0
33. freetype=2.10.4
34. fsspec=0.8.7
35. genomelake=0.1.4
36. h5py=2.9.0
37. hdf5=1.10.5
38. heapdict=1.0.1
39. html5lib=1.1
40. htslib=1.7
41. immutables=0.15
42. importlib-metadata=3.7.2
43. ipykernel=5.5.0
44. ipython=7.16.1
45. ipython_genutils=0.2.0
46. jedi=0.17.2
47. jinja2=2.11.3
48. jpeg=9d
49. jsonschema=3.2.0
50. jupyter_client=6.1.12
51. jupyter_core=4.7.1
52. jupyterlab_pygments=0.1.2
53. krb5=1.17.2
54. lcms2=2.12
55. ld_impl_linux-64=2.35.1
56. libblas=3.8.0
57. libcblas=3.8.0
58. libcurl=7.71.1
59. libedit=3.1.20191231
60. libffi=3.3
61. libgcc=7.2.0
62. libgcc-ng=9.3.0
63. libgfortran-ng=7.5.0
64. libgfortran4=7.5.0
65. liblapack=3.8.0
66. liblapacke=3.8.0
67. libllvm10=10.0.1
68. libpng=1.6.37
69. libsodium=1.0.18
70. libssh2=1.9.0
71. libstdcxx-ng=9.3.0
72. libtiff=4.2.0
73. libuv=1.41.0
74. libwebp-base=1.2.0
75. llvm-openmp=11.0.1
76. llvmlite=0.36.0
77. locket=0.2.0
78. lz4-c=1.9.3
79. lzo=2.10
80. markupsafe=1.1.1
81. mistune=0.8.4
82. mkl=2020.4
83. mock=4.0.3
84. msgpack-python=1.0.2
85. nb_conda=2.2.1
86. nb_conda_kernels=2.3.1
87. nbclient=0.5.3
88. nbconvert=6.0.7
89. nbformat=5.1.2
90. ncurses=6.2
91. nest-asyncio=1.4.3
92. ninja=1.10.2
93. notebook=6.2.0
94. numba=0.53.0
95. numexpr=2.7.3
96. numpy=1.18.5
97. olefile=0.46
98. openssl=1.1.1j
99. packaging=20.9
100. pandoc=2.12
101. pandocfilters=1.4.2
102. parso=0.7.1
103. partd=1.1.0
104. patsy=0.5.1
105. pexpect=4.8.0
106. pickleshare=0.7.5
107. pillow=8.1.2
108. pip=21.0.1
109. prometheus_client=0.9.0
110. prompt-toolkit=3.0.17
111. psutil=5.8.0
112. ptyprocess=0.7.0
113. pybedtools=0.7.10
114. pybigwig=0.3.17
115. pycparser=2.20
116. pygments=2.8.1
117. pyparsing=2.4.7
118. pyrsistent=0.17.3
119. pysam=0.14.0
120. pytables=3.6.1
121. python=3.6.13
122. python-dateutil=2.8.1
123. python-snappy=0.5.4
124. python_abi=3.6
125. pytorch=1.7.1
126. pytz=2021.1
127. pyyaml=5.4.1
128. pyzmq=22.0.3
129. readline=8.0
130. samtools=1.7
131. scipy=1.4.1
132. send2trash=1.5.0
133. setuptools=49.6.0
134. six=1.15.0
135. snappy=1.1.8
136. sortedcontainers=2.3.0
137. sqlite=3.34.0
138. statsmodels=0.11.1
139. tblib=1.6.0
140. terminado=0.9.2
141. testpath=0.4.4
142. thrift=0.11.0
143. tk=8.6.10
144. toolz=0.11.1
145. tornado=6.1
146. tqdm=4.46.1
147. traitlets=4.3.3
148. typing_extensions=3.7.4.3
149. wcwidth=0.2.5
150. webencodings=0.5.1
151. wheel=0.36.2
152. xz=5.2.5
153. yaml=0.2.5
154. zeromq=4.3.4
155. zict=2.0.0
156. zipp=3.4.1
157. zlib=1.2.11
158. zstd=1.4.9
159. absl-py==0.12.0
160. ansiwrap==0.8.4
161. appdirs==1.4.4
162. argcomplete==1.12.2
163. argh==0.26.2
164. arrow==1.0.3
165. astor==0.8.1
166. attr==0.3.1
167. binaryornot==0.4.4
168. black==20.8b1
169. bpnet==0.0.23
170. chardet==4.0.0
171. colorlog==4.7.2
172. comet-ml==3.5.0
173. concise==0.6.7
174. configobj==5.0.6
175. configparser==5.0.2
176. cookiecutter==1.7.2
177. coverage==5.5
178. cycler==0.10.0
179. deepexplain==0.1
180. deprecation==2.1.0
181. descartes==1.1.0
182. distlib==0.3.1
183. docker-pycreds==0.4.0
184. dulwich==0.20.20
185. everett==1.0.3
186. filelock==3.0.12
187. future==0.18.2
188. gast==0.4.0
189. gdown==3.12.2
190. gffutils==0.10.1
191. gin-config==0.4.0
192. gitdb==4.0.5
193. gitpython==3.1.14
194. google-pasta==0.2.0
195. grpcio==1.36.1
196. gtfparse==1.2.1
197. hyperopt==0.2.5
198. idna==2.10
199. imageio==2.9.0
200. importlib-resources==5.1.2
201. iniconfig==1.1.1
202. ipywidgets==7.6.3
203. jinja2-time==0.2.0
204. joblib==0.15.1
205. jupyterlab-widgets==1.0.0
206. keras==2.2.4
207. keras-applications==1.0.8
208. keras-preprocessing==1.1.2
209. kipoi==0.6.25
210. kipoi-conda==0.2.2
211. kipoi-utils==0.3.8
212. kipoiseq==0.3.4
213. kiwisolver==1.3.1
214. markdown==3.3.4
215. matplotlib==3.3.4
216. mizani==0.7.2
217. modisco==0.5.3.0
218. mypy-extensions==0.4.3
219. networkx==2.5
220. nvidia-ml-py3==7.352.0
221. palettable==3.3.0
222. pandas==1.1.5
223. papermill==2.3.3
224. pathspec==0.8.1
225. pathtools==0.1.2
226. plotnine==0.7.1
227. pluggy==0.13.1
228. poyo==0.5.0
229. promise==2.3
230. protobuf==3.15.6
231. py==1.10.0
232. pyfaidx==0.5.9.5
233. pysocks==1.7.1
234. pytest==6.2.2
235. pytest-cov==2.11.1
236. python-slugify==4.0.1
237. pywavelets==1.1.1
238. regex==2020.11.13
239. related==0.7.2
240. requests==2.25.1
241. requests-toolbelt==0.9.1
242. scikit-image==0.17.2
243. scikit-learn==0.21.3
244. seaborn==0.11.1
245. sentry-sdk==1.0.0
246. shapely==1.7.1
247. shortuuid==1.0.1
248. simplejson==3.17.2
249. smmap==3.0.5
250. subprocess32==3.5.4
251. tenacity==7.0.0
252. tensorboard==1.14.0
253. tensorflow==1.14.0
254. tensorflow-estimator==1.14.0
255. termcolor==1.1.0
256. text-unidecode==1.3
257. textwrap3==0.9.2
258. threadpoolctl==2.1.0
259. tifffile==2020.9.3
260. tinydb==4.4.0
261. toml==0.10.2
262. typed-ast==1.4.2
263. urllib3==1.26.3
264. vdom==0.6
265. virtualenv==20.4.2
266. wandb==0.10.22
267. websocket-client==0.58.0
268. werkzeug==1.0.1
269. widgetsnbextension==3.5.1
270. wrapt==1.12.1
271. wurlitzer==2.0.1
