## Supplementary Material S5 for "Comprehensive analysis of computational approaches in plant transcription factors binding regions discovery"

**Supplementary Materials**

**Tables**

**Table 1:** Hyper-parameters optimized for TFBSs/TFBRs identification software.

| **S.No.** | **Tools** | **Hyper-parameters** |
| --- | --- | --- |
| **1** | **DeepBind [1]** | Motif length= 24, number of motifs= 16, neural net hidden layer= 32, learning rate= 0.001, batch size= 128, learning steps= 20000, dropout= 0.50 |
| **2** | **gkmSVM [2]** | C = 1 |
| **3** | **LS-GKM [3]** | Kernel functions= wgkmrbf-kernel, C = 1, M = 50, H = 50.  Here, C= the regularization parameter, M = the initial value of the exponential decay function, H = the half-life parameter that is the distance required to fall to half of its initial value in the exponential decay function for kernel. |
| **4** | **KEGRU [4]** | Batch size= 200, kmer length=5, RNN unit= 70, Dropout= 0.5, activation function= softmax, optimizer= Adam, loss=Binary cross entropy |
| **5** | **k-mer grammar [5]** | *k*mer length = 8, C = 1.0, intercept scaling = 1, maximun iteration = 1000, penalty = l2, solver = newton-cg |
| **6** | **WSCNNLSTM [6]** | Convolutional neurons= 16, Convolutional kernel size= 1x24, Pooling size= 1x8, bi-LSTM neurons= 32, Dropout= 0.2, optimizer= Adadelta, Momentum in AdaDelta= 0.99, Learning rate= 1, Weight decay= 0.005, activation function= softmax, Epochs= 60, batchsize= 128 |
| **7** | **DeepRAM [7]** | Batch size= 128, learning steps= 5000, Optimizer= Adagrad, Dropout= 0.15, Activation function= ReLU |
| **8** | **DESSO [8]** | Convolutional filters= 16, 32, dropout = 0.2, Activation function= ReLU, Activation function (output layer)= Sigmoid, Optimizer= mini-batch gradient descent, Epochs= 30, batch size= 64 |
| **9** | **TSPTFBS [9]** | Activation function= ReLU, Hidden layers units= 100, 50, Dropout layer = 0.24, Activation function= Sigmoid, Optimizer= SGD, batch size= 64, epochs=50 |
| **10** | **DNABERT [10]** | *k*mer length= 6, learning rate= 0.0005, epochs= 5, batch size= 32, weight decay= 0.001, Optimizer= AdamW, adam epsilon= 1e-6, beta1= 0.9, beta2= 0.98 |
| **11** | **SeqConv [11]** | Convolutional filters= 16, Hidden layer units= 64 with activation function = ReLU, Activation function (output layer) = Sigmoid, loss= Mean squared error, Optimizer= Adam, batch size= 64, epochs= 1000 |
| **12** | **MAResNet [12]** | Optimizer= Adagrad, weight decay= 0.005, momentum= 0.9, dropout = 0.2, learning rate= 0.004, batch size= 256, loss= cross-entropy, activation function= softmax |
| **13** | **Wimtrap [13]** | Depth of the tree (maximum)= 6, learning rate = 0.3, number of iterations = 100, coefficient at the L2 regularization term of the cost function = 10, proportion of features used at each split selection = 1 and minimum instance in a leaf = 1, booster = tree, minimum loss reduction required to make a further partition on a leaf node of the tree = 0 |
| **14** | **PlantBind [14]** | Batch size=1024, learning rate = 0.01, Dropout= 0.2, Activation function= sigmoid |
| **15** | **TSPTFBS 2.0 [15]** | Two convolution layers with filters = 64, kernel size = 3 and activation function = ReLu, Four DenseBlocks (layer units = 6, 12, 24, 16 and three Transition Layers, Epochs= 40, learning rate= 0.001, Dropout = 0.2, activation function (output layer)= sigmoid, Optimizer= Adam, loss=binary crossentropy, batch_size=128 |
| **16** | **DeepSTF [16]** | Learning rate = 0.2, Optimizer = Adagrad, Dropout ratio = 0.2, Activation function= ReLU |
